# Analysis of geographic patterns of molecular, morphological, and bioclimatic variation to evaluate hypotheses of species boundaries in the South American montane genus *Escallonia* (Escalloniaceae)

**DOI:** 10.1101/009811

**Authors:** Felipe Zapata

## Abstract

*Escallonia* is a morphologically and ecologically diverse clade of shrubs and small trees widely distributed in three hotspots of plant diversity. Previous studies suggested that extant species may have radiated rapidly and/or recently resulting in complex patterns of molecular variation across this genus. This result is apparently mirrored in morphology because species also display complex and overlapping patterns of morphological variation. Taken together, these patterns call into question the identity of all species within *Escallonia*. To evaluate the currently proposed hypotheses of species boundaries, I used molecular, morphological, and bioclimatic a data for 35 species and assessed three species criteria: genealogical exclusivity, morphological gaps, and climatic niche differences. Interpreting these data in the context of species as segments of evolutionary lineages, I provide evidence that most species (ca. 70%) within *Escallonia* represent distinct species on independent evolutionary trajectories. Instead of rejecting the current hypotheses of species limit, I argue for taxonomic stability as it provides a useful framework for studies aiming to understand the mechanisms driving the origin and evolution of species in hotspots of biodiversity.

---

Understanding species boundaries provides valuable insight to help us elucidate the processes driving the origin and maintenance of biological diversity (Coyne and Orr, 2004). Since species boundaries emerge from the isolation and differentiation of populations (de Queiroz, 2005, 2007), analyzing the patterns of variation of molecular, phenotypic and ecological characters across the geographic range of a taxon is a robust approach to evaluate hypotheses of species boundaries (e.g., Sites and Crandall, 1997; Wiens and Penkrot, 2002; Wood and Nakazato, 2009; Cadena and Cuervo, 2010). However, there are few of these analyses for plant taxa, particularly from the hyperdiverse mountains of South America, where spatial heterogeneity may be an important driver of isolation and differentiation of species (Hughes and Eastwood, 2006).

Trees and shrubs of the genus *Escallonia* make an excellent case study for carrying out such analyses. These plants occur in a variety of habitats throughout the Andes and the mountains of southeastern Brazil, as well as in isolated mountain ranges like the Sierra de Córdoba (Argentina), Sierra Nevada de Santa Marta (Colombia), and Cordillera de Talamanca (Costa Rica). Most species have broad geographic ranges, with some species having populations separated by thousands of kilometers; a few narrowly distributed species span less than 200 kilometers. Several species seem to segregate according to habitat or elevation, nevertheless the geographic ranges of many species overlap completely or partially, such that individuals of one species can occur within the range of potential dispersal of gametes (seeds or pollen) of other species (i.e., species exhibit mosaic sympatry *sensu* Mallet, 2008). In all species, the fruit is a dry capsule that dehisces and releases the seeds, which fall out and are likely dispersed by wind or gravity. The only pollination study available in one species of *Escallonia* (Valdivia and Niemeyer, 2006) revealed that floral traits such as color and scent do not correlate with the expected specialist pollinator (Faegri and van der Pijl, 1979), and from circumstantial observations of several species in the field, the flowers of different species of *Escallonia* appear to be visited by the same diverse group of local insects that also visit unrelated plant genera. Some species are common locally, with approximately 30-40 plants per locality, while others are rare, few individuals being found in any one place (pers. obs.).

The geographic pattern of morphological variation within *Escallonia* is complex. All plants have a characteristic growth form with a distinctive long- and short-shoot construction (Bell, 2008), but the length of theses shoots varies extensively within and between species. There is substantial intra- and interspecific geographic variation in leaf size, shape and the density of serrations. Species have either single flowers, or inflorescences with tens to hundreds of flowers. The flowers show considerable intra- and interspecific geographic variation in the size and shape of sepals, petals and ovaries, the appearance of a nectary disk (flat or elevated), and in the pigmentation of petals, which range from greenish-white to deep red and several hues of pink in between. In some species, hairs and glands can occur on leaves, branches, or floral organs. This variation offers a unique opportunity for studying in detail the geographic patterns of variation in morphological, as well as molecular and ecological characters to evaluate hypotheses of species boundaries within *Escallonia*.

Phylogenies, multivariate statistics and geospatial analyses permit the analysis and description of patterns of variation within and among species with increasing statistical rigor. Such tools help systematists weigh the strength of empirical data to meet different operational species criteria to evaluate hypotheses of species boundaries (Sites and Marshall, 2003, 2004 and references therein; Wiens, 2007 and references therein; de Queiroz, 2005, 2007). Here, I provide the first comprehensive assessment of species boundaries in *Escallonia* examining the patterns of variation in molecular, morphological and ecological characters throughout the geographic range of this genus. In particular, I gauge the extent to which these data sets support the currently proposed hypotheses of species boundaries within *Escallonia* (Sleumer, 1968) by evaluating three operational species criteria.

## Criterion 1. Genealogical exclusivity

Evolutionary isolation and lineage divergence generate differential patterns of shared ancestry (Avise and Ball, 1990). I use the available phylogenetic hypotheses of *Escallonia* (Zapata, 2013) to examine the geographic patterns of intra- and interspecific molecular variation. This allows me to determine whether members of currently recognized species are more closely related to each other than they are to any individuals outside each species (i.e., genealogical exclusivity *sensu* Baum and Donoghue, 1995; Wiens and Penkrot, 2002).

## Criterion 2. Morphological discontinuity

Evolutionary isolation and lineage divergence generate discontinuities in the distribution of morphological variation (Futuyma, 1998; Rieseberg et al., 2006; Mallet, 2008). I use morphological characters to examine the geographic patterns of intra- and interspecific morphological variation, and evaluate whether currently recognized species are separated by morphological gaps (Wiens and Servedio, 2000; Zapata and Jiménez, 2012).

## Criterion 3. Niche differentiation

Evolutionary isolation and lineage divergence can be caused or reinforced by ecological divergence (Funk et al., 2006; Nosil and Crespi, 2006). I use bioclimatic information to describe broadly the realized ecological niche, and evaluate whether currently recognized species occur in significantly different environments, displaying differences in present day (environmental) selective regimes (Andersson, 1990; Rissler and Apodaca, 2007; Bond and Stockman, 2008; Leaché et al. 2009).

I further complement these analyses with estimates of genetic distance within and among species and relate this measurement to geographic distance as a crude measurement of reproductive isolation (Coyne and Orr 2004; Mallet 2008). Lastly, I examine differences in flowering time among species to get an initial idea of potential premating barriers.

## MATERIALS AND METHODS

### Study system

There are 39 species currently recognized in the genus *Escallonia* (Sleumer, 1968), which I interpreted as hypotheses of species boundaries. In this study, I analyzed 35 of these hypotheses, including only species for which molecular, morphological and climatic data were available. The four species I did not include are known from only few herbarium collections, and I failed to extract DNA from these collections or locate any populations of these species in the field.

All the specimens included in this study were assigned to each of the 35 hypothesized species after detailed comparative studies of morphology and using the dichotomous key provided by Sleumer (1968). In many instances, I was able to use the same specimens that Sleumer observed in his study of *Escallonia*. All specimens used for molecular analyses were also included in the morphological and ecological analyses, except for both specimens of *E. ledifolia* and *E. petrophila,* one specimen of *E. illinita, E. leucantha*, *E. myrtoidea*, *E. pulverulenta*, *E.revoluta,* and *E. rosea,* and two specimens of *E. megapotamica*, none of which had flowers and so could not be measured for all the phenotypic characters studied here (see below). However, habit, vegetative characters, geographic locality and the available comparative material from herbarium collections allowed me to reliably assign these specimens to species.

### Molecular data

To examine the geographic pattern of molecular variation, I used two haplotype phylogenies based on Bayesian and maximum likelihood analyses of the first intron of a MYC-like gene and the third intron of the NIA gene (for details, see Zapata, 2013). The MYC matrix consisted of 887 bp for 89 individuals and 102 terminals, while the NIA matrix consisted of 843 bp for 88 individuals and 107 terminals; several individuals were heterozygotes for both loci. In total, between one and seven individuals were sampled for all species (Table 1). These samples covered the whole geographic range of *Escallonia*; for species with broad geographic distributions, samples from well-spaced localities from across their geographic ranges were included whenever possible (Zapata, 2013). Since nothing is known about the position of MYC and NIA in the genome of *Escallonia*, I interpreted each haplotype tree as an independent source of evidence of lineage divergence (de Queiroz, 2007).

**TABLE 1.**
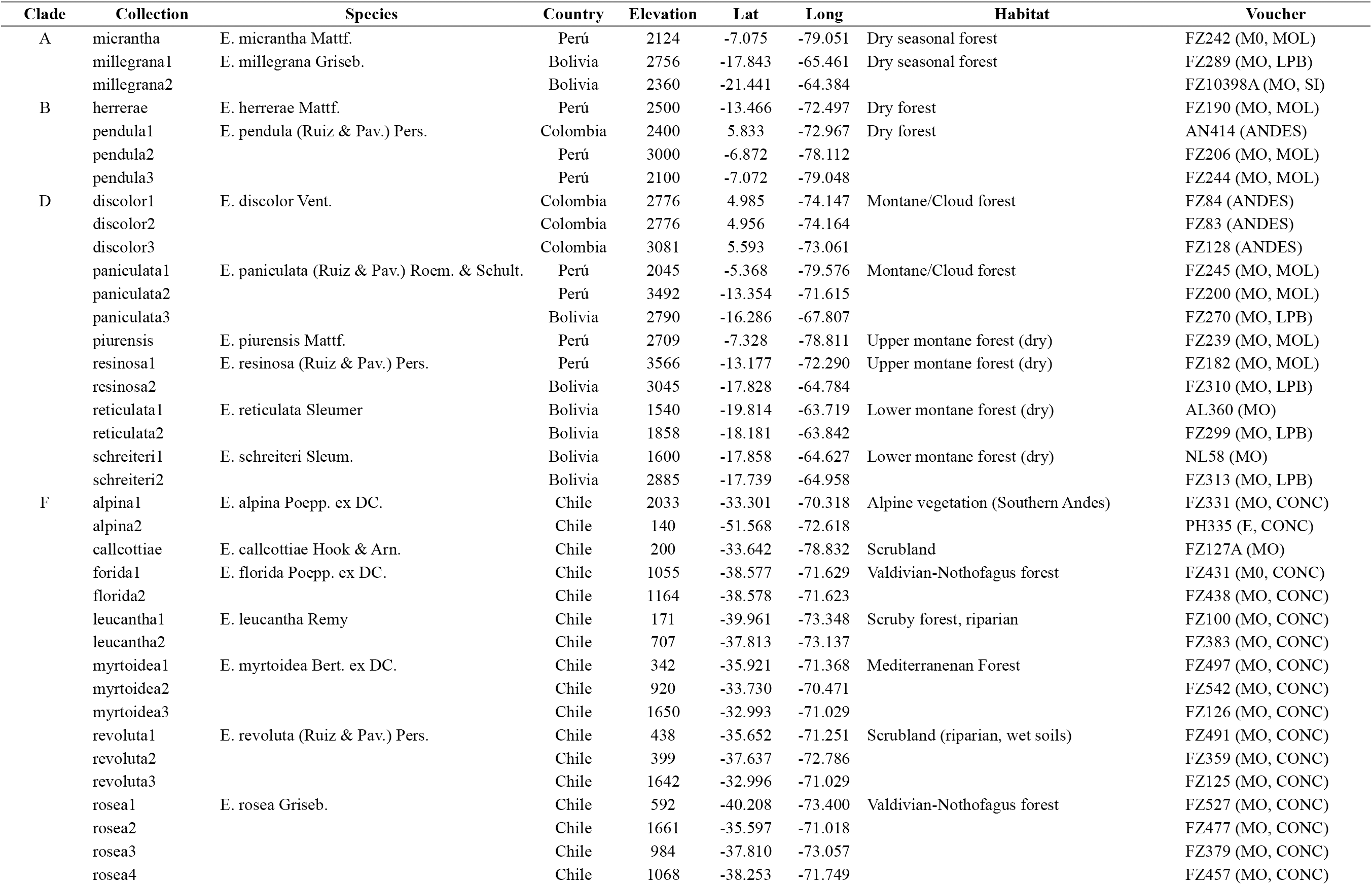
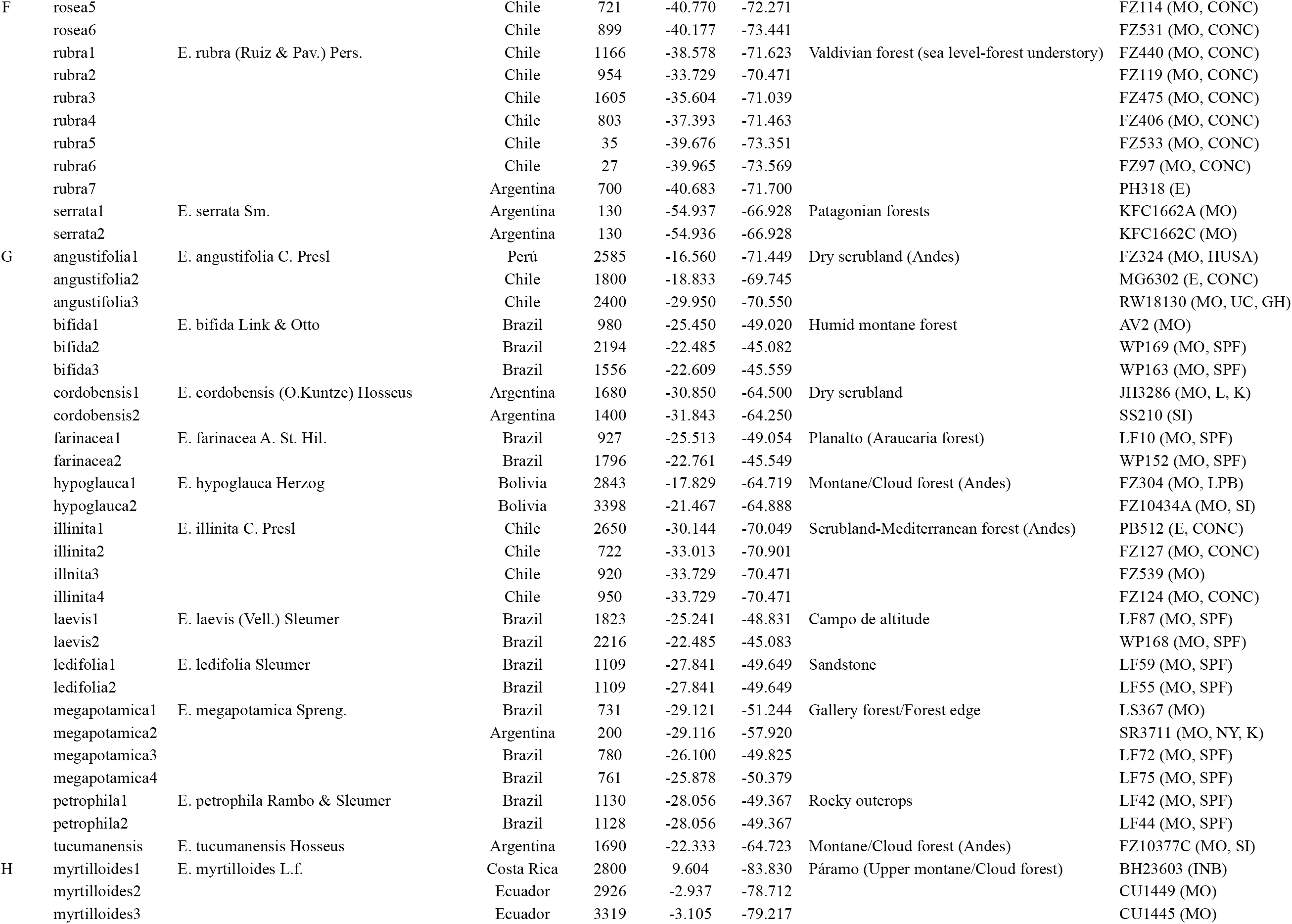
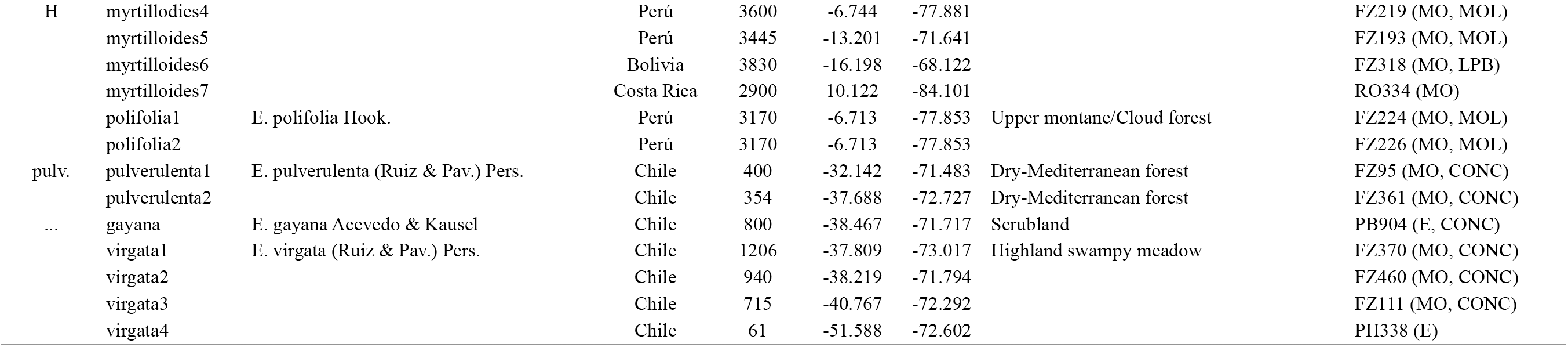
Specimens included in molecular study. Lat. and Long. in decimal degrees. Voucher: Collector Initials+Collection Number (herbaria where collections are deposited; for herbaria name see Thiers, [continuously updated]). For clade names see Fig. 1.

I implemented the tree-based method of Wiens and Penkrot (2002) to weigh the strength of molecular data in supporting the hypothesis that the members of the currently recognized species within *Escallonia* are more closely related to each other than they are to any individuals outside each putative species (see Baum and Donoghue, 1995). Briefly, the method of Wiens and Penkrot (2002) uses a haplotype phylogeny derived from multiple individuals and populations of a hypothesized species (the focal species) and one or more closely related species, to evaluate the concordance between the geographic origin of the haplotypes, and their phylogenetic relationships. From this, they infer the taxonomic status of the focal species (see Fig. 1 in Wiens and Penkrot, 2002). Focal species can be genealogically exclusive or non-exclusive, and depending on the geographic concordance of clades at shallow and deep nodes (i.e., ‘basal lineages’ *sensu* Wiens and Penkrot, 2002), the method presents a decision tree to allow one to infer whether there is enough evidence to suggest the focal species is a single species. This species can be: i) a monophyletic species, that is an exclusive lineage concordant with geography (see Fig. 1b in Wiens and Penkrot, 2002); or ii) a paraphyletic species (“plesiospecies” *sensu* Olmstead, 1995), that is a non-exclusive lineage concordant with geography (see Fig. 1d in Wiens and Penkrot, 2002). In other cases, the focal species is not a single species, rather it can be: iii) a conspecific lineage, that is a broader group including several species perhaps linked through gene flow (see Fig. 1f in Wiens and Penkrot, 2002); or iv) multiple distinct species, that is the focal species includes more than one species (see Fig. 1a, c, e in Wiens and Penkrot, 2002).

**Fig. 1.**
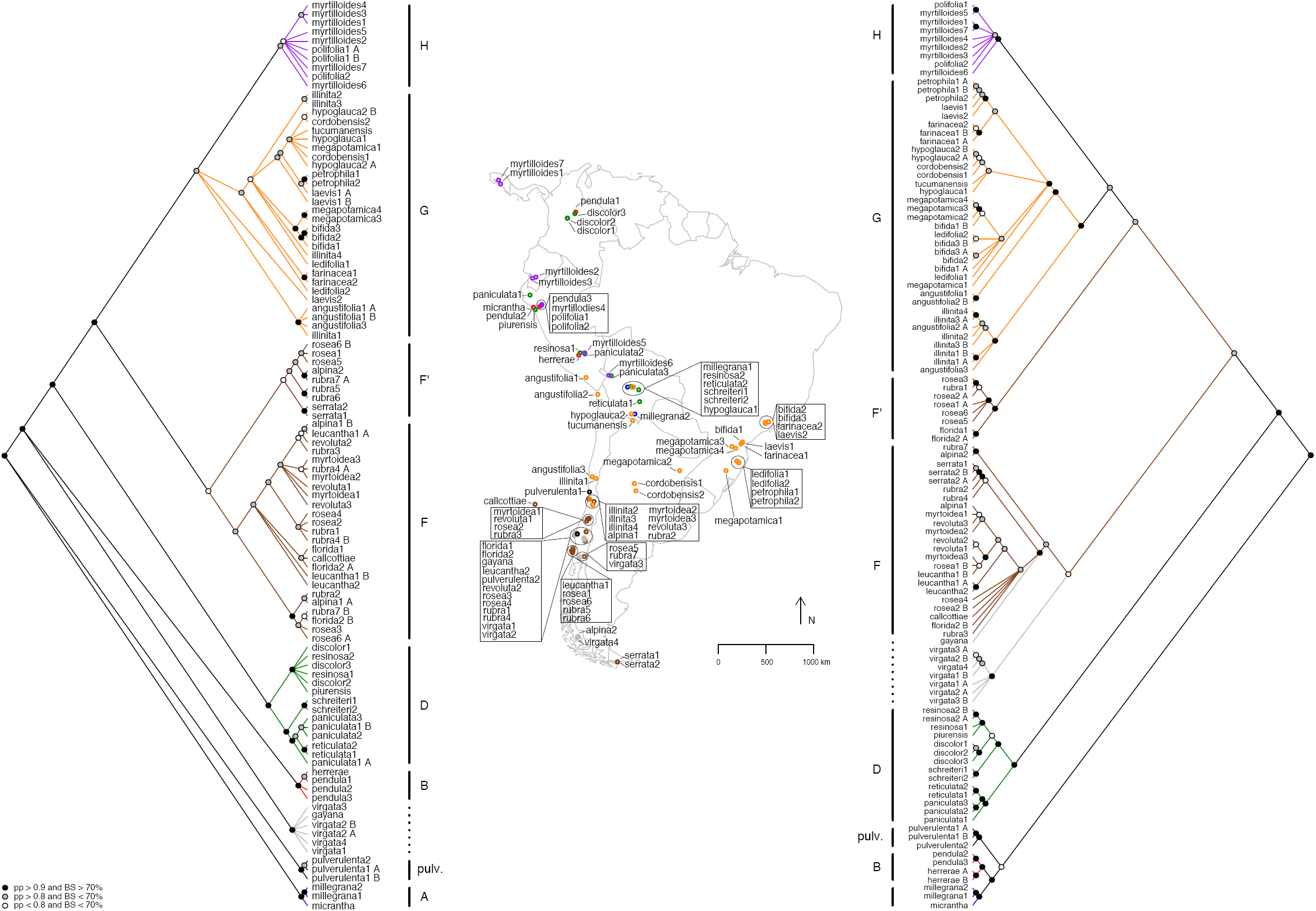
Haplotype trees for species of *Escallonia* based on Bayesian and maximum likelihood phylogenetic inference for MYC (left) and NIA (right) loci (Zapata, 2013), and map of sampling localities. In trees, node color indicates Bayesian posterior probability (pp) and maximum likelihood bootstrap (BS) (2000 replicates): black: pp > 0.9 and BS > 70%; gray: pp > 0.8 and BS < 70%; white: pp < 0.8 and BS < 70%. The figure is modified from Zapata (2010), and only topology is depicted (for phylogram, see Zapata, 2013). Each sampled haplotype is given the specific epithet and a number, this latter in no particular order. For heterozygote individuals, A/ B inidicates each allele. Letters in front of bars indicate the names of groups of species as indicated by Zapata (2010). Each group of species is color-coded: clade A: blue; clade B: red; clade D: green; clade F: brown; clade G: orange; clade H: purple; pulv. (*E. pulverulenta*): black; *E. virgata + E. gayana*: gray. *E. virgata + E. gayana* are indicated with dotted line because these species do not form a clade in both analyses. Clade F’ indicates a paralogous copy for each locus, found only for species within clade F (for details, see text and Zapata, 2013). Map of South America with sampling localities color coded with respect to colors of clades in the haplotype trees. Both trees show phylogenetic geographic structure (see text for details).

This method assumes exhaustive sampling of individuals, populations and species, and recommends the use of quantitative approaches (e.g., Templeton, 2001, 2010; but see Beaumont et al., 2010) to assess objectively the general concordance between geography and phylogeny. The sampling in the study of Zapata (2013) was too coarse and non-exhaustive at the population level to implement here appropriate statistical methods to estimate such concordance (Knowles, 2009). Nonetheless, both haplotype phylogenies revealed a remarkable level of phylogenetic geographic structure (Zapata, 2013; see below), which provides a useful framework to guide the assessment of species boundaries that I complement with the analyses of the patterns of variation in the other types of data studied here. Furthermore, for species with broad geographic ranges, multiple samples from well-spaced localities allowed some rigor in assessing the genealogical exclusivity of species. Therefore the study of Zapata (2013) provided a reasonable first step to evaluate hypotheses of species boundaries within *Escallonia* using molecular data at a continental scale.

Using PAUP* 4.0 (Swofford, 2002), I calculated the genetic distance within and among species using the TrN+I+Γ and the GTR + I+Γ models of sequence evolution for MYC and NIA, respectively. These models were selected using the model testing procedure implemented in DT-ModSel (Minin et al., 2003), and used to infer the haplotype trees (Zapata, 2013). To determine whether geography could explain the overall pattern of genetic variation, I assessed the relationship between geographic and genetic distance using a simple Mantel test with 999 permutations using R 3.0.2 (R Core Team, 2013) and the library ade4 (Dray and Dufour, 2007).

### Morphological data

To examine the geographic pattern of morphological variation, I sampled a total of 679 specimens from field and herbarium collections to represent the overall spectrum of morphological variation and the geographic range of each of the 35 hypothesized species (minimum 2; maximum 61 specimens per species; Appendix 1). I estimated the pattern of phenotypic variation among these species in 27 quantitative (6 vegetative; 21 reproductive) and 13 qualitative (1 habit; 6 vegetative; 6 floral) characters (Table 2, Appendix 2). These 40 characters were selected after careful study of specimens and a thorough literature review (Kausel, 1953; Sleumer, 1968). I used only mature leaves and flowers in all specimens to measure each character. Quantitative vegetative characters were measured using a standard metric ruler on dried specimens. Quantitative floral characters were measured using a digital caliper (Digimatic CD-6” CS, Mitutoyo Japan) on flowers that were rehydrated and examined on a stereoscopic dissecting microscope (SMZ645, Nikon USA). All quantitative measurements were recorded from three different structures for each specimen whenever possible, and then averaged to generate character measurements for each specimen. Habit and petal color were gathered from herbarium labels and field observations. All other qualitative characters were measured using a stereoscopic dissecting microscope on a single organ.

**TABLE 2.** Morphological characters measured for all specimens, and used in morphometric analyses to assess morphological gaps.

| Character | Unit | Abbreviation |
| --- | --- | --- |
| QUANTITATIVE |  |  |
| Petiole Length | mm | PETLEN |
| Lamina length | mm | LAMLEN |
| Lamina width at widest point | mm | LAMWID |
| Lamina length to widest point | mm | LAMLENWID |
| Lamina length:Lamina width | ratio | LAMWID:LAMLEN |
| Lamina length:Lamina length to widest point | ratio | LAMLENWID:LAMLEN |
| Length of reproductive shoot | mm | INFLLEN |
| Length of 2ary inflorescence axis | mm | INFBRANLEN |
| Pedicel length | mm | PEDLEN |
| Pedicel width | mm | PEDWID |
| Ovary length | mm | OVALEN |
| Ovary width | mm | OVAWID |
| Ovary length:Ovary width | ratio | OVAWID:OVALEN |
| Calyx tube length | mm | CALYTUBLEN |
| Calyx lobe length | mm | CALLOBLEN |
| Calyx lobe width | mm | CALLOBWID |
| Calyx tube length:Calyx lobe length | ratio | CALYTUBLEN:CALLOBLEN |
| Calyx lobe length:Calyx lobe width | ratio | CALLOBWID:CALLOBLEN |
| Petal length | mm | PETLEN |
| Petal length to widest point | mm | PETLENWID |
| Petal length to point before spreading | mm | PETLENSPR |
| Petal basal width | mm | PETBASWID |
| Petal width at widest point | mm | PETWID |
| Petal length:Petal width at widest point | ratio | PETWID:PETLEN |
| Petal length:Petal length to widest point | ratio | PETLENWID:PETLEN |
| Petal length:Petal length to point before spreading | ratio | PETLENSPR:PETLEN |
| Petal width at widest point:Petal basal width | ratio | PETBASWID:PETWID |
| QUALITATIVE |  |  |
| Habit | QL: 0(Tree), 1(Shrub), 2(Subahrub) | HABIT |
| Glands on adaxial surface of lamina | QL: 0(Absent), 1(Present) | ADLAMGLA |
| Indumentum on adaxial surface of lamina | QL | ADLAMPUB |
| Glands on abaxial surface of lamina | QL | ABLAMGLA |
| Indumentum on abaxial surface of lamina | QL | ABLAMPUB |
| Margin-Teeth | QL | LAMMARG |
| Margin Appearance | QL: 0(Flat), 1(Revolute) | MARGAPP |
| Glands on pedicel | QL | PEDGLAND |
| Indumentum on pedicel | QL | PEDPUB |
| Glands on ovary | QL | OVAGLAN |
| Indumentum on ovary | QL | OVAPUB |
| Style-Disk | QL: 0(Flat), 1(Elevated) | STYDISK |
| Petal color | QL: 1(Red), 2W(hite), 3(Pink), 4(Green) | FLOCOLOR |

I used the method proposed by Zapata and Jiménez (2012) to weigh the strength of the quantitative characters to support the hypothesis that currently recognized species within *Escallonia* correspond to lineages separated by morphological gaps (see Futuyma, 1998; Rieseberg, 2006; Mallet, 2008). In this method, a gap in multivariate morphological space between a pair of hypothesized species is inferred when i) the mixture of the two species is bimodal, and ii) intermediate phenotypes in the mixture occur at a frequency below a predetermined cutoff. To assess bimodality, the method uses the samples of the two hypothesized species to estimate the *pdf* of the mixture of two normal distributions describing the morphological variation of each hypothesized species, and plots the values of this function along the ridgeline manifold, a curve guaranteed to include all the critical points (maxima, minima and saddles) of the estimated *pdf* (for details, see Ray and Lindsay, 2005; Zapata and Jiménez, 2012). If this plot reveals bimodality, the frequency of the intermediate phenotypes between the two hypothesized species is estimated. The method calculates a series of ellipsoids defining tolerance regions–the regions covering a proportion β of a population with statistical confidence γ (Krishnamoorthy and Mathew, 1999; Zapata and Jiménez, 2012)–for each hypothesized species that share a point along the ridgeline manifold, and calculates the proportions of the distributions describing each hypothesized species covered by each of these ellipsoids. Since each pair of these ellipsoids also shares a tangent line at the ridgeline manifold that splits the bivariate morphological space occupied by the species, here I present a modification of the original method proposed by Zapata and Jiménez (2012), and I also calculated the proportions of the areas of the distributions describing each hypothesized species that lie away from each tangent line. A plot of the proportions (for both, tolerance regions and areas) for several points along the ridgeline manifold reveals whether the distributions describing the morphological variation of each hypothesized species overlap for a proportion equal to or less than a predetermined frequency cutoff. If so, this result would support the hypothesis that the samples of the two hypothesized species are separated by a morphological gap, i.e., the intermediate phenotypes occur at a frequency below the predetermined cutoff, and thus it can be inferred that they represent two distinct species (Zapata and Jiménez, 2012). To account for the possibility that the morphological gap simply reflects geographic differentiation within a single species (see de Queiroz and Good, 1997; de Queiroz, 2007), the geographic coordinates of the specimens included in the morphometric analysis are used to derive a series of orthogonal vectors (i.e., spatial eigenvectors) to model morphological variation at different spatial scales assuming no species boundary (for details, see Borcard and Legendre, 2002; Dray et al., 2006; Griffith and Peres-Neto, 2006; Zapata and Jiménez, 2012). Multivariate multiple regression (Rao, 1964) is used to evaluate whether this model is significantly worse than a contrasting model requiring a species boundary, in which case the hypothesis that the morphological gap reflects true evolutionary isolation (i.e., two distinct species) is considerably strengthened (for details, see Zapata and Jiménez, 2012).

In this study, I implemented this method in the following way. Based on the results of the molecular analyses (see Results), I used the measurements of the continuous characters to derive orthogonal axes using Principal Component Analysis (PCA) calculated on a correlation matrix to define a morphological space for the species within well supported and concordant clades. In particular, I always used the space defined by PC1 and PC2 to estimate the *pdf* of the mixture of the distributions describing the pattern of morphological variation of each pair of species, and I plotted the values of the *pdf* along the ridgeline manifold (hereafter referred to as “elevation plot”, see Ray and Lindsay, 2005) to inspect visually for bimodality. When the *pdf* was not bimodal, I stopped the analyses because the first necessary condition to support the hypothesis of a morphological gap separating a pair of species failed (Zapata and Jiménez, 2012). If the *pdf* was bimodal, I plotted the proportions of the areas of the distributions describing the pattern of morphological variation of each species along the ridgeline manifold (hereafter referred to as “proportion plot”), and I inferred there was enough evidence to suggest a morphological gap when proportions ≤ 0.1 of these distributions overlapped (i.e, frequency cutoff = 0.1); I also plotted the proportions of the distributions covered by the ellipsoids defining tolerance regions for comparative purposes. All ellipsoid tolerance regions were calculated using statistical confidence γ = 0.95.

When there was more than one pair of hypothesized species within a clade, I used the morphological space derived using the measurements of the samples of all the species in the clade, and I estimated morphological gaps between all pairs of species in that morphological space. If the samples of a hypothesized species appeared non-overlapping and the samples of other species appeared overlapping, I estimated the morphological gaps between the non-overlapping species and all other species in that PCA morphological space, and subsequently I derived a new morphological space using only the samples of the species that overlapped initially. In this new morphological space, I evaluated morphological gaps for the species that overlapped in the original morphological space. I repeated this approach iteratively so long as samples of a species appeared non-overlapping and samples of the other species appeared overlapping in any morphological space. This approach allowed me to estimate morphological gaps on the axes of variation relevant for each group of species.

When I inferred a morphological gap between a pair of species, I evaluated whether there was enough evidence to suggest such a gap represented a species limit rather than geographic differentiation within a single species. I used the geographic coordinates of all the specimens included in each pairwise comparison to derive a series of orthogonal vectors (i.e., spatial eigenvectors) to model morphological variation at different spatial scales assuming no species boundary. This model was contrasted against a model that required the hypothesized species boundary represented by an indicator matrix [0,1] and the interactions between this matrix and the spatial eigenvectors (for details, see Zapata and Jiménez, 2012). I assessed statistical significance of each model using RDA with 9999 permutations. All analyses were conducted in R 3.0.2 (R Core Team, 2013) using the same packages, scripts and settings of Zapata and Jiménez (2012).

When I found no evidence of a gap in quantitative continuous characters separating a pair of species, I looked for gaps in qualitative characters. For these analyses, I compared the frequencies of the character states between such pairs of species to search for potentially non-overlapping characters. Characters that were invariant for alternative states within a putative species were considered non-overlapping and thus potentially useful to support the hypothesis of a species boundary. I used the method of Wiens and Servedio (2000) to weigh the strength of these characters to support the hypothesis that species were separated by a morphological gap in qualitative characters. Specifically, I used the equation 3 of Wiens and Servedio (2000) to calculate a *P*-value to evaluate whether in a sample of apparently invariant characters, at least one of these characters was not polymorphic above a selected frequency cutoff. For this test, I used a frequency cutoff 1-*β* = 0.1 and statistical confidence γ = 0.95. Thus, failing this test would mean that there is a > 0.05 probability that all the apparently invariant characters for a given species are actually polymorphic, with a frequency of the alternative character state above 0.1 (for details, see Wiens and Servedio, 2000).

### Bioclimatic data

To examine the pattern of ecological variation, I obtained point locality data from herbarium labels or online gazetteers (Guralnick et al. 2006) for the same specimens I analyzed for morphological variation, and I checked these data with high resolution maps to ensure locality precision. Using ArcView (ESRI), I mapped these specimens and extracted the values of 19 bioclimatic variables (Table 3) at each point locality from climatic layers with resolution of a square kilometer obtained from WorldClim v1.4 (Hijmans et al., 2005). These variables reflect annual trends in temperature and precipitation, seasonality, and extreme environmental conditions (e.g., temperature of the coldest month), and are thus likely to represent general limiting factors for plant physiology, survival, and growth (Taiz and Zeiger, 2006). Based on the results of the molecular analyses (see Results), I used the variation in these bioclimatic variables along with elevation to derive orthogonal axes using Principal Component Analysis (PCA) calculated on a correlation matrix as a reasonable approximation to characterize the realized ecological niche, i.e., the present-day selective regime (see Stockman and Bond, 2007; Bond and Stockman 2008) of each species within well supported and concordant clades. I used MANOVA (multivariate analyses of variance) with PC1 and PC2 scores as dependent variables and species as the fixed factors (see Graham et al., 2004; Rissler and Apodaca, 2007; Stockman and Bond, 2007; Bond and Stockman, 2008; Leaché et al., 2009) to weigh how strongly these data supported the hypothesis that currently recognized species within *Escallonia* differed in their realized ecological niche (see Andersson, 1990). When a MANOVA was significant in a clade with more than one pair of species, I ran Tukey’s Honest Significant Difference tests (Tukey’s HSD) to determine which species were different.

**TABLE 3.** Bioclimatic variables extracted from point localities of all specimens included in morphometric analyses (see Appendix 1). These variables were used in analyses of ecological differentiation among species.

| Abbreviation | Original Variable |
| --- | --- |
| BIO1 | Annual Mean Temperature |
| BIO2 | Mean Diurnal Range (Mean of monthly (max temp - min temp)) |
| BIO3 | Isothermality (P2/P7) (* 100) |
| BIO4 | Temperature Seasonality (standard deviation *100) |
| BIO5 | Max Temperature of Warmest Month |
| BIO6 | Min Temperature of Coldest Month |
| BIO7 | Temperature Annual Range (P5-P6) |
| BIO8 | Mean Temperature of Wettest Quarter |
| BIO9 | Mean Temperature of Driest Quarter |
| BIO10 | Mean Temperature of Warmest Quarter |
| BIO11 | Mean Temperature of Coldest Quarter |
| BIO12 | Annual Precipitation |
| BIO13 | Precipitation of Wettest Month |
| BIO14 | Precipitation of Driest Month |
| BIO15 | Precipitation Seasonality (Coefficient of Variation) |
| BIO16 | Precipitation of Wettest Quarter |
| BIO17 | Precipitation of Driest Quarter |
| BIO18 | Precipitation of Warmest Quarter |
| BIO19 | Precipitation of Coldest Quarter |

### Phenological data

Because only specimens with flowers were used for morphological and ecological analyses, I used the collection date of each specimen taken from herbarium labels to examine the overlap in the range of flowering time among species within clades. In the absence of quantitative information on phenological variation within *Escallonia*, I used these data as a proxy to assess potential premating isolating barriers.

## RESULTS

Independent Bayesian and maximum likelihood phylogenetic analyses of the nuclear loci MYC and NIA recovered at least eight groups of species in common (hereafter referred to as clades A, B, D, F, G, H, and groups *E. pulverulenta* and *E. virgata* + *E. gayana;* Fig. 1; see also; for details, see Zapata, 2013). For some species within clade F, an apparent paralog of MYC and NIA was also recovered. Haplotypes of this paralog formed a clade in both gene trees (hereafter referred to as clade F’). Although the phylogenetic position of clade F’ was not consistent between gene trees (Fig. 1), all the haplotypes within this clade were always more closely related to each other than to haplotypes in any other clade (for details, see Zapata, 2013). Therefore, I used clade F’ as an extra locus to assess species limits for the species within this clade independently of their position within clade F (see below). All clades were markedly restricted to geographic regions, except clade G; this was mainly restricted to southeastern Brazil and northeastern Argentina, but included some species in the Andes and the Sierra de Córdoba in central Argentina (Fig. 1). Phylogenetic analyses using either one or several individuals per species, consistently recovered all eight groups of species, and these groups always received very high to moderately high Bayesian posterior probabilities and bootstrap support. The exceptions were clade F, mainly because of the occurrence of F’ within it, and the group *E. virgata* + *E. gayana* (see below). Relationships among the eight groups were broadly concordant between gene trees, except for clade B and groups *E. pulverulenta* and *E. virgata* + *E. gayana*; nevertheless, all but one of these conflicting phylogenetic positions received low statistical support. Conversely, phylogenetic relationships within most groups were largely discordant between gene trees at all nodes (Fig. 1; for details, see Zapata, 2013).

Both loci showed relatively low levels of overall sequence divergence. Corrected genetic distances ranged from 0 to 0.06 in the MYC matrix (Fig. 2) and from 0 to 0.15 in the NIA matrix (Fig. 3). A simple Mantel test (Fig. 4) showed that geographic distance was strongly correlated with genetic distance for NIA (*r* = 0.2, *P* = 0.002) and weakly correlated, but not statistically significant, for MYC (*r* = 0.09, *P* = 0.064). This is consistent with the phylogenetic geographic structure revealed by the gene trees (Fig 1); the low correlation in MYC was likely due to the low level of sequence divergence shown by this locus (Fig. 2). Consistent with the Mantel test results, genetic distances were generally lower within groups than among groups for both loci. The average sequence divergence within groups was 0.01 and 0.02, and among groups was 0.04 and 0.08 for MYC and NIA, respectively. This suggests that species within groups are less divergent than species among groups. Furthermore, genetic distances were generally lower within species than among species at different geographic distances (Fig. 4). A closer examination of the relationship between geographic and genetic distances revealed that when different groups of species co-occurred in mosaic sympatry (*sensu* Mallet, 2008) these groups were genetically distant, for example clades A and B, or the clade F and groups *E. pulverulenta* and *E. virgata* + *E. gayana,* (see Figs. 2-4). This result, along with the marked phylogenetic geographic concordance of groups, the consistent composition of species within groups in both phylogenetic analyses, and the differences in genetic distance within and among groups, suggests that these groups are likely evolutionarily isolated. Therefore, I called these groups “basal lineages” (see Wiens and Penkrot, 2002), within which I analyzed species limits. In particular, I examined the patterns of variation in molecular, morphological and ecological data for all the species within each group, and did not use these data to compare species from different groups (except for *E. pulverulenta* and *E. virgata + E. gayana*, see below).

**Fig. 2.**
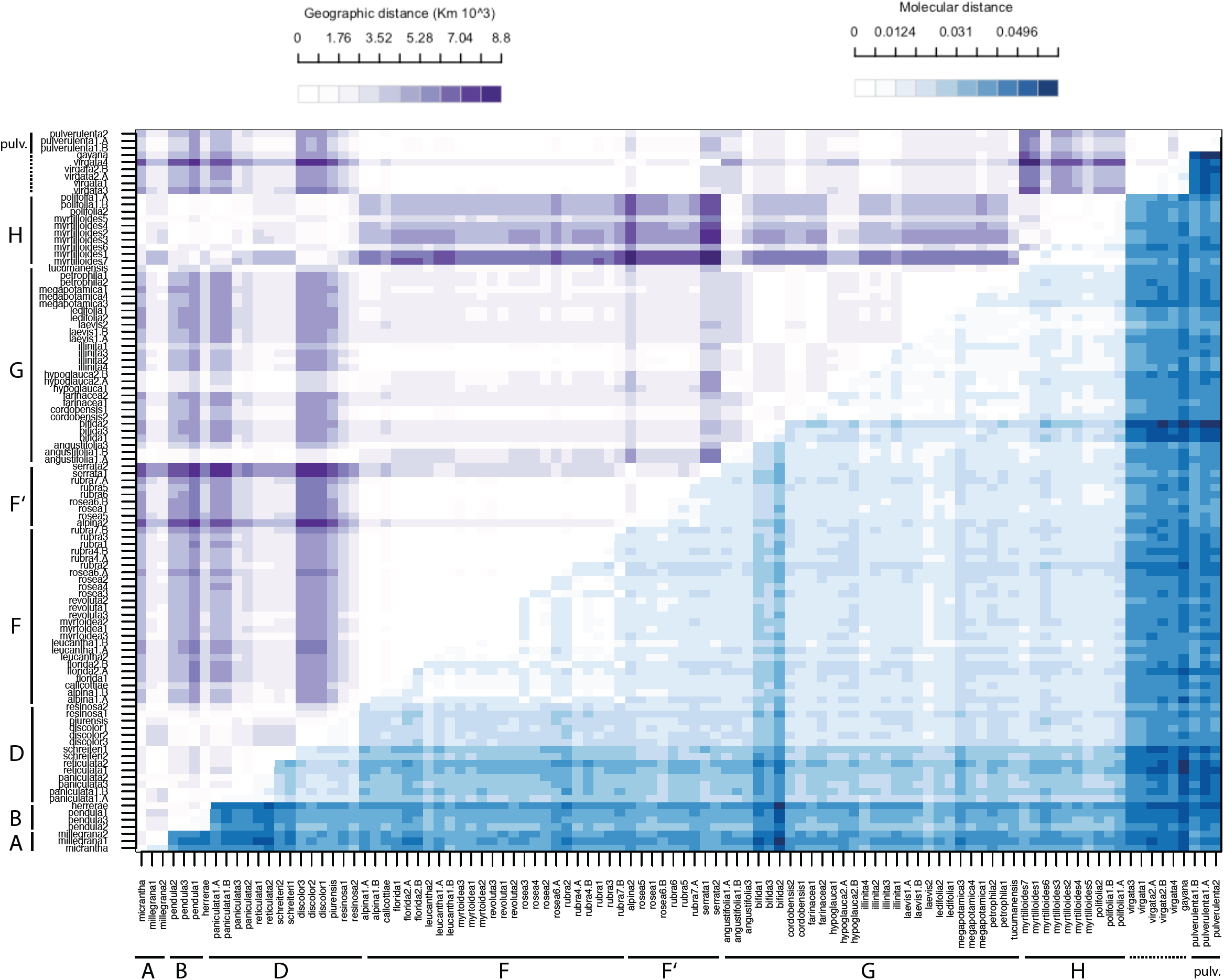
Heatmap showing relationship between geographic and genetic distance for the MYC locus. Dark shade: higher values; pale shade: lower values. See scale for variation. For clades names, see Fig. 1.

**Fig. 3.**
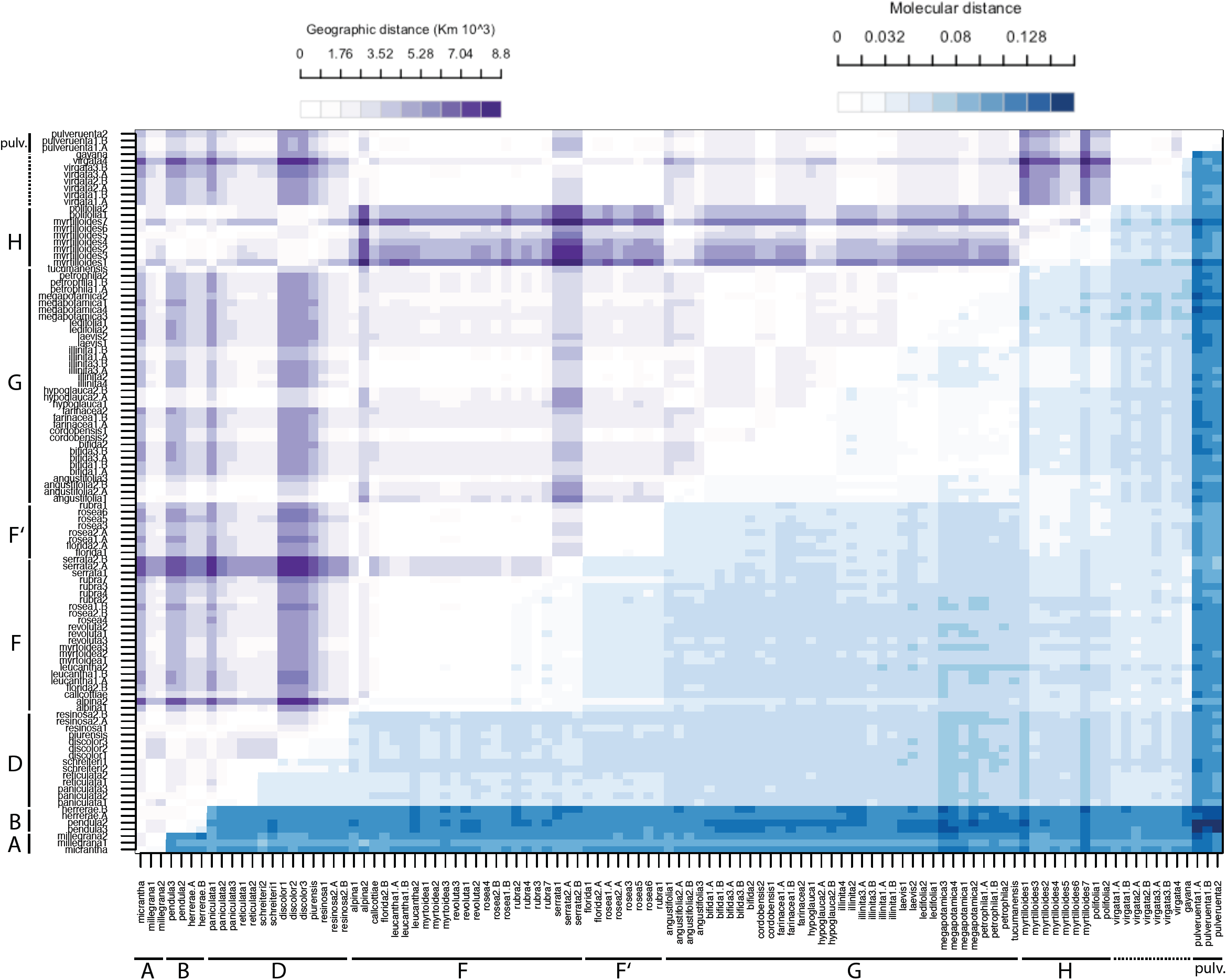
As in Figure 2, but for the NIA locus. For clades names, see Fig. 1.

**Fig. 4.**
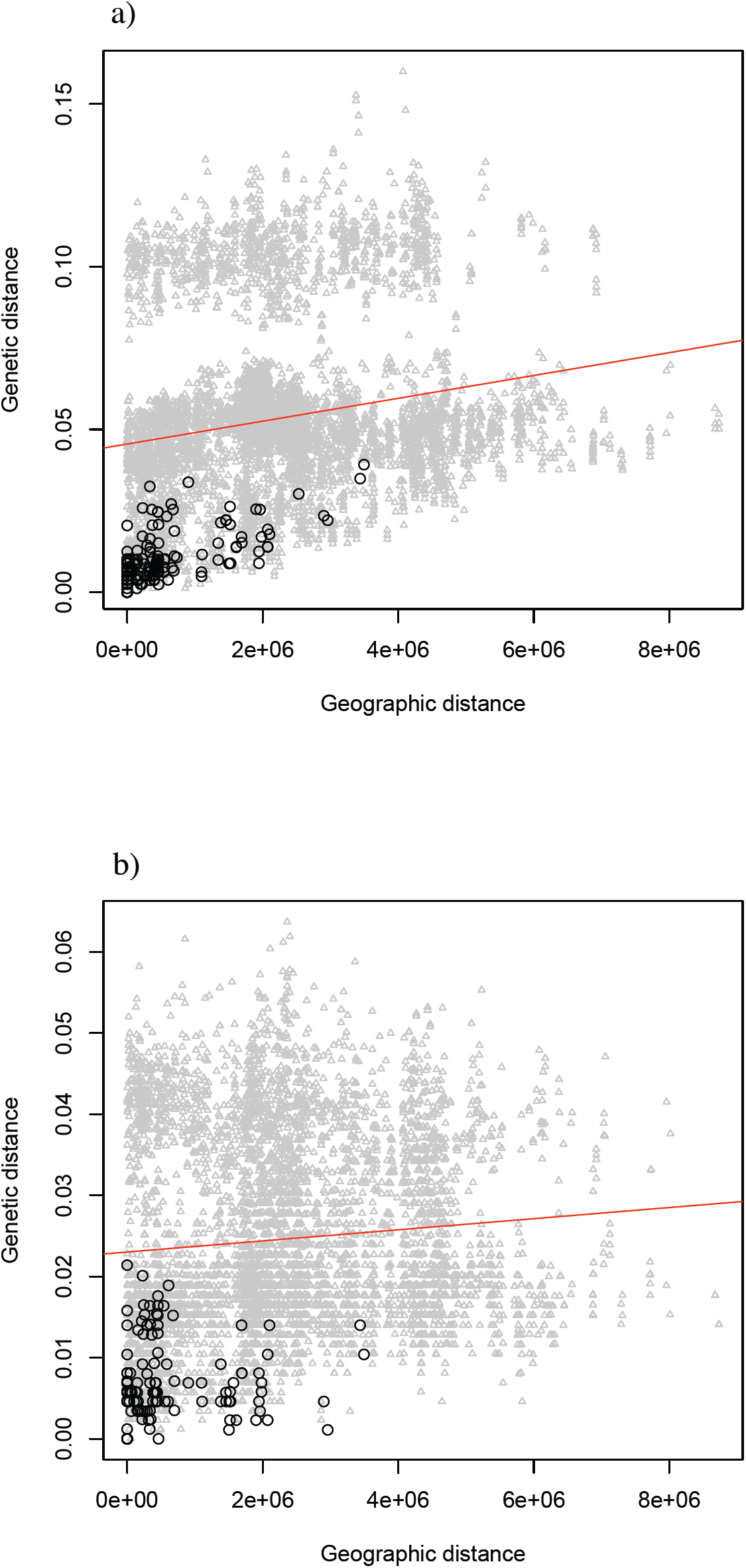
Results of simple Mantel test for a) MYC and b) NIA. Geographic distance (abscissa) in meters; genetic distance (ordinate) substitutions per site corrected according to the model of evolution used to reconstruct the haplotype trees (for details, see Zapata 2010). Gray triangles: pairwise comparisons among samples from different species; black circles: pairwise comparisons among samples within species. Red line: Mantel correlation of geographic and genetic distance matrices: MYC (*r* = 0.09, *P* = 0.064); NIA (*r* = 0.2, *P* = 0.002).

<u>Clade A.</u> This clade included *E. micrantha* and *E. millegrana* (Fig. 1), two allopatric species narrowly distributed in the dry inter Andean valleys of the Tropical Andes (Table 1, Appendix 3). *E. micrantha* (Fig. 5a) occurs in the valleys of northern Perú and *E. millegrana* in the valleys of central-south Bolivia (Fig. 5b). Molecular data for *E. micrantha* were available from only one individual; with this sampling, it was not possible to evaluate the genealogical exclusivity of this species. Phylogenetic analyses of MYC and NIA showed that the haplotypes of *E. millegrana,* sampled from northern and southern localities of its geographic range, were genealogically exclusive and concordant with geography (Fig. 1). The maximum level of sequence divergence between this pair of species was 0.015 in MYC and 0.020 in NIA for samples collected more than 2000 km apart (Fig. 1).

**Fig. 5.**
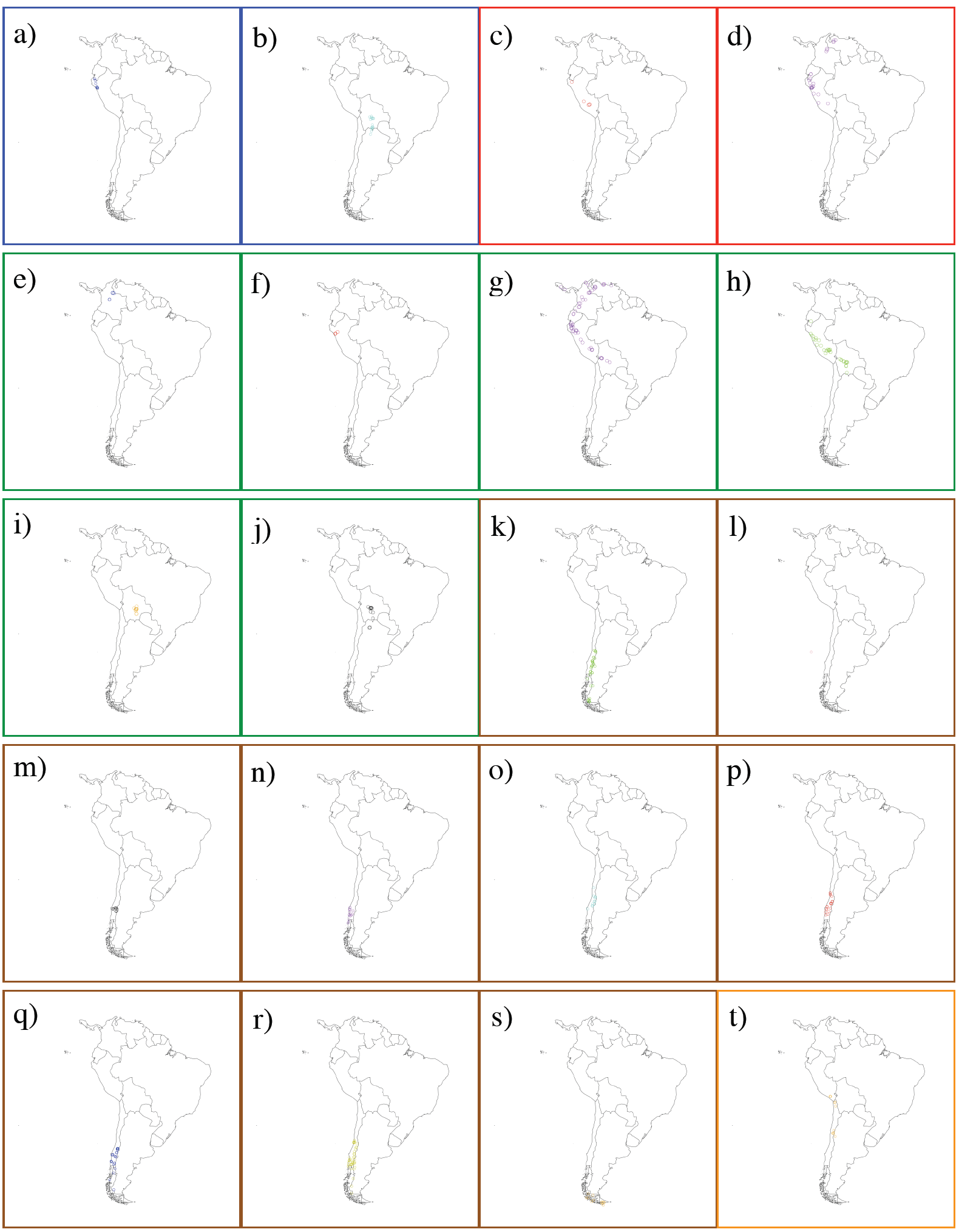
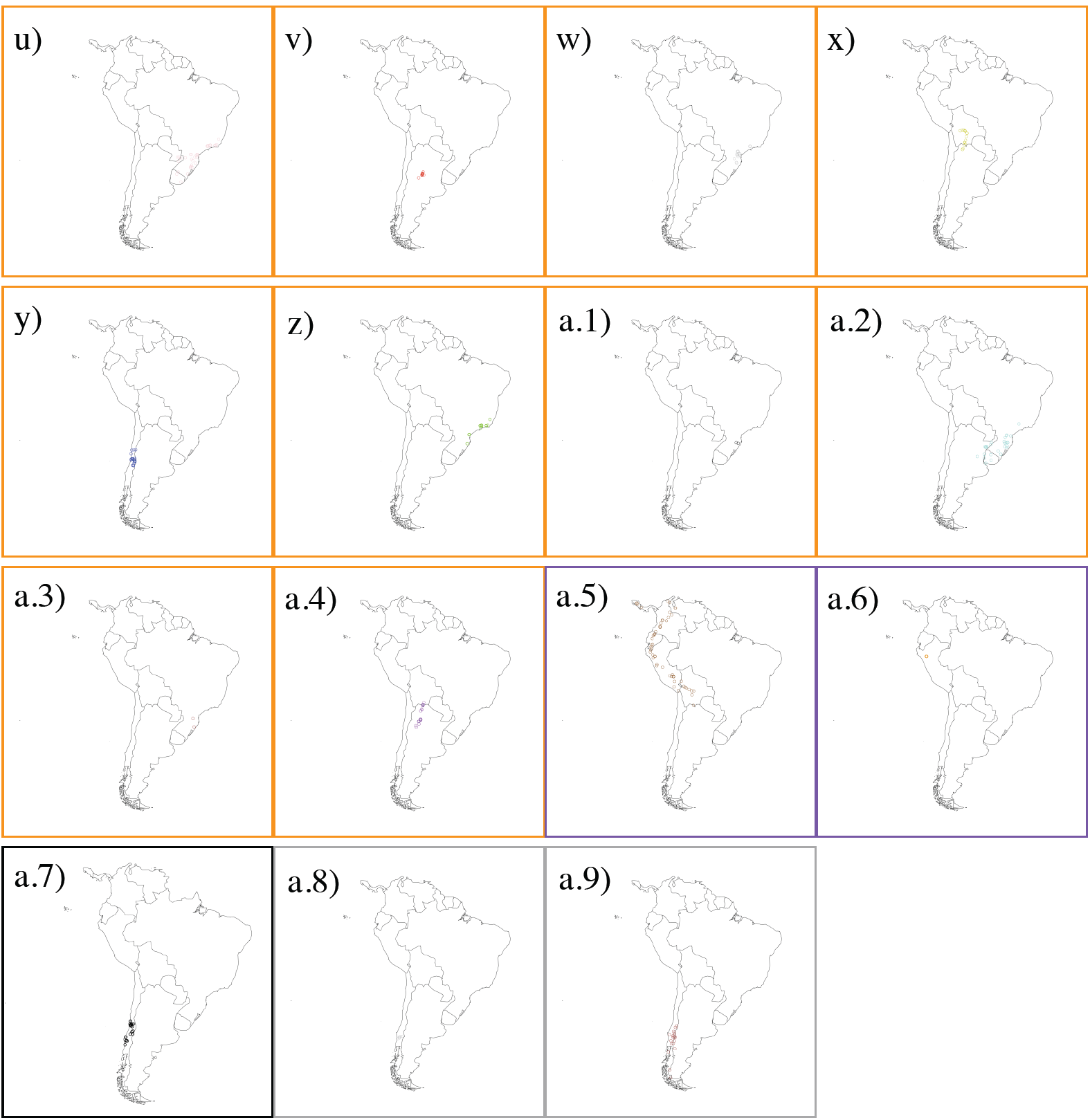
Geographic distribution of species of *Escallonia* included in this study. The individuals mapped are the individuals used for morphometric analyses (see Appendix 1, 2). Each map is surrounded by a colored box according to the color of the clade to which the species belong (for clade color, see Fig. 1). Each species is color-coded: a) *E. micrantha*: dark blue; b) *E. millegrana*: pale blue; c) *E. herrerae*: red; d) *E. pendula*: purple; e) *E. discolor*: dark blue; f) *E. piurensis*: red; g) *E. paniculata*: purple; h) *E. resinosa*; i) *E. reticulata*: orange; j) *E. schreiteri*: black; k) *E. alpina*: green; l) *E. callcottiae*: pink; m) *E. florida*: black; n) *E. leucantha*: purple; o) *E. myrtoidea*: pale blue; p) *E. revoluta*: red; q) *E. rosea*: dark blue; r) *E. rubra*: yellow; s) *E. serrata*: orange; t) *E. angustifolia*: orange; u) *E. bifida*: pink; v) *E. cordobensis*: red; w) *E. farinacea*: gray; x) *E. hypoglauca*: yellow; y) *E. illinita*: dark blue; z) *E. laevis*: green; a.1) *E. ledifolia*: black; a.2) *E. megapotamica*: pale blue; a.3) *E. petrophila*: brown; a.4) *E. tucumanensis*: purple; a.5) *E. myrtilloides*: brown; a.6) *E. polifolia*: red; a.7) *E. pulverulenta*: black; a.8) *E. gayana*: gray; a.9) *E. virgata*: brown.

Morphological data showed that the samples of *E. micrantha* and *E. millegrana* clearly separated in morphological PCA space (Fig. 6a). The elevation plot was bimodal (Fig. 7a), however the proportion plot revealed that proportions > 0.9 overlapped (Fig. 8a), implying that the frequency of intermediate phenotypes was > 0.1, and thus there was not enough evidence to suggest a morphological gap separating these species. No qualitative characters were fixed for alternative states in either species (Table 4, Appendix 2).

**TABLE 4.** Species profiles (see Appendix 2 for scoring of specimens; see Davis and Nixon, 1992) for the qualitative characters states. For polymorphic states, all the states are shown. N: sample size for each species. Significant results of Wiens and Servedio (2000) test with *. For character states and abbreviations, see Table 2. For clade names, see Fig. 1.

| Clade | Species | N | HABIT | ADLAMGLA | ADLAMPUB | ABLAMGLA | ABLAMPUB | LAMMARG | MARGAPP | PEDGLAND | PEDPUB | OVAGLAN | OVAPUB | DIS<br>K | FLOCOLOR |
| --- | --- | --- | --- | --- | --- | --- | --- | --- | --- | --- | --- | --- | --- | --- | --- |
| **A** | *E. millegrana* | 17 | 0-1 | 0-1 | 0-1 | 0-1 | 0-1 | 1-2-5 | 0 | 0-1 | 1 | 0-1 | 1 | 0-1 | 2 |
|  | *E. micrantha* | 11 | 0-1 | 0-1 | 0-1 | 0-1 | 1 | 1-5 | 0 | 0-1 | 1 | 0-1 | 1 | 0-1 | 2-3 |
| **B** | *E. herrerae* | 5 | 0 | 1 | 0-1 | 0-1 | 0-1 | 0-2 | 0 | 1 | 0-1 | 1 | 0-1 | 0 | 1-2-3-4 |
|  | *E. pendula* | 28 | 0-1 | 0-1 | 0-1 | 1 | 0-1 | 0-2-5 | 0 | 0-1 | 0-1 | 0-1 | 0-1 | 0 | 1-2-3-4 |
| **D** | *E. piurensis* | 5 | 0-1 | 0-1 | 0-1 | 0-1 | 0-1 | 2 | 0 | 1 | 1 | 0-1 | 0-1 | 0 | 2 |
|  | *E. discolor* | 5 | 0-1 | 0 | 0-1 | 0-1 | 0-1 | 0-6 | 0 | 0-1 | 1 | 0-1 | 1 | 0 | 2 |
|  | *E. resinosa* | 35 | 0-1 | 0-1 | 0-1 | 0-1 | 0-1 | 2 | 0 | 0 | 1 | 0-1 | 0-1 | 0 | 2 |
|  | *E. reticulata* | 11 | 0-1 | 0 | 0-1 | 1 | 0-1 | 6 | 0 | 1 | 1 | 1 | 0-1 | 0 | 2 |
|  | *E. schreiteri* | 12 | 0-1 | 0-1 | 0-1 | 0-1 | 0-1 | 6 | 0 | 0-1 | 1 | 0-1 | 0-1 | 0 | 2 |
|  | *E. paniculata* | 61 | 0-1 | 0 | 0-1 | 0-1 | 0-1 | 2-6 | 0 | 0-1 | 0-1 | 0-1 | 0-1 | 0 | 2 |
| **F** | *E. serrata* | 13 | 2 | 0 | 0-1 | 0 | 0 | 1 | 0 | 0 | 0 | 0 | 0 | 1 | 2-3 |
|  | *E. rosea* | 29 | 1 | 0-1 | 1 | 0-1 | 0-1 | 1 | 0 | 0 | 0-1 | 0 | 0-1 | 1 | 1-2-3 |
|  | *E. alpina* | 39 | 1-2 | 0-1 | 0-1 | 0-1 | 0-1 | 1-3 | 0 | 0-1 | 0-1 | 0-1 | 0-1 | 0-1 | 1-2-3 |
|  | *E. virgata* | 19 | 1-2 | 0 | 0-1 | 0 | 0 | 1-2 | 0 | 0 | 0-1 | 0-1 | 0-1 | 0 | 2-3 |
|  | *E. florida* | 10 | 1 | 0 | 0-1 | 0 | 0-1 | 1 | 0 | 0 | 0-1 | 0 | 0-1 | 0 | 2 |
|  | *E. revouta* | 20 | 0-1 | 0-1 | 0-1 | 0-1 | 0-1 | 1 | 1 | 0-1 | 0-1 | 0-1 | 0-1 | 1 | 2-3 |
|  | *E. myrtoidea* | 13 | 0-1 | 0 | 0-1 | 0-1 | 0-1 | 1 | 0 | 0-1 | 0-1 | 0-1 | 0-1 | 1 | 2-3 |
|  | *E. rubra* | 46 | 1-2 | 0-1 | 0-1 | 1 | 0-1 | 1-3 | 0 | 0-1 | 0-1 | 1 | 0-1 | 1 | 1-3 |
|  | *E. leucantha* | 12 | 1 | 0 | 0-1 | 0-1 | 0-1 | 1-2 | 0 | 0 | 1 | 0 | 1 | 1 | 2 |
|  | *E. callicottiae* | 6 | 1 | 0 | 1 | 0-1 | 0 | 1 | 0 | 0 | 0-1 | 0 | 0-1 | 0-1 | 1-2-3 |
|  | *E. gayana* | 6 | 1 | 0 | 0-1 | 0 | 0-1 | 1-2 | 0 | 0-1 | 1 | 0 | 1 | 0 | 2 |
| **G** | *E. angustifolia* | 13 | 0-1 | 1 | 0-1 | 1 | 0-1 | 1-6 | 0 | 1 | 0-1 | 1 | 0-1 | 0-1 | 1-2 |
|  | *E. illinita* | 21 | 1 | 1 | 0-1 | 1 | 0-1 | 1-2-6 | 1 | 1 | 0-1 | 1 | 0-1 | 1 | 2-3 |
|  | *E. laevis* | 26 | 0-1-2 | 0-1 | 0-1 | 0-1 | 0-1 | 1-3-6 | 0 | 0-1 | 0-1 | 1 | 0-1 | 0 | 1-2-3 |
|  | *E. petrophila* | 2 | 1 | 0 | 0 | 0 | 0 | 1 | 0 | 0 | 0 | 0 | 0 | 0 | 1 |
|  | *E. ledifolia* | 2 | 1-2 | 0 | 1 | 1 | 1 | 2 | 0 | 1 | 1 | 1 | 1 | 0 | 1 |
|  | *E. cordobensis* | 11 | 0-1 | 0 | 0-1 | 0-1 | 0-1 | 1 | 0 | 0-1 | 0-1 | 0-1 | 0 | 0 | 2 |
|  | *E. megapotamica* | 27 | 1 | 0-1 | 0-1 | 1 | 0-1 | 1-2 | 0 | 0-1 | 0-1 | 0-1 | 0-1 | 0 | 2 |
|  | *E. hypoglauca* | 19 | 0-1 | 0-1 | 0-1 | 0-1 | 0-1 | 1-3 | 0 | 0-1 | 0-1 | 1 | 1 | 0 | 2-3 |
|  | *E. tucumanensis* | 18 | 0-1 | 0-1 | 0-1 | 0-1 | 0-1 | 1-3 | 0 | 0-1 | 0-1 | 0-1 | 0-1 | 0 | 2 |
|  | *E. bifida* | 36 | 0-1 | 0 | 0-1 | 0-1 | 0-1 | 2-6 | 0 | 0-1 | 1 | 0-1 | 1 | 0 | 2 |
|  | *E. farinacea* | 17 | 1 | 1 | 0-1 | 1 | 0-1 | 1-2 | 0 | 1 | 0-1 | 1 | 0-1 | 0 | 2 |
| **H** | *E. polifolia\** | 9 | 1 | 0-1 | 1 | 0 | 1 | 0 | 1 | 1 | 1 | 1 | 1 | 0 | 2-4 |
|  | *E. myrtilloides* | 55 | 0-1 | 0 | 0-1 | 0-1 | 0-1 | 1-2-6 | 0 | 1 | 0-1 | 1 | 0-1 | 0-1 | 2-4 |
| **pulv.** | *E. pulverulenta* | 20 | 1 | 0-1 | 0-1 | 0-1 | 0-1 | 2-3-5 | 0 | 0-1 | 0-1 | 1 | 0-1 | 0-1 | 2 |

**Fig. 6.**
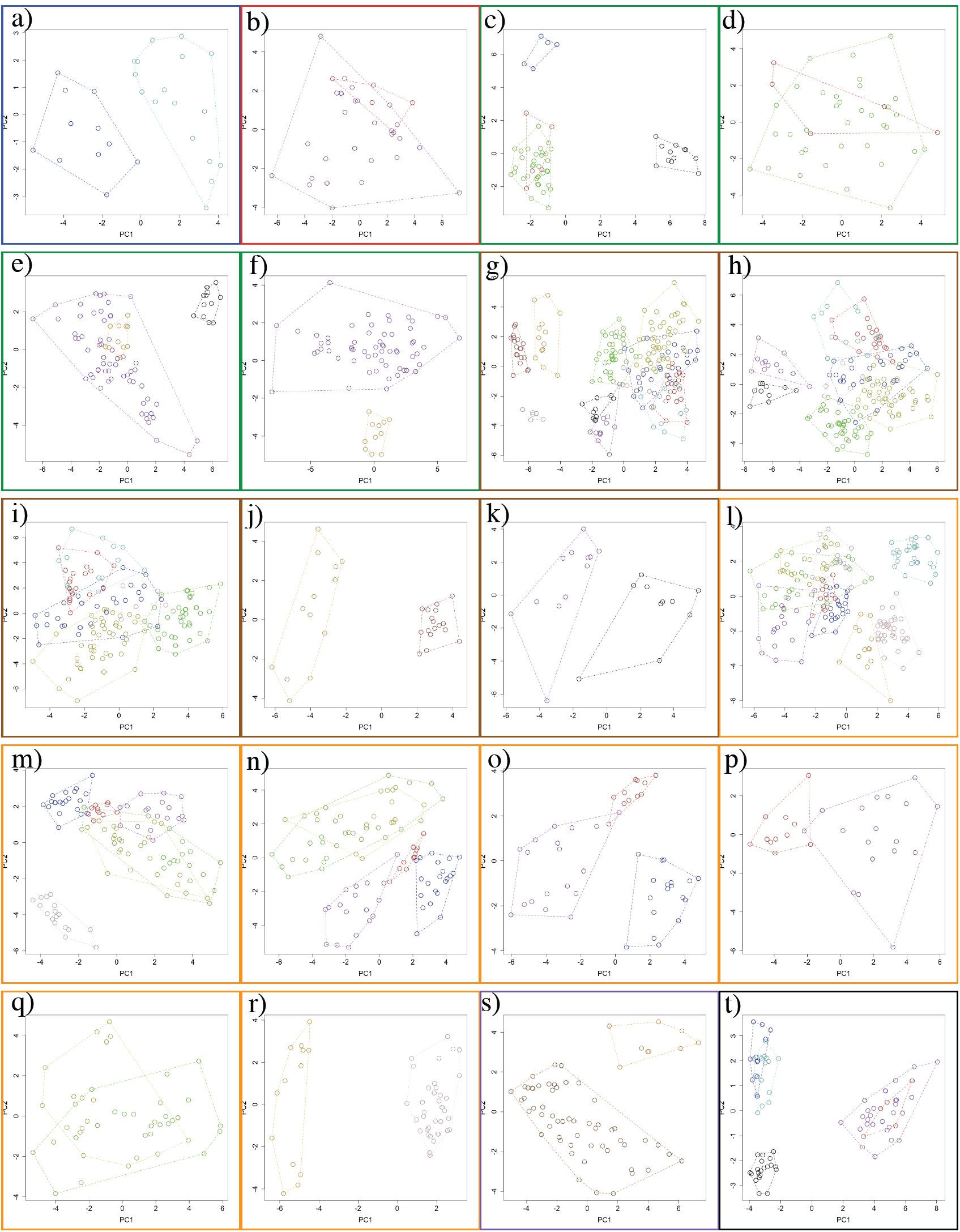
Principal Component Analyses (PCA) describing the pattern of morphological variation for species within clades. Each PCA is surrounded by a colored box according to color of the clades to which the species in the morphological PCA space belong (for clade color, see Fig. 1). Each species is color-coded (for species color, see Fig 5). The dashed polygons correspond to the minimum convex hulls for the samples of each species. For clades D, F, and G, several PCA were used because samples of several species overlapped in initial and subsequent PCA (see text for details).

**Fig. 7.**
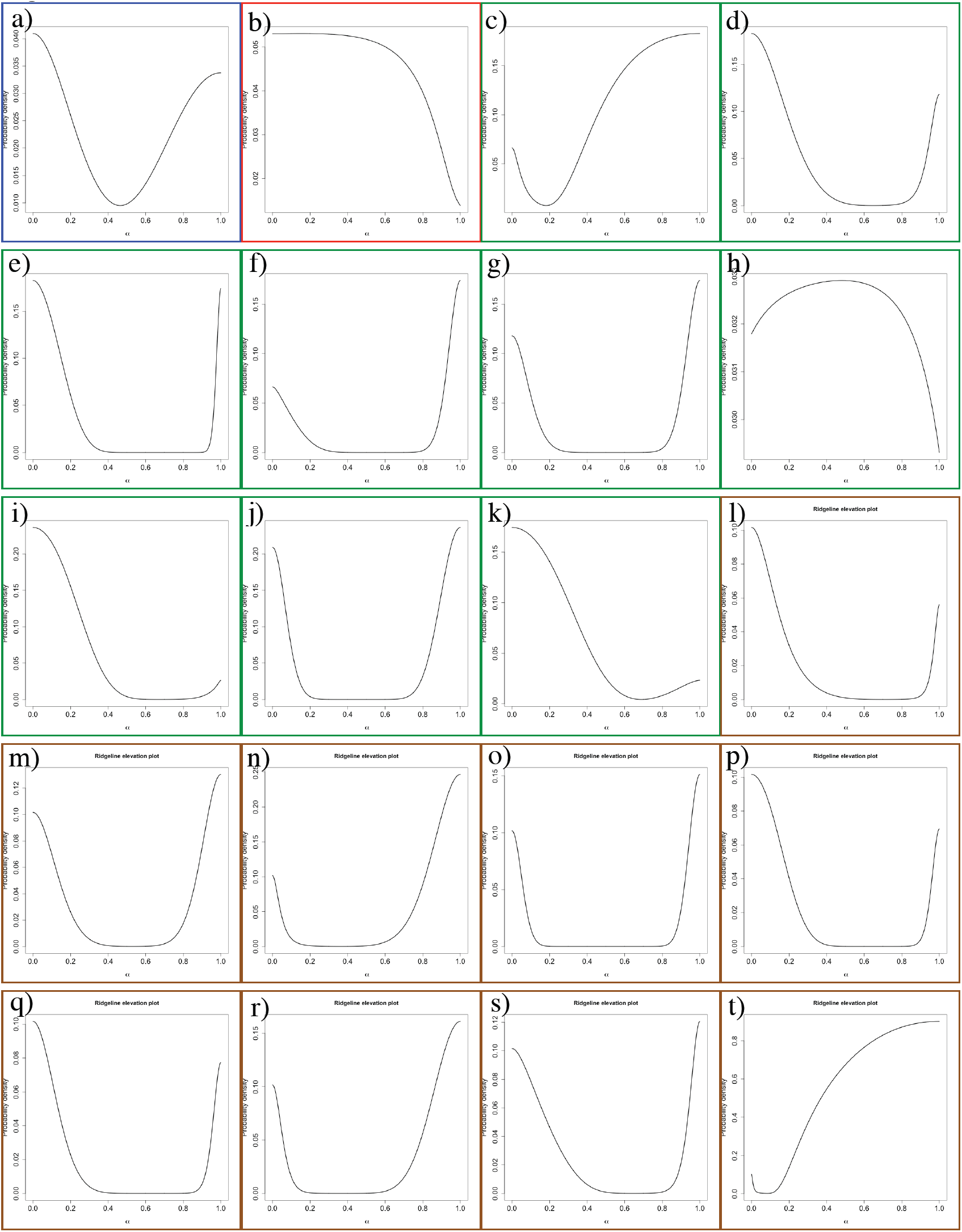
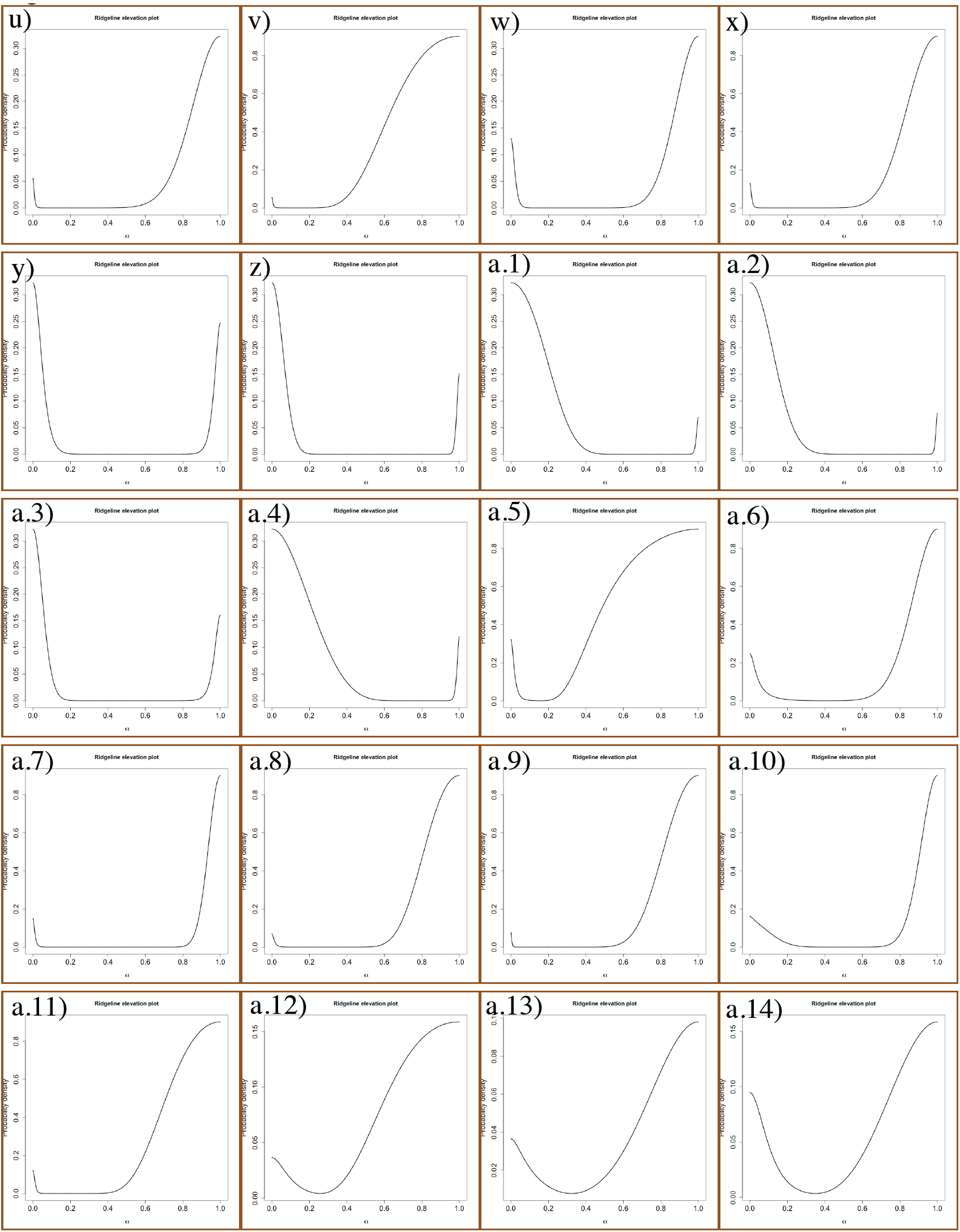
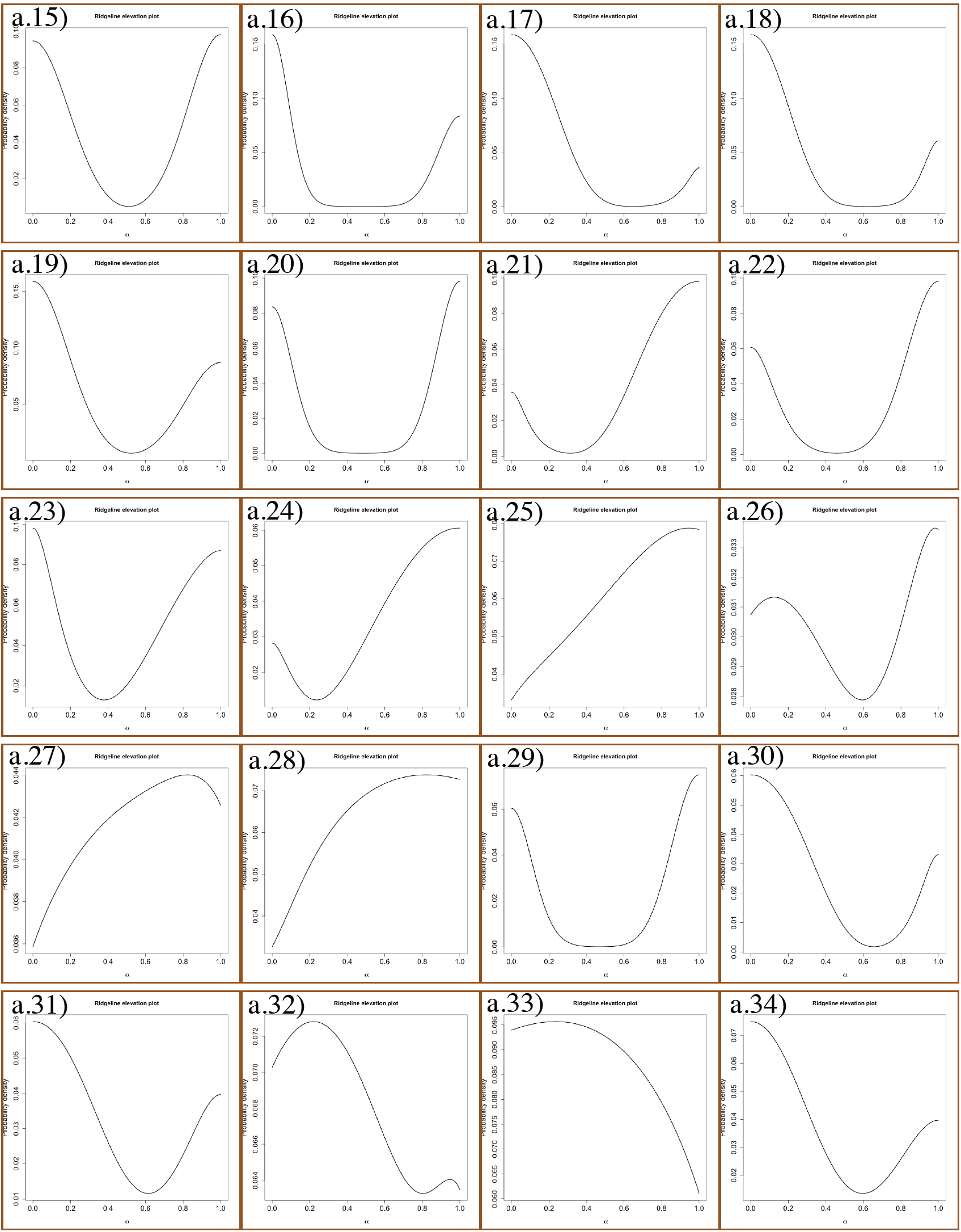
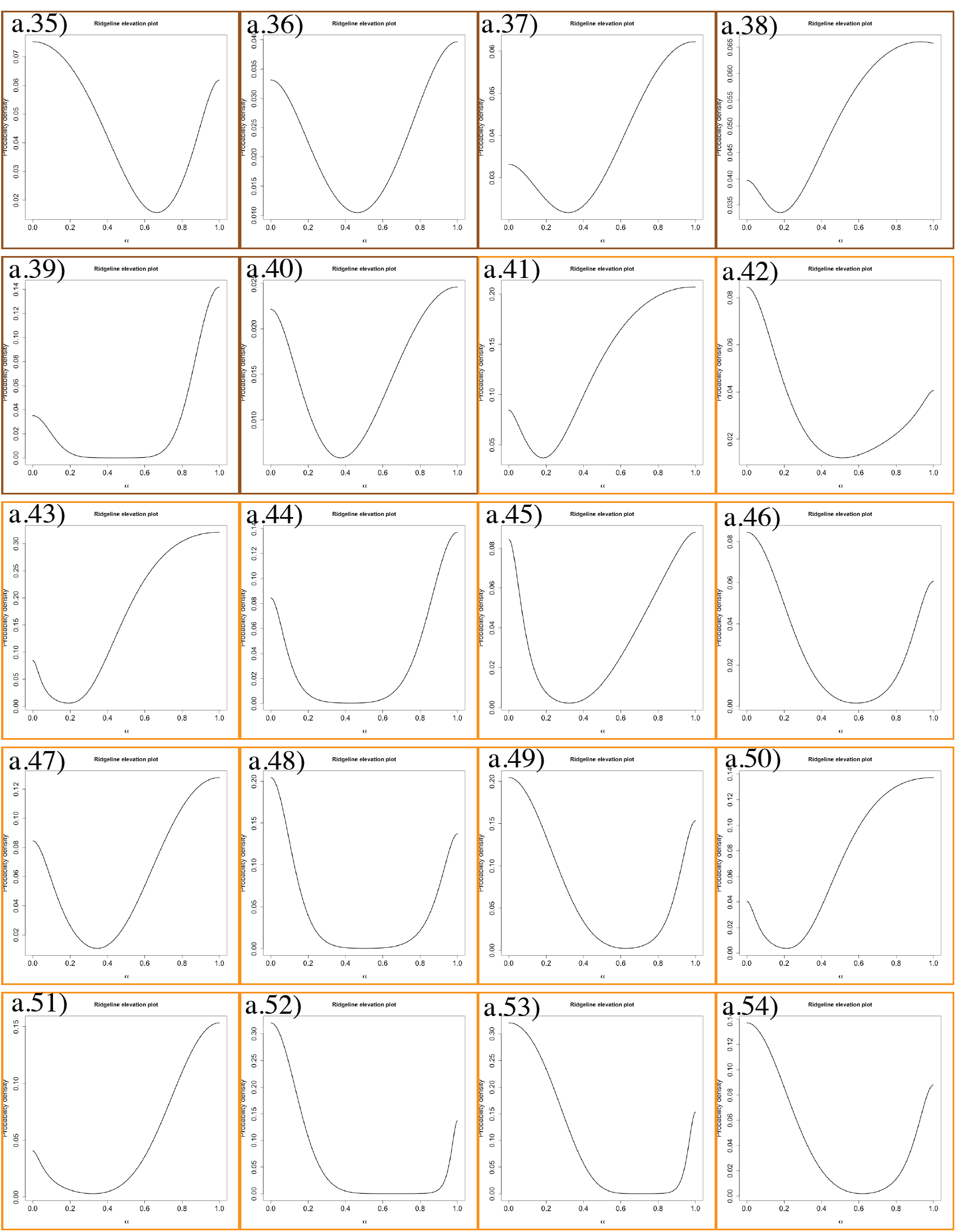
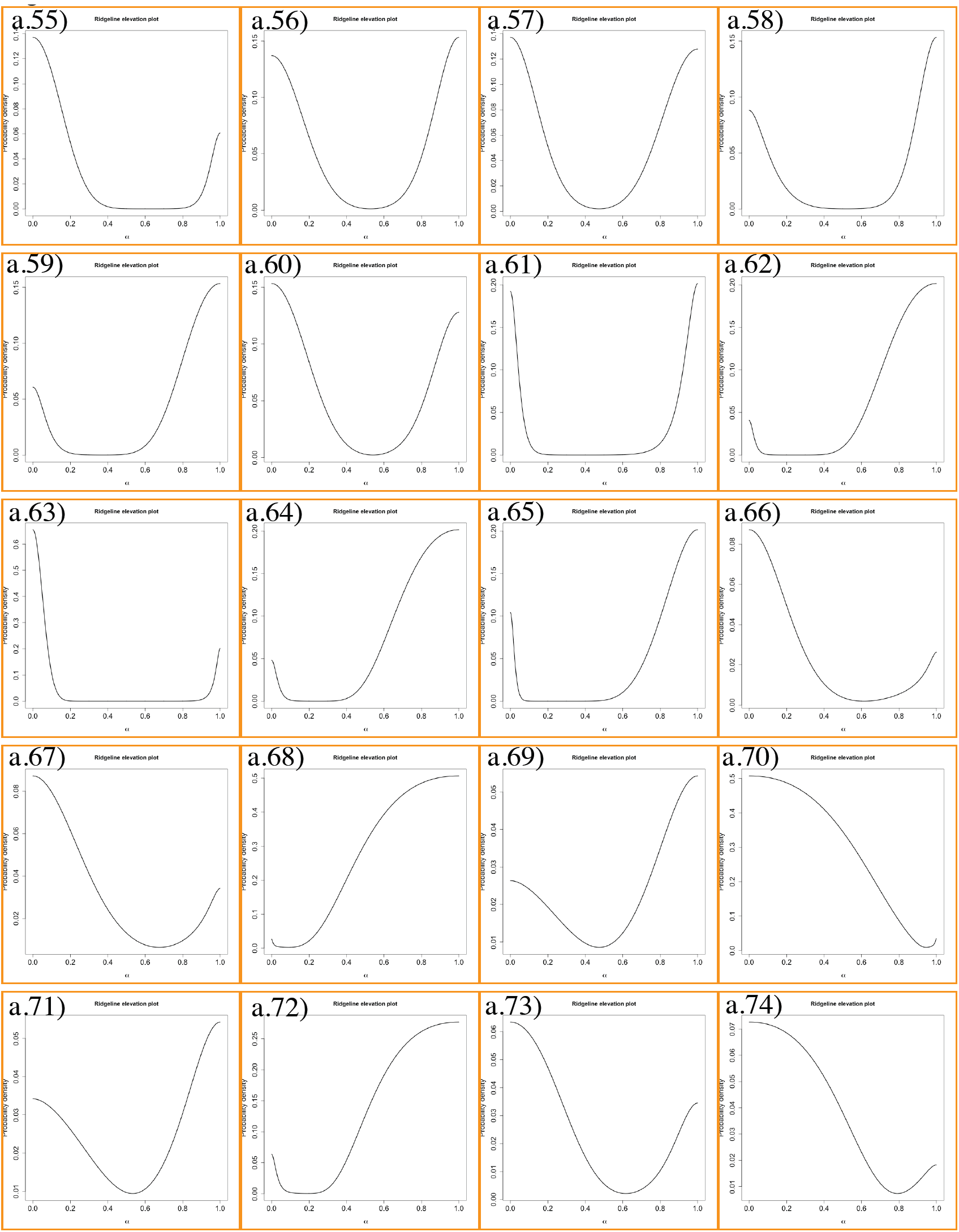
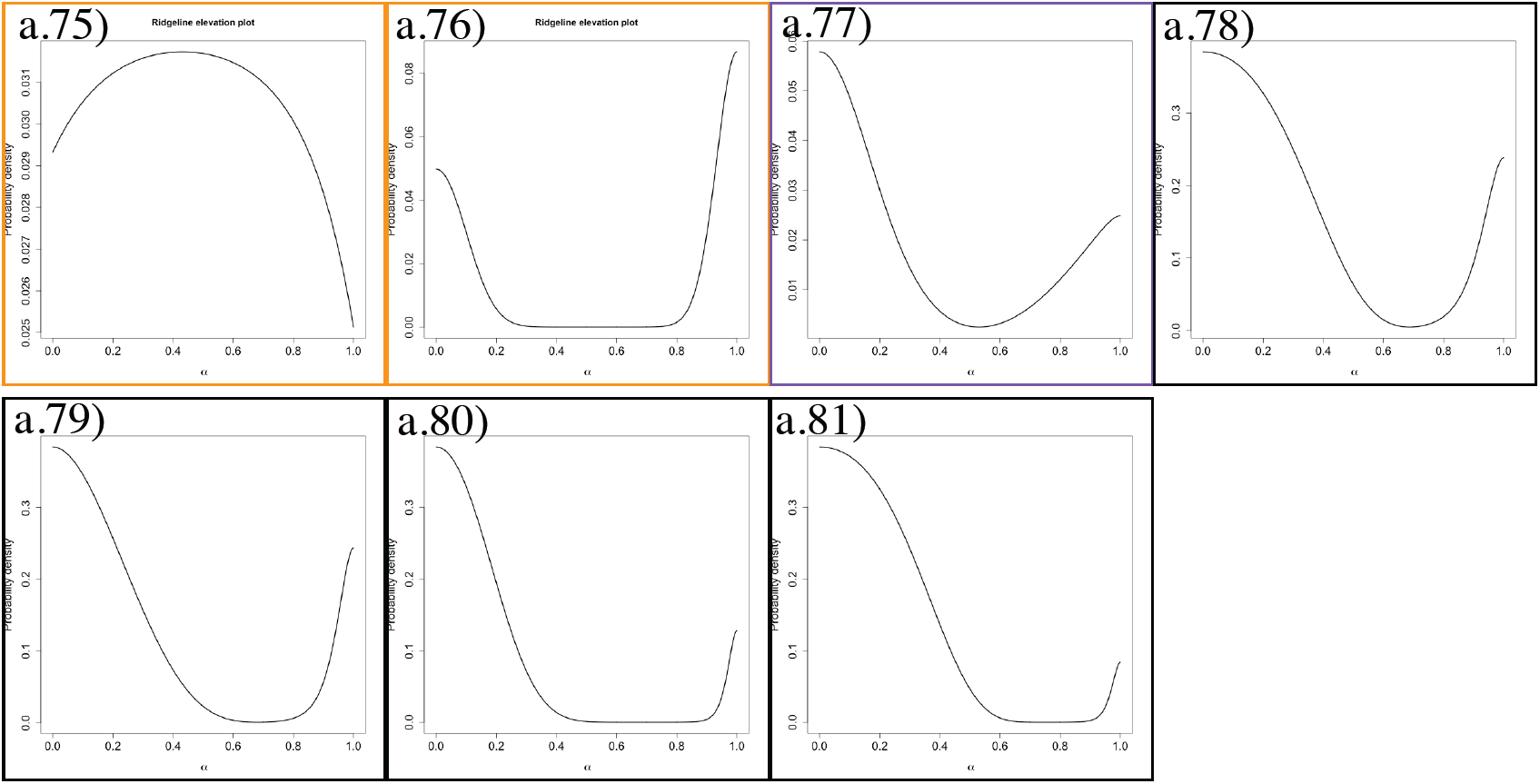
Estimated probability density function (*pdf*) of the mixture of the distributions describing the pattern of morphological variation of pairs of species along the ridgeline manifold (here referred to as “elevation plots”), for all pairwise species comparisons within clades. The ridgeline manifold (abscissa, a) ranges from zero (the bivariate mean of one species) to one (the bivariate mean of the other species). Each elevation plot is surrounded by a colored box according to the color of the clade to which the species being compared belong (for clade color, see Fig. 1). See Fig. 8 for species pairs shown in each elevation plot. For details on elevation plots, see Zapata and Jiménez (2012).

**Fig. 8.**
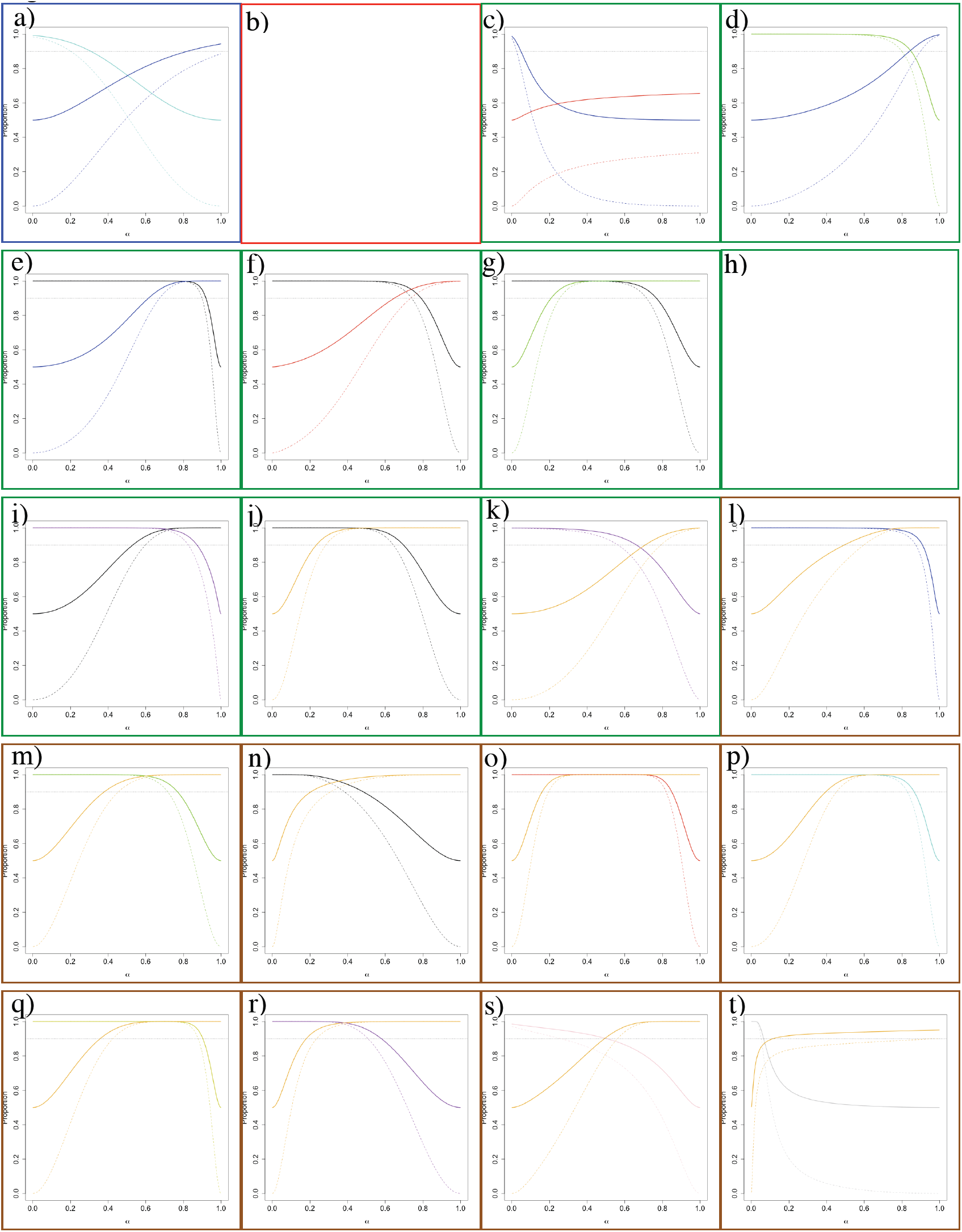
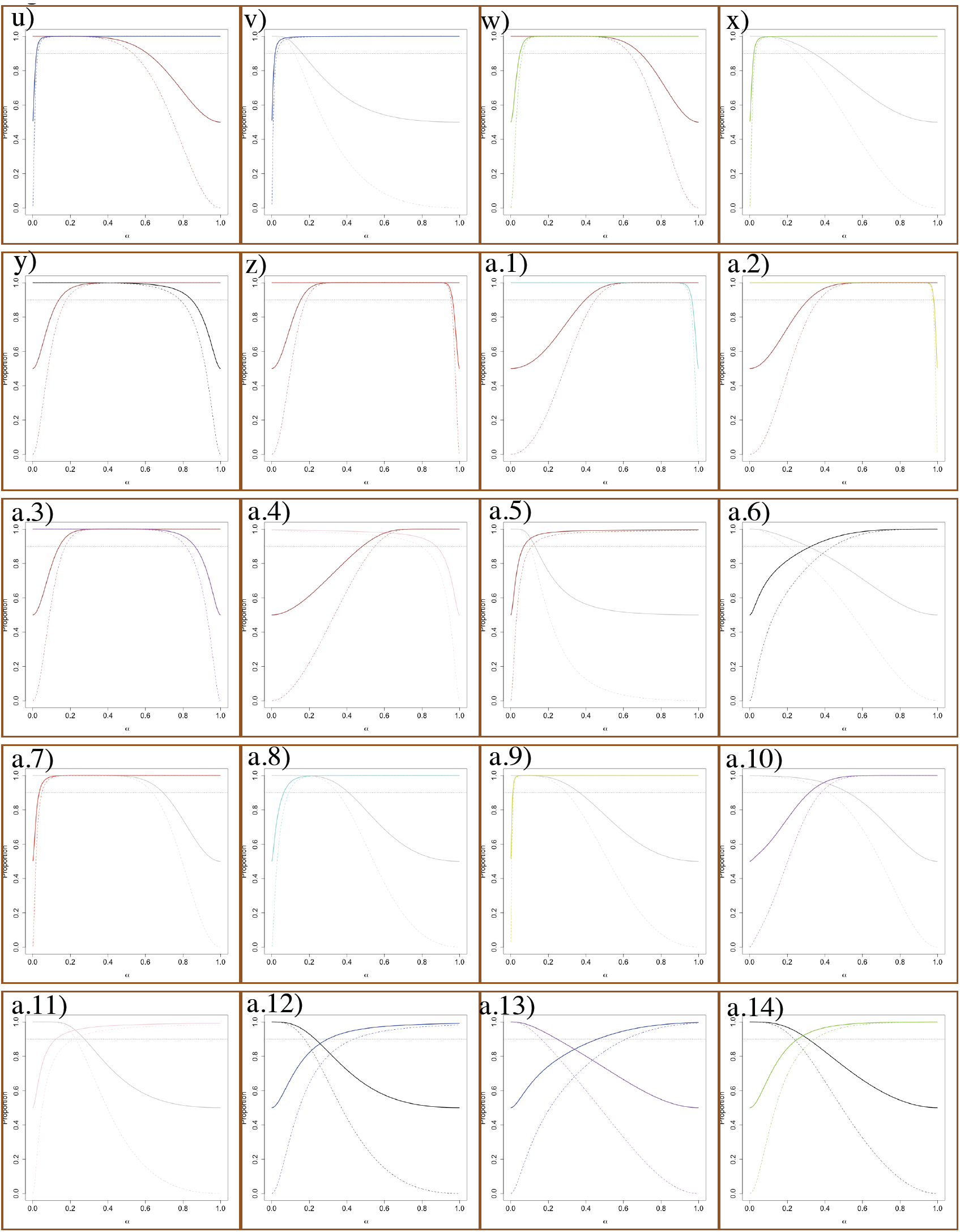
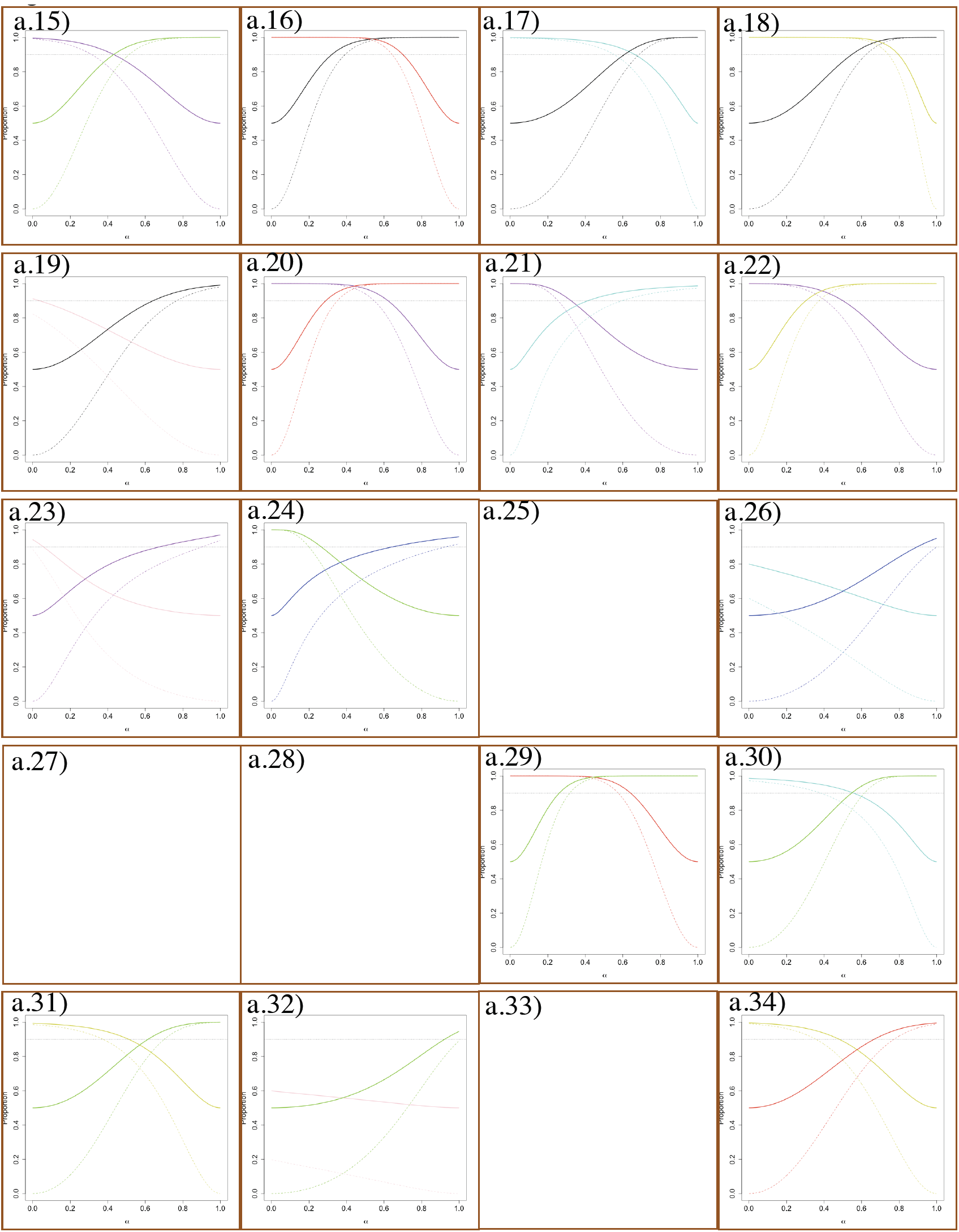
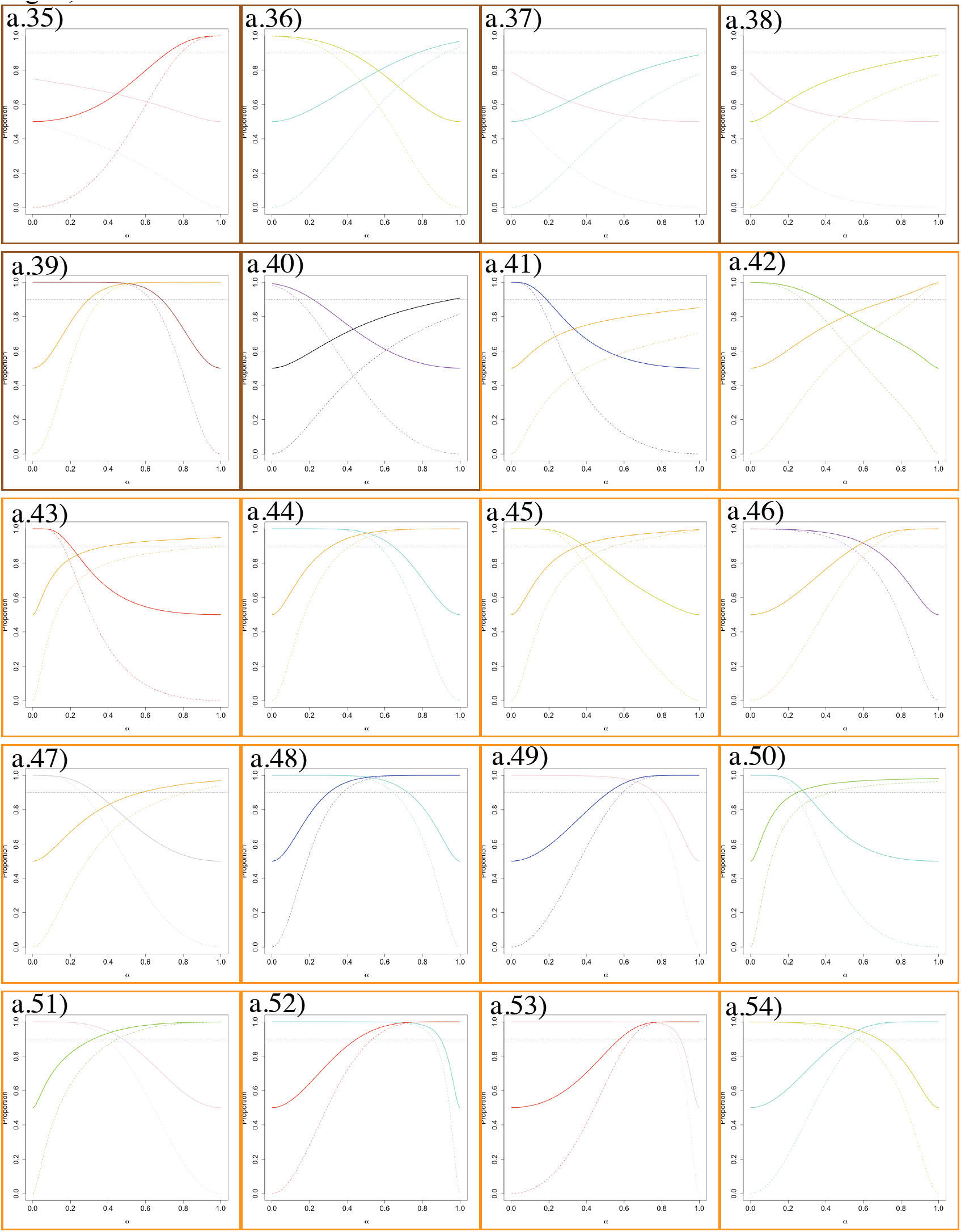
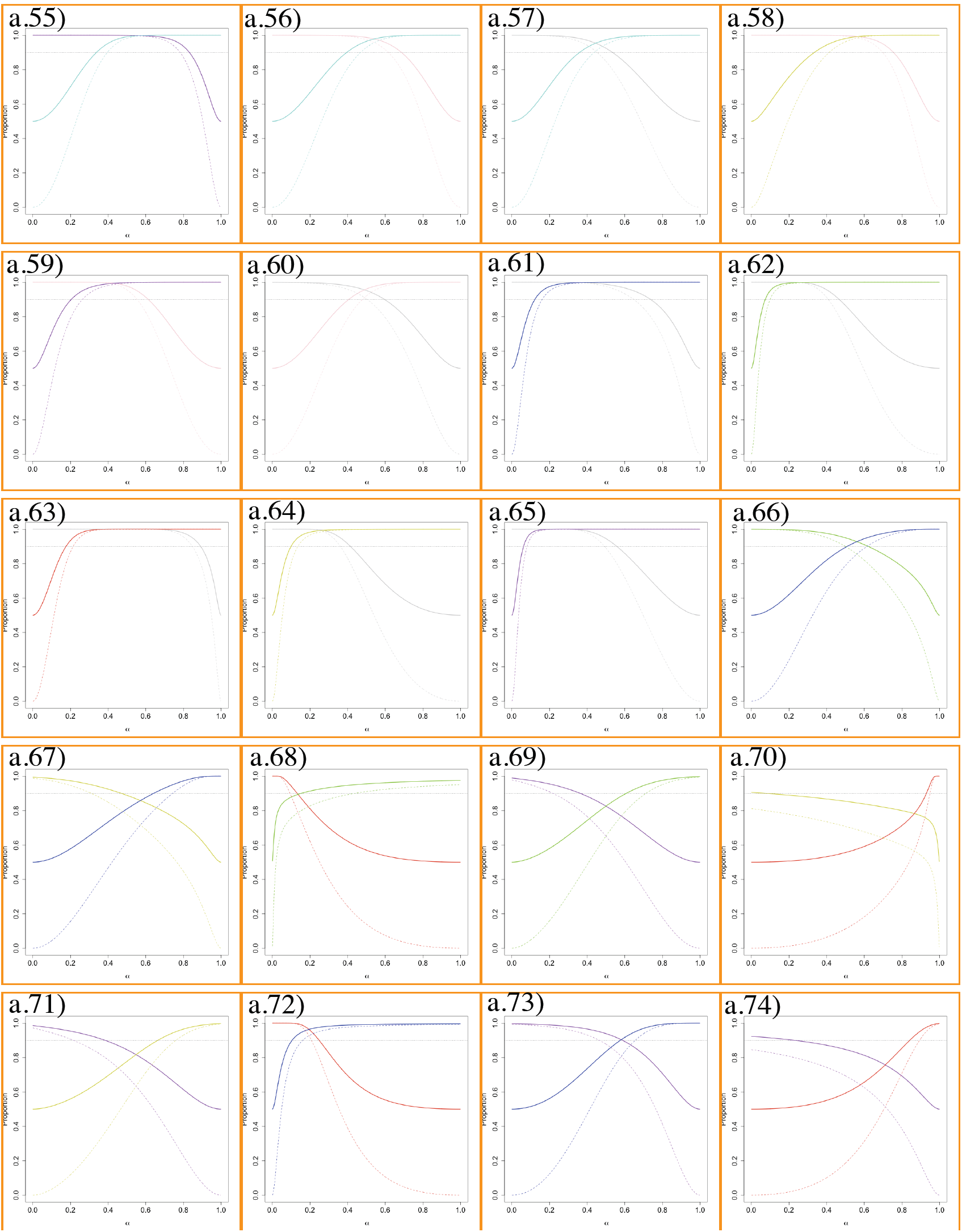
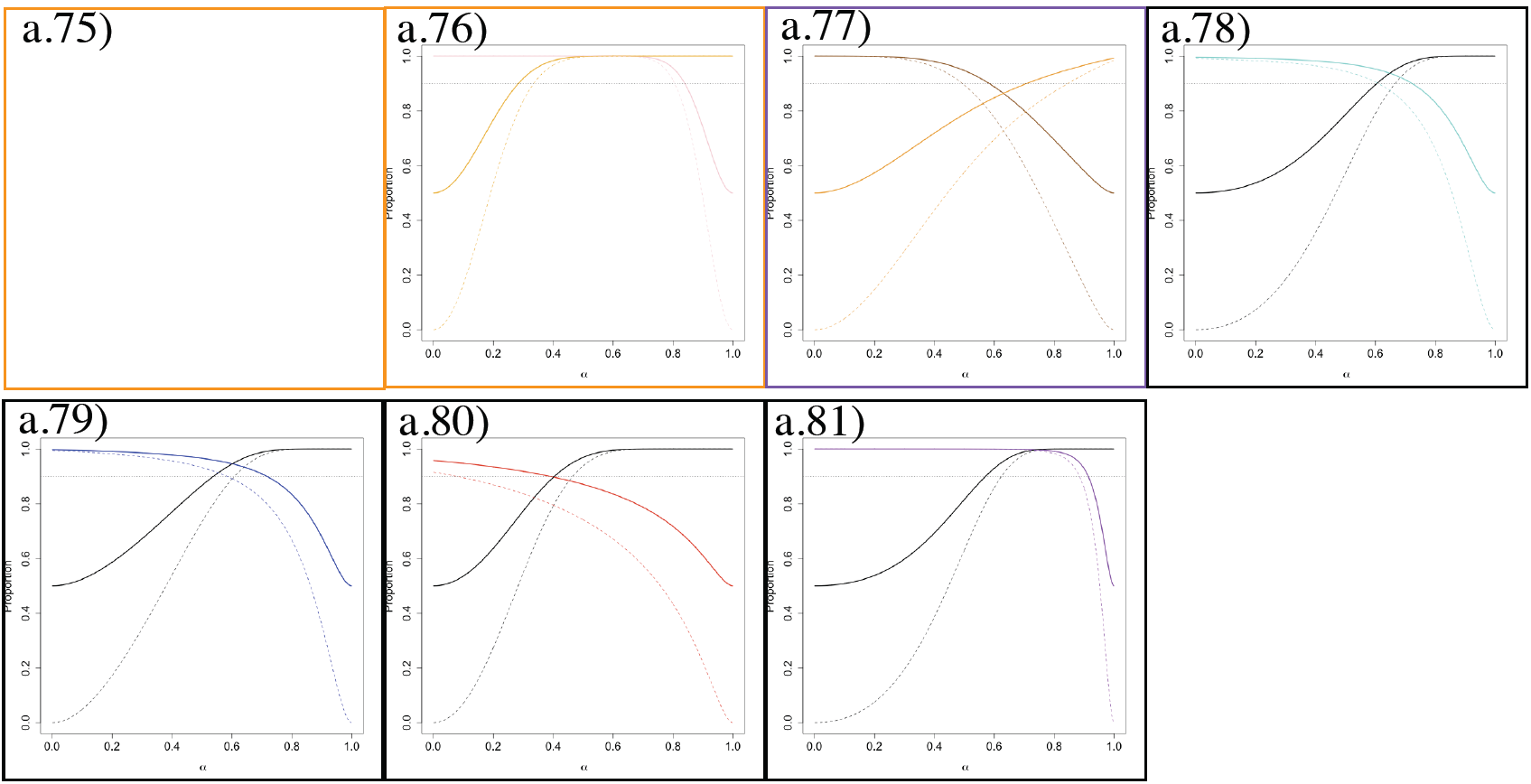
Estimated proportions of the distributions describing the pattern of morphological variation of pairs of species along the ridgeline manifold (here referred to as “proportion plots”), for all pairwise species comparisons within clades. The dashed lines correspond to the proportions of the distributions describing the pattern of morphological variation covered by bivariate ellipsoid tolerance regions (γ = 0.95) for each species sharing a point along the ridgeline manifold. The continuous lines correspond to the proportions of the areas of the distributions describing the pattern of morphological variation for each species, which lie on either side of the tangent line that splits morphological space on the point shared by ellipsoid tolerance regions along the ridgeline manifold. The dotted line corresponds to the proportion = 0.9 (i.e., a frequency cutoff of 0.1). Each elevation plot is surrounded by a colored box according to the color of the clade the species being compared belong to (for clade color, see Fig. 1). Dashed and continuous lines are color coded for each species (for species color, see Fig. 5). Each panel corresponds to a panel in Fig. 7 and both figures should be analyzed together. For details on proportion plot, see Zapata and Jiménez (2012).

Bioclimatic data revealed that samples of *E. micrantha* and *E. millegrana* separated strongly, but not completely, in bioclimatic PCA space (Fig. 9a). The PC scores differed significantly between these two species (Pillai’s trace = 0.731; *F*_2, 25_ = 34.055; *P* < 0.000) with significant difference found only along PC1 (*F*_1_ = 59.957; *P* < 0.000). This suggests there is evidence that *E. micrantha* and *E. millegrana* occur in significantly different environments, with *E. micrantha* occurring in slightly wetter, colder and less seasonal places than *E. millegrana* (Table 5).

**TABLE 5.**
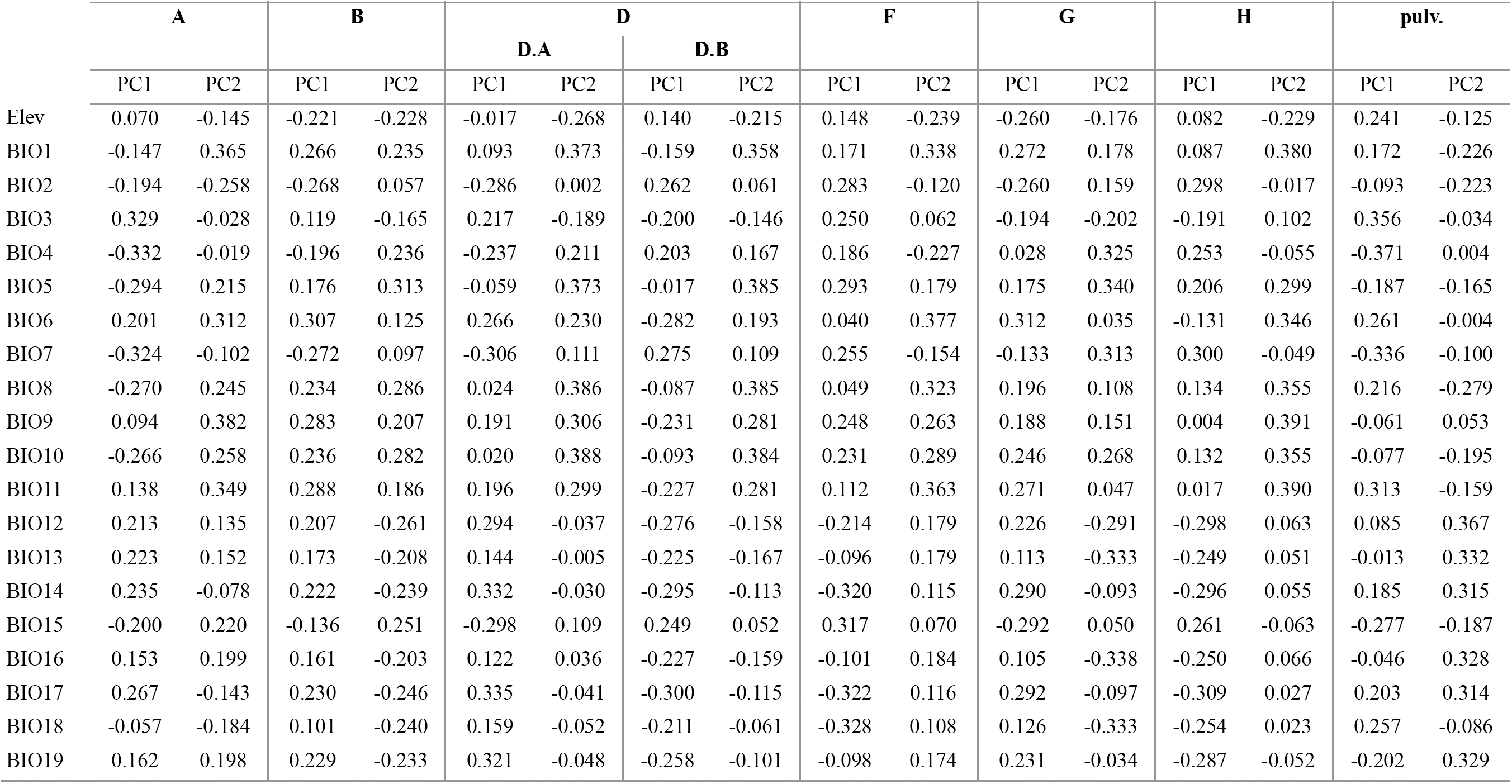
Loadings of elevation and bioclimatic variables on PC1 and PC2 for each PCA within clade. For clade names, see Fig. 1. For bioclimatic variable names, see Table 2.

**Fig. 9.**
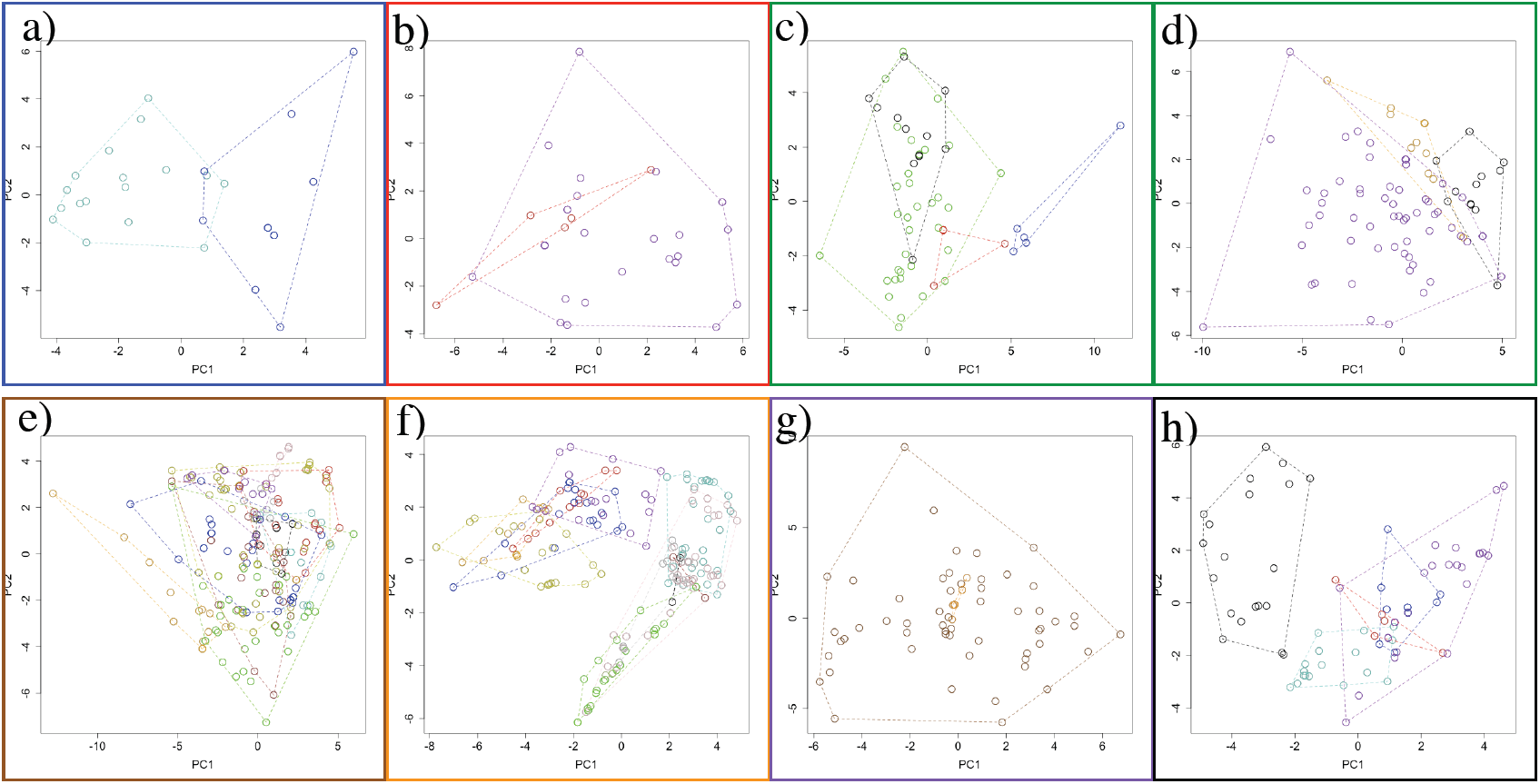
Principal Component Analyses (PCA) describing the pattern of ecological variation for species within clades. Each PCA is surrounded by a colored box according to color of the clades to which the species in the bioclimatic PCA space belong (for clade color, see Fig. 1). Each species is colored-coded (for species color, see Fig 5). The dashed polygons correspond to the minimum convex hulls for the samples of each species. For clade D: a) PCA for clade D.A; b) PCA for clade D.B (see text for details).

Both species overlapped in the range of flowering time, although there was a tendency for *E. micrantha* to flower slightly later than *E. millegrana* (Fig. 10a, b).

**Fig. 10.**
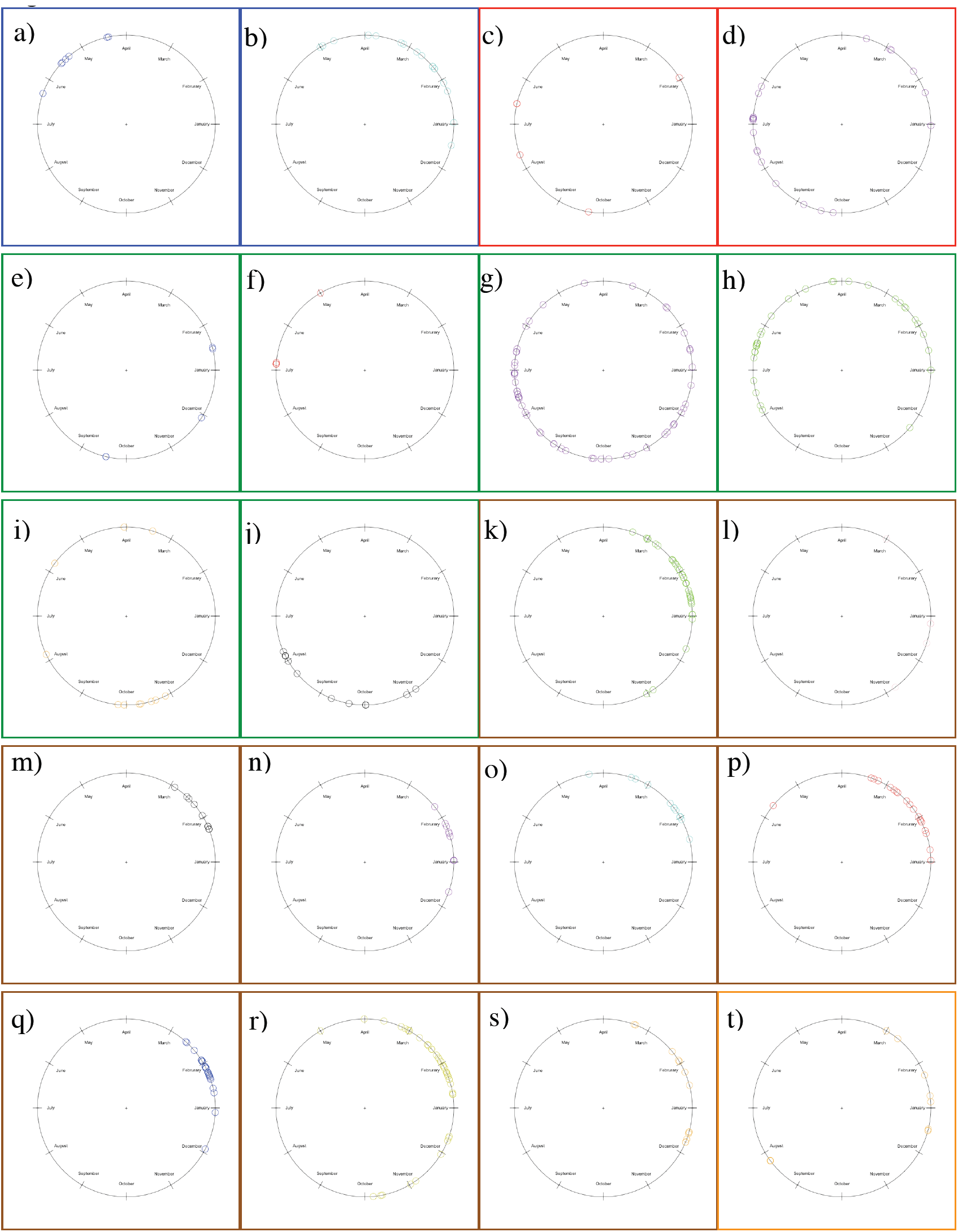
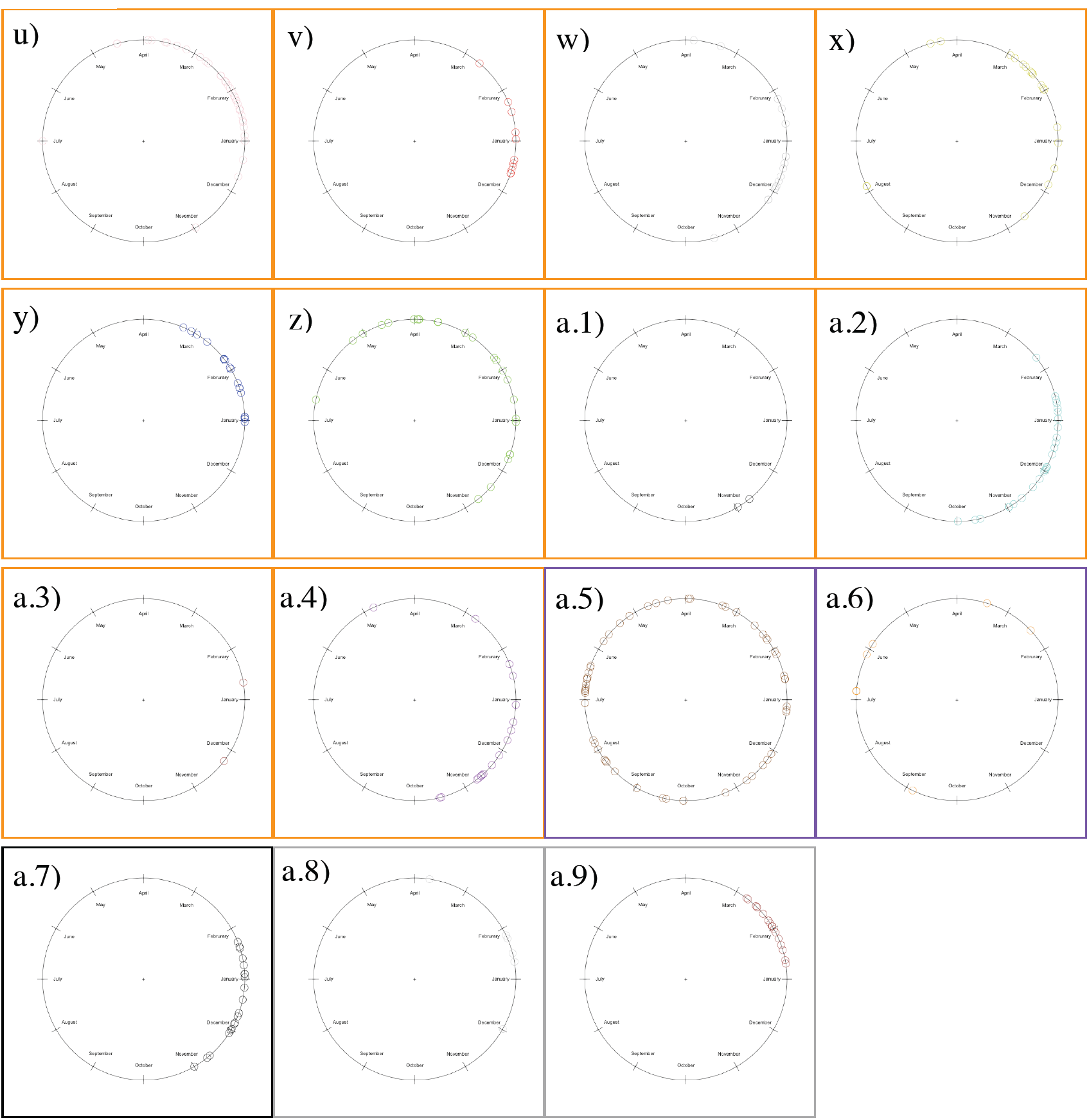
Flowering time of species of *Escallonia* included in this study. The individuals plotted are the individuals used for morphometric analyses (see Appendix 1, 2). Each plot is surrounded by a colored box according to the color of the clade the species belongs to (for clade color, see Fig. 1). In all circular plots, 0°: January; 90°: April; 180°: July; 270°: October.

<u>Clade B.</u> This clade included the pair *E. herrerae* and *E. pendula* (Fig. 1), two species occurring in the dry inter Andean valleys of the Tropical Andes (Table 1, Appendix 3). While *E. herrerae* (Fig. 5c) is restricted to the valleys of southern Perú (Apurímac River Valley), *E. pendula* (Fig. 5d) occurs throughout the dry valleys from Northwestern Venezuela to southern Perú, where it co-occurs with *E. herrerae*.

Molecular data were available from only one individual of *E. herrerae*. Phylogenetic analysis of MYC showed that the haplotypes of *E. pendula* were not genealogically exclusive because a haplotype from Colombia was more closely related to the haplotype of *E. herrerae* (with moderate statistical support) than to other haplotypes of *E. pendula* (Fig. 1); this pair of haplotypes were sampled from localities separated by more than 2000 km. Phylogenetic analysis of NIA showed that two haplotypes of *E. pendula* were genealogically exclusive and concordant with geography (Fig 1); amplification and sequencing of NIA for the sample of *E. pendula* from Colombia was not possible (Zapata, 2013). The maximum level of sequence divergence between *E. pendula* (pendula3) and *E. herrerae* (herrerae) was 0.01 in MYC and 0.031 in NIA, sampled at approximately 1000 km apart (Fig. 1).

Morphological data showed that the samples of *E. herrerae* and *E. pendula* did not separate completely in morphological PCA space (Fig. 6b). The elevation plot was not bimodal (Fig. 7b), and thus it failed the first necessary condition to support the hypothesis of a morphological gap separating these species. No qualitative characters were fixed for alternative states in each species (Table 4, Appendix 2).

Bioclimatic data showed that the samples of these species overlapped markedly in bioclimatic PCA space (Fig. 9b). The PC scores did not differ significantly between this pair of species (Pillai’s trace = 0.087; *F*_2, 30_ = 1.438; *P* = 0.253), suggesting there is no evidence *E. herrerae* and *E. pendula* occur in significantly different environments; both species occur in areas with similar temperature and precipitation regimes (Table 5).

Both species overlapped completely in the range of flowering time (Fig. 10c, d).

<u>Clade D.</u> This clade was strongly supported by both loci and comprised *E. discolor, E. piurensis, E. paniculata, E. resinosa, E. reticulata* and *E. schreiteri* (Fig. 1), six species from the montane forests in the Tropical Andes (Table 1, Appendix 3). *E. discolor* (Fig. 5e) and *piurensis* (Fig. 5f) are narrowly restricted to central Colombia and northern Perú, respectively. *E. paniculat*a (Fig. 5g) is broadly distributed from Venezuela to central Bolivia. *E. resinosa* (Fig. 5h) is distributed from southern Ecuador to southern Bolivia. *E. reticulata* (Fig. 5i) is narrowly restricted to central Bolivia where it occurs in parapatry to *E. paniculata*. *E. schreiteri* (Fig. 5j) is distributed from central Bolivia to northern Argentina.

Two subclades were consistently recovered within clade D, one including *E. discolor, E. piurensis* and *E. resinosa* (hereafter clade D.A), and one including *E. paniculata* and *E. reticulata* (hereafter clade D.B); *E. schreiteri* was sister to D.B in the MYC tree and to D.A in the NIA tree (Fig. 1). Molecular data was available from only one individual of *E. piurensis.* Within clade D.A, phylogenetic analysis of MYC showed that the haplotypes of *E. discolor* and *E. resinosa* were not genealogically exclusive and instead they interdigitated forming a single conspecific lineage (along with *E. piurensis)* of unclear phylogenetic resolution (Fig. 1). Phylogenetic analysis of NIA showed that the haplotypes of *E. discolor* were genealogically exclusive and concordant with geography (Fig. 1). The haplotypes of *E. resinosa* were not genealogically exclusive because the haplotype of *E. piurensis* appeared to be related to *E. resionsa* (the basal relationship was a polytomy) creating a single conspecific lineage (Fig. 1); these haplotypes were sampled from localities separated by approximately 1000 km apart. The maximum level of sequence divergence was 0.01 for MYC between *E. resinosa* (resinosa1) and *E. piurensis* (piurensis) sampled at approximately 1000 km apart, and 0.026 for NIA between *E. resinosa* (resinosa2) and *E. discolor* (discolor1) sampled at approximately 3000 km apart (Fig. 1).

I used two morphological spaces to analyze morphological variation and weigh the strength of morphological data in support of the hypothesis of morphological gaps separating the species within clade D.A. In the first morphological PCA space, the samples of *E. discolor* and *E. schreiteri* did not overlap with the samples of *E. piurensis* and *E. resinosa* (Fig. 6c). The elevation plots of all pairwise comparisons in this morphological space were bimodal (Fig. 7c-g), and the corresponding proportions plots revealed that proportions ≤ 0.1 always overlapped, except for the pair *E. piurensis*-*E. discolor* (see below) (Fig. 8c-g). This implied that the frequency of intermediate phenotypes in all pairwise comparisons was ≤ 0.1, and thus there was enough evidence to suggest a morphological gap separating *E. schreiteri* from *E. discolor*, and both these species from *E. resinosa* and *E. piurensis* (except *E. piurensis* from *E. discolor,* see below). The hypothesis that these gaps represented species limits and not geographic variation within a single species was rejected only for the pair *E. discolor*-*E. schreiteri* (Appendix 4). The result obtained for *E. discolor*-*E. piurensis* (Fig. 8c) implied that the frequency of intermediate phenotypes between these species was > 0.1, and thus there was not enough evidence to suggest a morphological gap between them. No qualitative characters were fixed for alternative states in this pair of species (Table 4, Appendix 2).

In the second morphological PCA space, including only the samples of *E. resinosa* and *E. piurensis* that overlapped in the initial PCA (Fig. 6c), the samples of these species did not separate completely (Fig. 6d), and the elevation plot was not bimodal (Fig. 7h). Absence of glands in the pedicel was fixed in *E. resinosa* (Table 4, Appendix 2), however this difference was not statistically significant (*P* = 0.214).

Bioclimatic data revealed that samples of *E. discolor* separated almost completely from the samples of *E. piurensis*, *E. resinosa* and *E. schreiteri*, all of which overlapped markedly in bioclimatic PCA space (Fig. 9c). The PC scores differed significantly among these species (Pillai’s trace = 0.889; *F*_6_, 106 = 14.151; *P* < 0.000), with significant differences found along both PC axes (PC1: *F*_3_ = 30.047; *P* < 0.000; PC2: *F*_3_ = 6.199; *P* = 0.001). Tukey’s HSD revealed that the significant differences for PC1 were between all pairs (all *P* < 0.05), except *E. resinosa-E. schreiteri* (*P* = 0.99), and for PC2 between *E. schreiteri-E. piurensis* and *E. schreiteri-E. resinosa* (all *P* < 0.001). This suggests there is evidence all species occur in significantly different environments, with *E. schreiteri* found at relatively lower elevations (see also Appendix 3) in warmer places with strong seasonality, and *E. discolor* in places with overall the highest precipitation followed by *E. piurensis* and *E. resinosa* (Table 5).

*E. discolor*, *E. piurensis* and *E. schreiteri* did not overlap in their ranges of flowering time with each other (Fig. 10e, f, j); however all these species overlapped with the range of flowering time of *E. resinosa* (Fig. 10h)

Within clade D.B, phylogenetic analyses of MYC and NIA showed that the haplotypes of *E. reticulata* were genealogically exclusive and concordant with geography (Fig. 1). The haplotypes of *E. paniculata* formed a single non-exclusive lineage suggesting that *E. paniculata* is a plesiospecies with respect to *E. reticulata* (Fig. 1). MYC and NIA haplotypes of *E. schreiteri* were genealogically exclusive and concordant with geography with respect to any other species in clade D (Fig. 1). The maximum sequence divergence for both loci was 0.01 between *E. paniculata* (paniculata3) and *E. reticulata* (reticulata1) sampled at approximately 700 km apart (Fig. 1). Sequence divergence between *E. schreiteri* and any other species within clade D was always above 0.035 for either locus.

I used two morphological spaces to analyze morphological variation and evaluate morphological gaps for the species within clade D.B. In the first morphological PCA space, the samples of *E. schreiteri* did not overlap with the samples of *E. paniculata* and *E. reticulata* (Fig. 6e). The elevation plots of both pairwise comparisons in this morphological space were bimodal (Fig. 7i-j) and the corresponding proportion plots revealed that proportions ≤ 0.1 overlapped in both cases (Fig. 8i-j). This implied there was enough evidence to suggest a morphological gap separating *E. schreiteri* from both *E. paniculata* and *E. reticulata*, which represented species limits, rather than geographic variation within a single species (Appendix 4).

In the second morphological PCA space, the samples of *E. paniculata* and *E. reticulata* separated completely (Fig. 6f) and the elevation plot was bimodal (Fig. 7y). However, the proportion plot revealed that proportions > 0.9 overlapped (Fig. 8k), implying there was not enough evidence to suggest a morphological gap separating *E. paniculata* from *E. reticulata*. No qualitative characters were fixed for alternative states in either species (Table 4, Appendix 2).

Bioclimatic data revealed that samples of *E. paniculata* occupied a broad area in bioclimatic PCA space, and overlapped with samples of *E. schreiteri* and *E. reticulata*, both occupying a narrower area in bioclimatic PCA space (Fig. 9d). The PC scores differed significantly among these species (Pillai’s trace = 0.489; *F*_4, 162_ = 13.107; *P* < 0.000), with significant differences found along both PC axes (PC1: *F*_2_ = 15.320; *P* < 0.000; PC2: *F*_2_ = 11.062; *P* < 0.000). Tukey’s HSD revealed that the significant differences were between *E. schreiteri-E. paniculata*, and *E. schreiteri-E. reticulata* for PC1 (all *P* < 0.01), and between *E. reticulata-E. paniculata* for PC2 (*P* < 0.000). This suggests there is evidence these species occur in significantly different environments, with *E. schreiteri* found in areas with overall low precipitation, *E. reticulata* in areas with high temperatures and *E. paniculata* in areas with high precipitation (Table 5).

There was some overlap in the ranges of flowering times between all species because *E. paniculata* apparently flowers throughout the year (Fig. 10g, i, j).

<u>Clade F.</u> This clade included *E. alpina, E. callcottiae, E. florida, E. leucantha, E. myrtoidea, E. revoluta, E. rosea, E. rubra* and *E. serrata*, nine species restricted geographically to the Southern Temperate Andes in Chile and Argentina (Fig. 1). For the most part, these species co-occur in mosaic sympatry and some segregate according to habitat or elevation (Table 1, Appendix 3). *E. alpina* (Fig. 5k)*, E. rosea* (Fig. 5q) and *E. rubra* (Fig. 5r) are broadly distributed from central Chile (ca. 33° S) to Patagonia; *E. myrtoidea* (Fig. 5o) and *E. revoluta* (Fig. 5p) occur from central to southern Chile (ca. 40° S); *E. florida* (Fig. 5m) and *E. leucantha* (Fig. 5n) are restricted to southern Chile; *E. callcottiae* (Fig. 5l) is endemic to Juan Fernández Island; and *E. serrata* (Fig. 5s) is endemic to southern Patagonia.

As indicated above, two types of sequences were recovered for some species of this clade (clades F and F’ in Fig. 1; for details, see Zapata, 2013); therefore I evaluated species limits within each of these clades independently. In the NIA tree, the group *E. virgata + E. gayana* appeared closely related to clade F although with low statistical support; I discuss this species pair below because haplotypes of these species did not mix with the haplotypes of the species in clade F.

Molecular data were available from only one individual of *E. callcottiae.* Phylogenetic analysis of MYC showed that in clade F the haplotypes of all species were not genealogically exclusive; instead they formed a single conspecific lineage with some subclades weakly concordant with geography (Fig. 1). In clade F’, the haplotypes of *E. serrata* were genealogically exclusive and concordant with geography (Fig. 1). Three haplotypes of *E. rosea,* sampled from localities approximately 200 km apart, were genealogically exclusive and concordant with geography, although the statistical support for this clade was moderate. The maximum level of sequence divergence in MYC in clade F was 0.023 between *E. rubra* (rubra2) and *E. florida* (florida2) sampled at approximately 700 km apart, and in clade F’ it was 0.018 between *E. serrata* (serrata1) and *E. alpina* (alpina2) sampled at approximately 850 km apart. Phylogenetic analysis of NIA showed that in clade F the haplotypes of *E. serrata* were genealogically exclusive and concordant with geography (Fig. 1). The haplotypes of all other species were not genealogically exclusive, and instead they formed a single conspecific lineage with some subclades concordant with geography(Fig. 1). In clade F’, the haplotypes of *E. florida,* sampled from the only localities where this species is known to occur, were genealogically exclusive and concordant with geography (Fig. 1). The haplotypes of *E. rosea* were not genealogically exclusive because a haplotype of *E. rubra* (rubra1), sampled from the same locality as the haplotype rosea3 (of *E. rosea*), mixed with the other haplotypes of *E. rosea* (Fig. 1). The maximum level of sequence divergence in NIA in clade F was 0.038 between *E. rubra* (rubra2) and *E. alpina* (alpina2) sampled at approximately 2500 km apart, and in clade F’ it was 0.017 between *E. rosea* (rosea3) and *E. florida* (florida1), sampled at approximately 300 km apart (Fig. 1).

<u>*E. virgata* and *E. gayana*</u>. This pair of species occur in the Temperate Andes of Chile and Argentina, where they co-occur in mosaic sympatry with several species of clade F (Fig. 1). While *E. gayana* is narrowly restricted to few localities around latitude 39° S in Chile (Fig. 5a.8), *E. virgata* has a broader distribution southwards to Patagonia in Chile and Argentina (Fig. 5a.9).

Molecular data were available from only one individual of *E. gayana*. Phylogenetic analysis of MYC showed that the haplotypes of *E. virgata* were not exclusive and mixed with the haplotype of *E. gayana* forming a single conspecific lineage with unclear phylogenetic resolution (Fig. 1). Phylogenetic analysis of NIA showed that the haplotypes of *E. virgata* and *E. gayana* did not form a clade and were associated with clade F (with low statistical support), nonetheless the haplotypes of *E. virgata* were genealogically exclusive and concordant with geography (Fig. 1). The maximum level of sequence divergence between *E. virgata* and *E. gayana* was 0.013 in MYC and 0.054 in NIA (Fig. 1).

For morphological analyses of species within clade F, I included the samples of *E. virgata* and *E. gayana* as suggested by the phylogenetic analysis of NIA (analysis of *E. virgata* and *E. gayana* alone, as suggested by MYC, did not alter the results). I used five morphological spaces to analyze morphological variation and evaluate morphological gaps separating the species within clade F plus *E. virgata* and *E. gayana.*

In the first morphological PCA space, the samples of *E. serrata*, *E. virgata* and *E. gayana* clearly separated from the samples of all other species (Fig. 6g). The elevation plots of all pairwise comparisons in this morphological space were bimodal (Fig. 7l-a.11), and the corresponding proportion plots revealed that proportions ≤ 0.1 always overlapped, except for the pairs *E. serrata-E. gayana*, and *E. gayana-E. florida* (see below) (Fig. 8l-a.11). This implied there was enough evidence to suggest a morphological gap separating *E. serrata* from *E. virgata, E. virgata* from *E. gayana,* and these three species from all other species in clade F (except *E. serrata* from *E. gayana,* and *E. florida* from *E. gayana*). There was evidence to suggest these gaps represented species limits, except in the pairs *E. gayana*-*E. callcottiae* and *E. virgata-E. callcottiae* (Appendix 4). The result obtained for the pairs *E. serrata-E. gayana* and *E. florida-E. gayana* (Fig. 8t, a.6) implied that there was not enough evidence to suggest a morphological gap separating these species. Subshrub habit, absence of indumentum in the pedicel and ovary, and the presence of an elevated disk were fixed for *E. serrata;* however sampling was insufficient to support the hypothesis that any of these characters was truly fixed (*P* = 0.18). No qualitative characters were fixed between *E. florida* and *E. gayana* (Table 4, Appendix 2).

In the second morphological PCA space the samples of *E. florida* and *E. leucantha* separated from the samples of all other species (Fig. 6h). The elevation plots of all pairwise comparisons in this morphological space were bimodal (Fig. 7a.12-a.23). The corresponding proportion plots that revealed that only proportions ≤ 0.1 overlapped corresponded to the pairs *E. florida-E. alpina* (Fig. 8.a15), *E. florida-E. revoluta* (Fig. 8.a17), *E. florida-E. myrtoidea* (Fig. 8a.17), *E. florida-E. rubra* (Fig. 8a.18), *E. leucantha*-*E. alpina* (Fig. 8a.15), *E. leucantha*-*E. revoluta* (Fig. 8a.20) and *E. leucantha*-*E. rubra* (Fig. a.22). Thus, for these species there was enough evidence to suggest a morphological gap between them, which represented species limits rather geographic variation within a single species (Appendix 4). In all other pairwise comparisons in this morphological space, proportion plots revealed that proportions > 0.9 always overlapped (Fig. 8a.12, a.13, a.19, a.21, a.23), suggesting there was not enough evidence to suggest a morphological gap between any of the other species pairs. Only an elevated nectary disk was fixed in *E. rosea* for the pairwise comparison *E. florida-E. rosea* (Table 4, Appendix 2); however, sampling was insufficient to support this character as being truly fixed (*P* = 0.21). There were no fixed characters in all other pairwise comparisons within this morphological PCA space (Table 4, Appendix 2).

In the third morphological PCA space, the samples of all species overlapped (Fig. 6i). The elevation plots for the pairs *E. rosea-E. revoluta* (Fig. 7a.25)*, E. rosea-E. rubra* (Fig. 7a.27)*, E. rosea-E. callcottiae* (Fig. 7a.28) and *E. myrtoidea-E. revoluta* (Fig. 7a.33) were not bimodal. Although revolute margin of leaves was fixed in *E. revoluta,* and the presence of glands in the ovary was fixed in *E. rubra*, sampling was insufficient to suggest these characters were truly fixed (both *P* > 0.05). No qualitative characters were fixed between *E. rosea-E. callcottiae* (Table 4, Appendix 2). The elevations plots of the remaining pairwise comparisons were all bimodal (Fig. 7.a.24, a.26, a.29-a.32, a.34-a.38). The corresponding proportion plots revealed that proportions ≤ 0.1 overlapped only for *E. alpina-E. revoluta* (Fig. 8a.29) and *E. alpina-E. myrtoidea* (Fig. 8a.30), implying there was enough evidence to suggest a morphological gap separating these species, which represented species limits (Appendix 4). The proportion plots of the remaining pairwise comparisons revealed that proportions > 0.9 always overlapped (Fig. 8a. 24, a.26, a.31-a.32, a.34-a.38), implying there was not enough evidence to suggest a morphological gap separating any of these species pairs. There were no fixed characters in the pairwise comparisons within this morphological PCA space (Table 4, Appendix 2).

In the fourth morphological PCA, the samples of *E. serrata* and *E. virgata* clearly separated in morphological space (Fig. 6j). The elevation plot was bimodal (Fig. 7a.39), and the proportion plot revealed that proportions ≤ 0.1 overlapped (Fig. 8a.39), implying there was enough evidence to suggest a morphological gap separating *E. serrata* from *E. virgata.* This gap represented a species limits rather than geographic variation in a single species. (Appendix 4).

In the fifth morphological PCA space, the samples of *E. florida* and *E. leucantha* clearly separated in morphological space (Fig. 6k). The elevation plot was bimodal (Fig. 7a.40), and the proportion plot revealed that proportions > 0.9 overlapped (Fig. 8a40), implying there was not enough evidence to suggest a morphological gap separating *E. florida* from *E. leucantha*. Elevated nectary disk was fixed in *E. leucantha*, but sampling was insufficient to suggest this character was truly fixed (*P* = 0.11) (Table 4, Appendix 2).

Bioclimatic data revealed a high degree of overlap among the samples of all species (Fig. 9e). The PC scores differed significantly among these species (Pillai’s trace = 0.851; *F*_20, 404_ = 14.968; *P* < 0.000), with significant differences found along both PC axes (PC1: *F*_10_ = 11.172; *P* < 0.000; PC2: *F*_10_ = 19.81; *P* < 0.000). Tukey’s HSD revealed that the significant differences for PC1 were between *E. serrata* and all other species, and between *E. myrtoidea*/*E. revoluta* and *E. leucantha*, *E. rosea, E.virgata, E. rubra,* and *E. alpina* (all *P* < 0.001). For PC2 differences were between *E. alpina* and all species (all *P* < 0.000), except *E. serrata* and *E. virgata* (all *P* > 0.1); *E. serrata* and all species (all *P* < 0.000), except *E. virgata, E. rosea*, and *E. myrtoidea* (all *P* > 0.1); *E. virgata* and all species (all *P* < 0.000), except *E. rosea, E. myrtoidea*, and *E. florida* (all *P* > 0.1); *E. rosea* and *E. rubra, E. revoluta, E. leucantha*, and *E. callcottiae* (all *P* < 0.02); and *E. callcottiae*-*E. rubra,* and *E. callcottiae*-*E. florida* (all *P* < 0.01). This suggests there is evidence these species occur in significantly different environments, with *E. serrata* found in areas with overall higher precipitation, but low seasonality in precipitation, *E. myrtoidea* and *E. revoluta* found in areas with the highest temperatures and low precipitation, *E. alpin*a found in highland areas with high seasonality, and *E. virgata, E. rosea* and *E. callcottiae* found in relatively warm areas with low precipitation (Table 5).

All species overlapped markedly in the range of flowering time, which is strongly associated with seasonality in the southern hemisphere (Fig. 10k-s).

<u>Clade G.</u> This clade included six species distributed in southeastern Brazil-northeastern Argentina (*E. bifida, E. farinacea, E. laevis, E. ledifolia, E. megapotamica, E. petrophila*), four in the Andes (*E. angustifolia, E. hypoglauca, E. illinita, E. tucumanensis*) and one endemic to the Sierra de Córdoba in central Argentina (*E. cordobensis*). The species from southeastern Brazil-northeastern Argentina co-occur in mosaic sympatry, and most of them segregate along habitat and elevational gradients (Table 1, Appendix 3). *E. bifida* (Fig. 5u) and *E. megapotamica* (Fig. 5a.2) have the broadest geographic ranges followed by *E. farinacea* (Fig. 5w), *E. laevis* (Fig. 5z), and the narrowly restricted *E. ledifolia* (Fig. 5a.1) and *E. petrophila* (Fig. 5a.3). In the Andean group, *E. angustifolia* (Fig. 5t) has a broad geographic distribution with populations in southern Perú, north of the Atacama desert, to central Chile, south of the desert; *E. illinita* (Fig. 5y) is narrowly distributed in central Chile, south of the Atacama desert. In central Chile, these two species co-occur in mosaic sympatry with other species of clade F (Fig. 1). In the Tropical Andes, *E. hypoglauca* (Fig. 5x) occurs from central Bolivia to northwestern Argentina where it is parapatric to *E. tucumanensis* (Fig. 5a.4), which extends southwards to the state of Tucumán. These two species co-occur in mosaic sympatry with species of clades A, D, and H (Fig. 1). *E. cordobensis* (Fig. 5v) is the only species that occur in the Sierra de Córdoba.

Molecular data were available from only one individual of *E. tucumanensis*. Phylogenetic analyses of MYC showed that the haplotypes of *E. bifida*, *E. farinacea* and *E. petrophila* were genealogically exclusive and concordant with geography (Fig. 1). The haplotypes of all other species were not genealogically exclusive and formed a single conspecific lineage with few subclades concordant with geography (Fig. 1). Phylogenetic analysis of NIA showed that the haplotypes of *E. farinacea* and *E. petrophila* were genealogically exclusive and concordant with geography (Fig. 1). The haplotypes of *E. laevis* formed a single non-exclusive lineage concordant with geography, suggesting that *E. laevis* is a plesiospecies with respect to *E. petrophila* (Fig. 1). The haplotypes of all other species were not genealogically exclusive and formed a single conspecific lineage with several subclades concordant with broad geographic regions, e.g., all species from the Tropical Andes and Sierra de Córdoba grouped together (Fig. 1). The maximum level of sequence divergence in MYC was 0.031 between *E. angustifolia* (angustifolia1) and *E. bifida* (bifida2) sampled approximately 3000 km apart, and 0.047 in NIA between *E. illinita* (illinita2) and *E. megapotamica* (megapotamica3) sampled approximately 2000 km apart (Fig. 1).

I used seven morphological spaces to analyze morphological variation and evaluate morphological gaps separating the species within clade G. There were only two specimens of *E. ledifolia* and *E. petrophila* available; with this sampling, the method of Zapata and Jiménez (2012) cannot be used, therefore these data were not included in these analyses. In the first morphological space, the samples of *E. angustifolia*, *E. bifida* and *E. megapotamica* clearly separated from the samples of all other species (Fig. 6l). The elevation plots of all pairwise comparisons in this morphological space were bimodal (Fig. 7a.41-a.60). The corresponding proportion plots revealed that proportions > 0.9 overlapped only for the pairs *E. angustifolia-E. illinta* (Fig. 8a.41), *E. angustifolia-E. laevis* (Fig. 8a.42), *E. angustifolia-E. cordobensis* (Fig. 8a. 43) and *E. angustifolia-E. farinacea* (Fig. 8a.47). This implied there was not enough evidence to suggest a morphological gap separating these species. Although revolute margin of leaves was fixed in *E. illinita*, and presence of glands in the adaxial lamina was fixed in *E. angustifolia* (Table 4, Appendix 2), sampling was insufficient to support these characters were truly fixed (all P > 0.05). The proportion plots of the remaining pairwise comparisons revealed that proportions ≤ 0.1 always overlapped (Fig. 8a.44-a.46, a.48-a.60), implying there was enough evidence to suggest a morphological gap separating all these species. The hypothesis that these gaps represented species limits and not geographic variation within a single species was not rejected, expect for the pairs *E. cordobensis-E. megapotamica* and *E. hypoglauca-E. megapotamica* (Appendix 4).

In the second morphological PCA space, the samples of *E. farinacea* clearly separated from the samples of all other species (Fig. 6m). The elevation plots of all pairwise comparisons in this morphological space were bimodal (Fig. 7a.61-a.65), and the corresponding proportion plots revealed that proportions ≤ 0.1 always overlapped (Fig. 8a.61-a65), implying there was enough evidence to suggest a morphological gap separating *E. farinacea* from all the other species. There was not enough evidence to suggest these gaps represented species limits in any pairwise comparison, except for the pair *E. farinacea*-*E. laevis* (Appendix 4).

In the third morphological PCA space, the samples of *E. hypoglauca and E. ledifolia* together separated from the samples of *E. cordobensis*, *E. illinita* and *E. tucumanensis* (Fig. 6m). The elevation plots of all pairwise comparison in this morphological space were bimodal (Fig. 7a.66-a.71), and t he corresponding proportion plots revealed that proportions > 0.9 overlapped for all pairs (Fig. 8a.67-a.71), except for the pair *E. illinita-E. laevis* (see below). This implied there was not enough evidence to suggest a morphological gap separating these species. Revolute margins of leaves and elevated nectary disk were fixed for *E. illinita*, while indumentum in ovary was fixed for *E. hypoglauca* (Table 4, Appendix 2), however sampling was insufficient to suggest these characters were truly fixed (all *P* > 0.05). In the *E. illinta-E. laevis* comparison (Fig. 8a.66) the frequency of intermediate phenotypes was ≤ 0.1, hence there was enough evidence suggesting a morphological gap separating these species. The hypothesis that this gap represented a species limit and not geographic variation within a single species was rejected (Appendix 4).

In the fourth morphological PCA, the samples of *E. illinita* separated from the samples of *E. cordobensis* and *E. tucumanensis* (Fig. 6o). The elevation plots both pairwise comparisons in this morphological space were bimodal (Fig. 7a.72, a.73), and the corresponding proportion plots revealed that proportions ≤ 0.1 overlapped in both cases (Fig. 8a.72-a.73). This implied there was enough evidence to suggest a morphological gap separating *E. illinta* from *E. cordobensis* and *E. tucumanensis*, but the evidence was not enough to suggest this gap represented a species limit for the pair *E. illinta*-*E. cordobensis* (Appendix 4).

In the fifth morphological PCA space, the samples of *E. cordobensis* and *E. tucumanensis* did not separate completely in morphological space (Fig. 6p). The elevation plot was bimodal (Fig. 7a.74), and the proportion plot revealed that proportions > 0.9 overlapped (Fig. 8a.74), implying there was not enough evidence to suggest a morphological gap between *E. cordobensis* and *E. tucumanensis.* No qualitative characters were fixed for alternative states in each species (Table 4, Appendix 2).

In the sixth morphological PCA space, the samples of *E. hypoglauca* and *E. laevis* did not separate completely (Fig. 6q), and the elevation plot was not bimodal (Fig. 7a.75). No qualitative characters were fixed for alternative states in each species (Table 4, Appendix 2).

In the seventh morphological PCA space, the samples of *E. angustifolia* and *E. bifida* clearly separated in morphological space (Fig. 6.r). The elevation plot was bimodal (Fig. 7a.76), and the proportion plot revealed that proportions ≤ 0.1 overlapped (Fig. 8a.76), implying there was enough evidence to suggest a morphological gap between *E. angustifolia* and *E. bifida.* The hypothesis that this gap represented a species limit and not geographic variation within a single species was not rejected (Appendix 4).

Bioclimatic data revealed that the group of samples of species from the Andes-Sierra de Córdoba did not overlap with the group of samples of species from southeastern Brazil-northeastern Argentina in bioclimatic PCA space (Fig. 9f). The PC scores differed significantly among all the species (Pillai’s trace = 1. 419; *F*_20, 362_ = 44.174; *P* < 0.000), with significant differences found along both PC axes (PC1: *F*_10_ = 60.84; *P* < 0.000; PC2: *F*_2_ = 33.318; *P* < 0.000). Tukey’s HSD revealed that the significant differences for PC1 were between the group of species from the Andes-Sierra de Córdoba and the group of species from southeastern Brazil-northeastern Argentina (all *P* < 0.000), except *E. tucumanensis-E. laevis* and *E. tucumanensis-E. ledifolia* (*P* > 0.09). This suggests the group of species from the Andes-Sierra de Córdoba occur at higher elevations (see also Appendix 3) in areas with higher seasonality in precipitation, while the group of species from southeastern Brazil-northeastern Argentina occur at considerably lower elevations in warmer areas with less precipitation seasonality (Table 5). Within the group of species from the Andes-Sierra de Córdoba, there were significant differences between all species (all *P* < 0.001), except for the pairs *E. hypoglauca*-*E. angustifolia*, *E. cordobensis*-*E. illinita*; *E. cordobensis*-*E. illinita, E. cordobensis*-*E. tucumanensis*; and *E. tucumanensis*-*E. illinita* (all *P* > 0.06). Within the group of species from southeastern Brazil-northeastern Argentina, only the pairs *E. laevis-E. bifida*, *E. laevis-E. farinacea*, and *E. laevis-E. megapotamica* showed statistically significant differences (all *P* < 0.001). Tukey’s HSD revealed that the significant differences for PC2 were between *E. laevis* and all other species (all *P* < 0.000), except the pairs *E. laevis-E. ledifolia* and *E. laveis-E. petrophila (*all *P* > 0.06); *E. farinacea* and *E. megapotamica, E. illinita, E. cordobensis*, and *E. tucumanensis* (all, *P* < 0.000); *E. bifida* and *E. angustifolia, E. megapotamica, E. illinita, E. cordobensis*, and *E. tucumanensis* (all, *P* < 0.000); and *E. hypoglauca* and *E. tucumanensis* (*P* < 0.000). Together, these results suggest there is evidence several species in clade G occur in significantly different environments, with most species from the Andes-Sierra de Córdoba occurring in areas with a gradient from high to low seasonality in precipitation, and from low to high temperatures in the coldest months, while most species from southeastern Brazil-northeastern Argentina occur in areas with a gradient from low to high seasonality in temperature, and from low to high overall precipitation (Table 5).

All species overlapped markedly in the range of flowering time, which is likely associated with seasonality in the southern hemisphere (Fig. 10k-s).

<u>Clade H.</u> This clade included *E. myrtilloides* and *E. polifolia* (Fig. 1), two species distributed in the highest peaks of the Tropical Andes and Costa Rica (Table 1, Appendix 3). While *E. polifolia* (Fig. 5a.5) is narrowly distributed in northern Perú (Chachapoyas Mountains), *E. myrtilloides* (Fig. 5a.6) is broadly distributed from Costa Rica to southern Bolivia.

Phylogenetic analyses of MYC and NIA showed that the haplotypes of both species were not genealogically exclusive; instead they interdigitated forming a single conspecific lineage with unclear phylogenetic resolution (Fig. 1). Two NIA haplotypes of *E. myrtilloides,* sampled from two localities separated by approximately 50 km in Costa Rica (Fig. 1 myrtilloides1, myrtilloides7), were genealogically exclusive and concordant with geography (Fig. 1). The maximum level of sequence divergence for MYC was 0.01 between polifolia1 and myrtilloides6 separated by approximately 1600 km, and for NIA was 0.04 between myrtilloides7 and myrtilloides6, which were collected approximately 3500 km apart (Fig. 1).

Morphological data showed that the samples of *E. myrtilloides* and *E. polifolia* clearly separated in morphological PCA space (Fig. 6s). The elevation plot was bimodal (Fig. 7a.77), however, the proportion plot revealed that proportions > 0.9 overlapped (Fig. 8a.77), implying there was not enough evidence to suggest a morphological gap separating *E. myrtilloides* from *E. polifolia*. Revolute leaf margins were fixed (*P* = 0.032) in *E. polifolia* (Table 4, Appendix 2). This implied that the frequency of non-revolute leaves in *E. polifolia* occurs at a frequency ≤ 0.1, and thus there was enough evidence to suggest a morphological gap in qualitative characters between *E. myrtilloides* and *E. polifolia.*

Bioclimatic data revealed that the samples of *E. myrtilloides* and *E. polifolia* overlapped completely in bioclimatic space (Fig. 9g). The PC scores did not differ significantly between these species (Pillai’s trace = 0.025; *F*_2, 61_ = 0.79; *P* = 0.46), suggesting there is no evidence *E. myrtilloides* and *E. polifolia* occur in significantly different environments; both species occur in areas with similar temperature and precipitation regimes (Table 5).

Both species overlapped in the range of flowering time because *E. myrtilloides* apparently flowers throughout the year (Fig. 10g, i, j).

*E. pulverulenta*. The distinctness of this species was strongly supported by both loci (Fig. 1). It has a broad distribution in central Chile (Fig. 5a.7) where it co-occurs in mosaic sympatry with several species of clade F (Fig. 1).

Phylogenetic analyses of MYC and NIA showed that the haplotypes of *E. pulverulenta* were genealogically exclusive and concordant with geography (Fig. 1). Although the phylogenetic position of *E. pulverulenta* was discordant between gene trees, the haplotypes of *E. pulverulenta* were always more closely related to each other than to the haplotypes of any other species. The maximum sequence divergence in MYC was 0.007, and in NIA was 0.02 between the specimens collected at approximately 700 km apart (Fig. 1).

Since *E. pulverulenta* showed conflicting relationships with clades A and B, I analyzed morphological variation in this species in the context of both these clades. The samples of *E. pulverulanta* separated completely from the samples of the species in clades A and B in morphological PCA space (Fig. 6t). The elevation plots of all pairwise comparisons were bimodal (Fig. 7a.78-a.81), and the corresponding proportion plots revealed that proportions ≤ 0.1 always overlapped (Fig. 8a.78-a.81). This implied that the frequency of intermediate phenotypes occurred at a frequency ≤ 0.1, and thus there was enough evidence to suggest a morphological gap separating *E. pulverulenta* from the species in clades A and B. The hypothesis that these gaps represented species limits and not geographic variation within a single species was rejected for all pairwise comparisons, except for the pair *E. pulverulenta-E. millegrana* (Appendix 4).

Bioclimatic data revealed that samples of *E. pulverulenta* separated almost completely from the samples of species in clades A and B (Fig. 9h). The PC scores differed significantly among these species (Pillai’s trace = 1.150; *F*_8, 152_ = 25.755; *P* < 0.000), with significant differences found along both PC axes (PC1: *F*_4_ = 70.58; *P* < 0.001; PC2: *F*_4_ = 10.829; *P* < 0.001). Tukey’s HSD (focusing only on *E. pulverulenta* differences) revealed that the significant differences were between *E. pulverulenta* and all other species for PC1 (all *P* < 0.000), and between *E. pulverulenta* and *E. millegrana* for PC2 (*P* < 0.000). This suggests there is evidence *E. pulverulenta* and species in clades A and B occur in significantly different environments, with *E. pulverulenta* found in areas of strong temperature and precipitation seasonality as compared to the other species (Table 5).

### Summary of results

Table 6 summarizes the results of the analyses of molecular data to evaluate the operational species criterion of genealogical exclusivity of alleles. Table 7 summarizes the results of the analyses of morphological variation to evaluate operational species criterion of morphological gaps and its relationship with geography. Table 8 summarizes the results of the analyses of bioclimatic data to evaluate operational species criterion of differences in realized ecological niche.

**TABLE 6.**
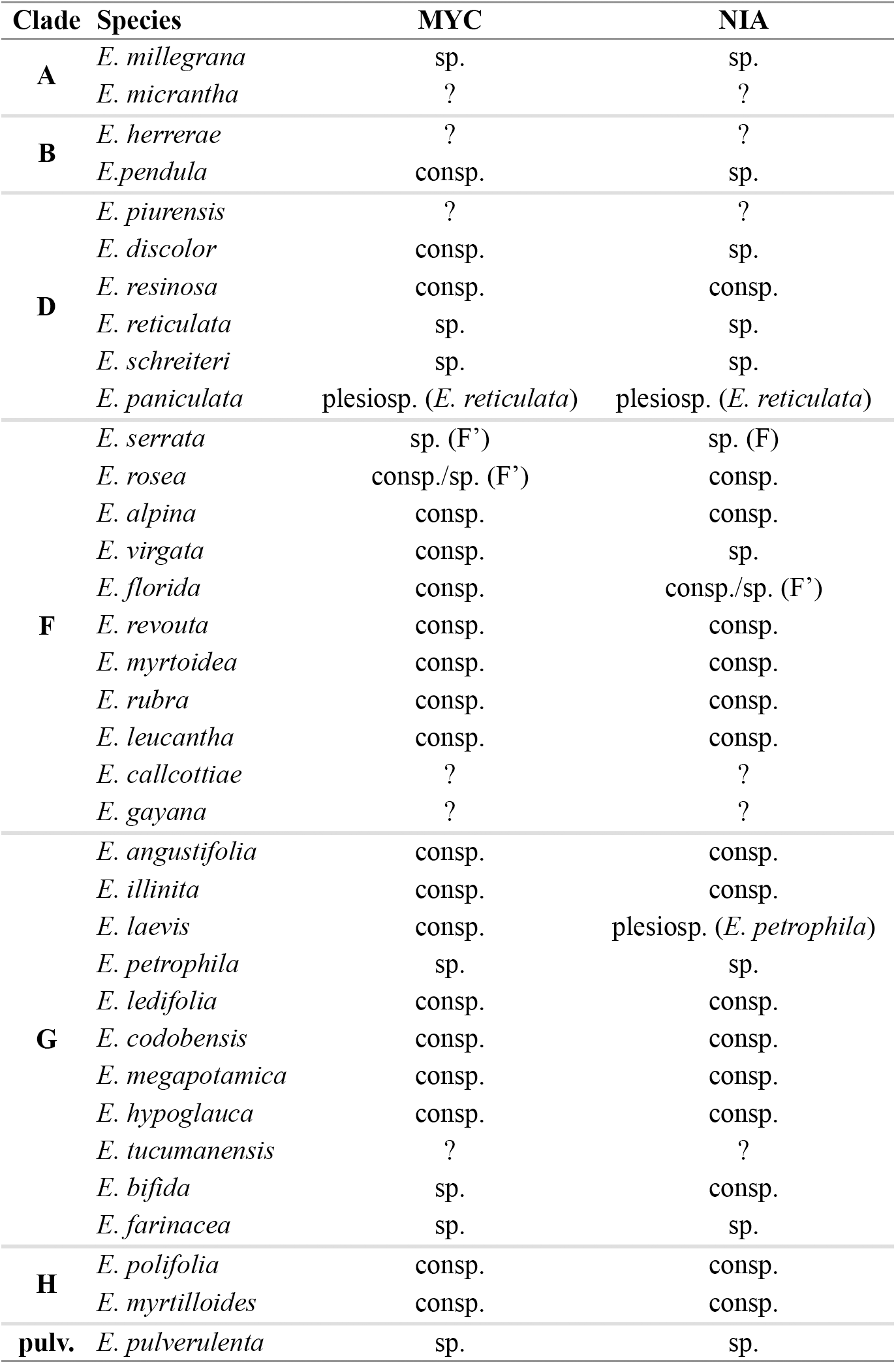
Summary of the results of the analysis of genealogical exclusivity for each locus. For species in clade F, the analysis was run for clades F and F’ independently. ?: not enough samples to assess exclusivity; sp.: one species; consp.: one conspecific lineage; plesiosp.: plesiospecies (relative to *species*). See Wiens and Servedio (2002) for details.

**TABLE 7.**
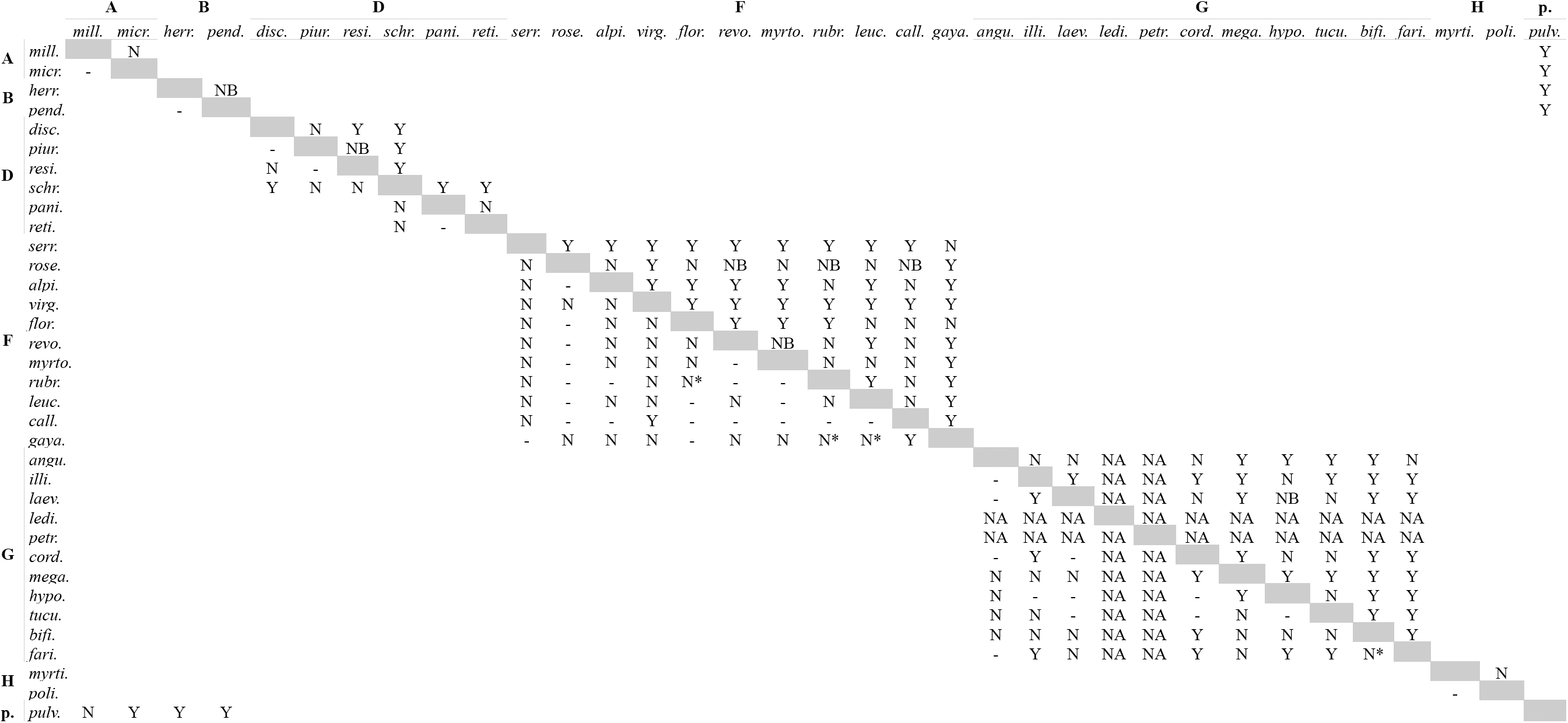
Analysis of morphological gaps (Zapata and Jiménez 2010). Above diagonal, comparisons indicate whether there was a gap (Y), no gap (N) or elevation plot was not bimodal (NB). Below diagonal, comparisons indicate whether the hypothesis that the gap represented a species limit rather could be rejected (Y) or not rejected (N). Comparisons that did not require detrending with asterisk. NA: not enough data. For clades names see Fig. 1.

## DISCUSSION

Here, I examined the geographic patterns of variation in two nuclear neutral loci, 40 morphological characters, and 19 bioclimatic variables to weigh the strength of this evidence to support 35 hypotheses of species boundaries within the genus *Escallonia* (Sleumer, 1968) by evaluating three operational species criteria (Sites and Marshall, 2003, 2004). Overall, my results showed that molecular data provided the weakest support to these hypotheses because the haplotypes of most species were more closely related to haplotypes of other species than to haplotypes of the same species (Fig. 1; Table 6). Conversely, morphological and bioclimatic data provided strong support to these hypotheses of species boundaries (Tables 7, 8). Several species were separated by morphological gaps (Figs. 7-8), and for the most part there was enough evidence to suggest these gaps represented species boundaries and not morphological differentiation within a single species (Table 7). Likewise, bioclimatic data provided support to these hypotheses because most species differed in their realized ecological niche (Fig. 9, Table 8). However, not all data sets provided support to all or the same hypotheses of species boundaries (Tables 6-8). I recognize species as independently evolving segments of population-level lineages (de Queiroz, 2005, 2007). Under this species concept, evidence from multiple operational species criteria provides support to a hypothesis of a species boundary but no single criterion is necessary to do so. That is, evidence from any one or more criteria provides support to the hypothesis of a species boundary, but the absence of evidence from one or more criteria does not constitute evidence contradicting such hypothesis (de Queiroz, 2007; see also Gotelli and Ellison, 2004). Thus, for instance, a hypothesis of a species boundary that is not supported by the analysis of morphological discontinuities can, nonetheless, correspond to a species boundary supported by molecular and/or ecological differences (for details, see de Quieroz, 2005, 2007).

**TABLE 8.**
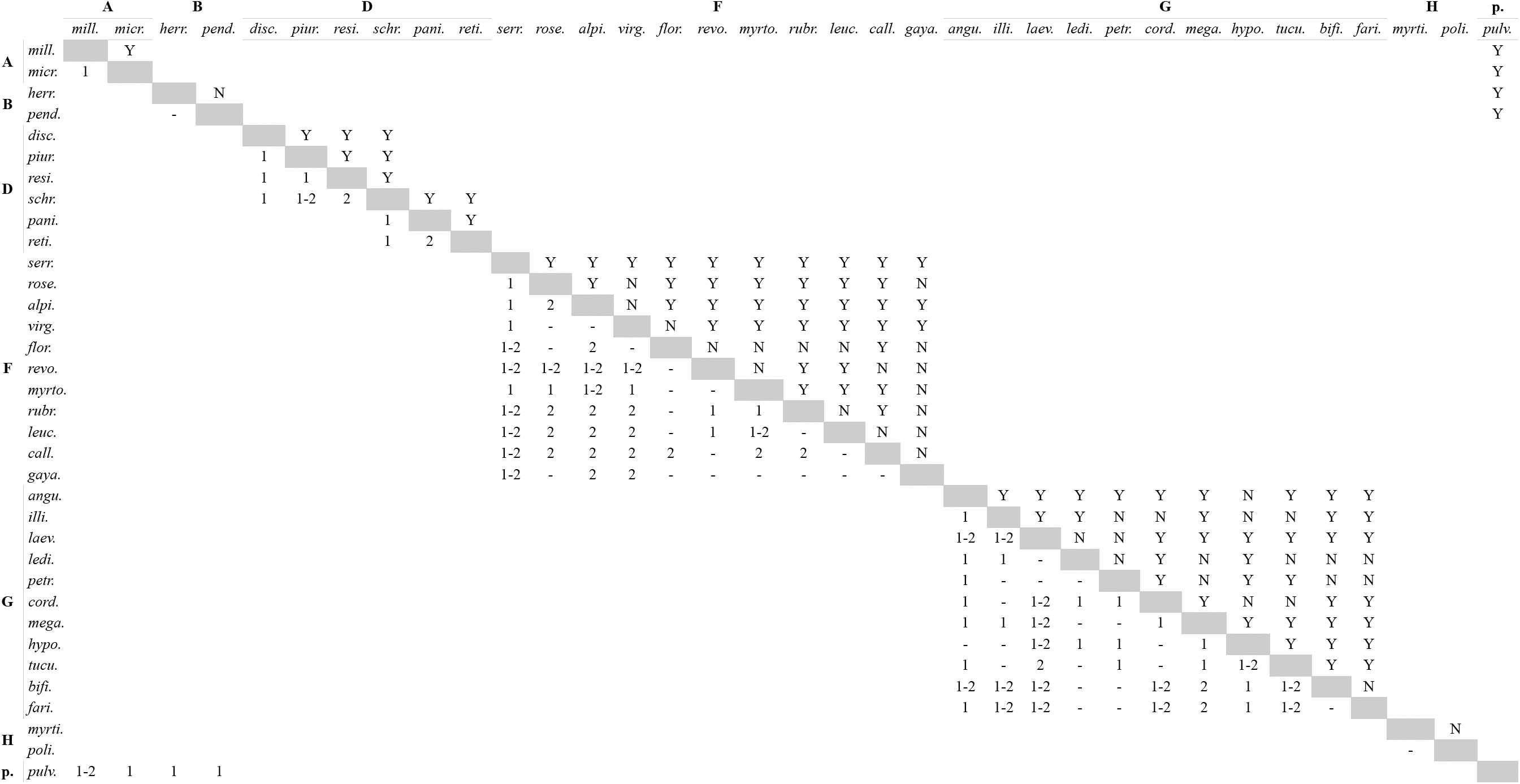
Analysis of differences in bioclimatic niche using PCA and MANOVA. Above diagonal, comparisons indicate whether there was a significant difference in central tendency (Y), or not (N). Below diagonal, comparisons indicate on which axis of the PCA was the difference statistically significant, 1: PC1, 2: PC2, 1-2: PC1 and PC2. For clades names see Fig. 1.

With this framework in mind, I summarize the results of the evaluation of the three operational species criteria that I examined in Fig. 11. This figure resembles a “crossing polygon” (Clausen et al., 1941) with species arranged around the periphery of a polygon connected with lines when both morphological and bioclimatic evidence failed to meet the operational species criteria I evaluated for these data. Consequently, species without connections represent cases in which morphological or bioclimatic data (or both) met the operational criteria, and thus there was enough evidence to support the hypothesis of a species boundary. I incorporated molecular evidence using circles to indicate when a species was monophyletic (continuous line) or paraphyletic (dashed lines). Below, I discuss the strength of the data to support the hypotheses of species boundaries within each clade in turn.

**Fig. 11.**
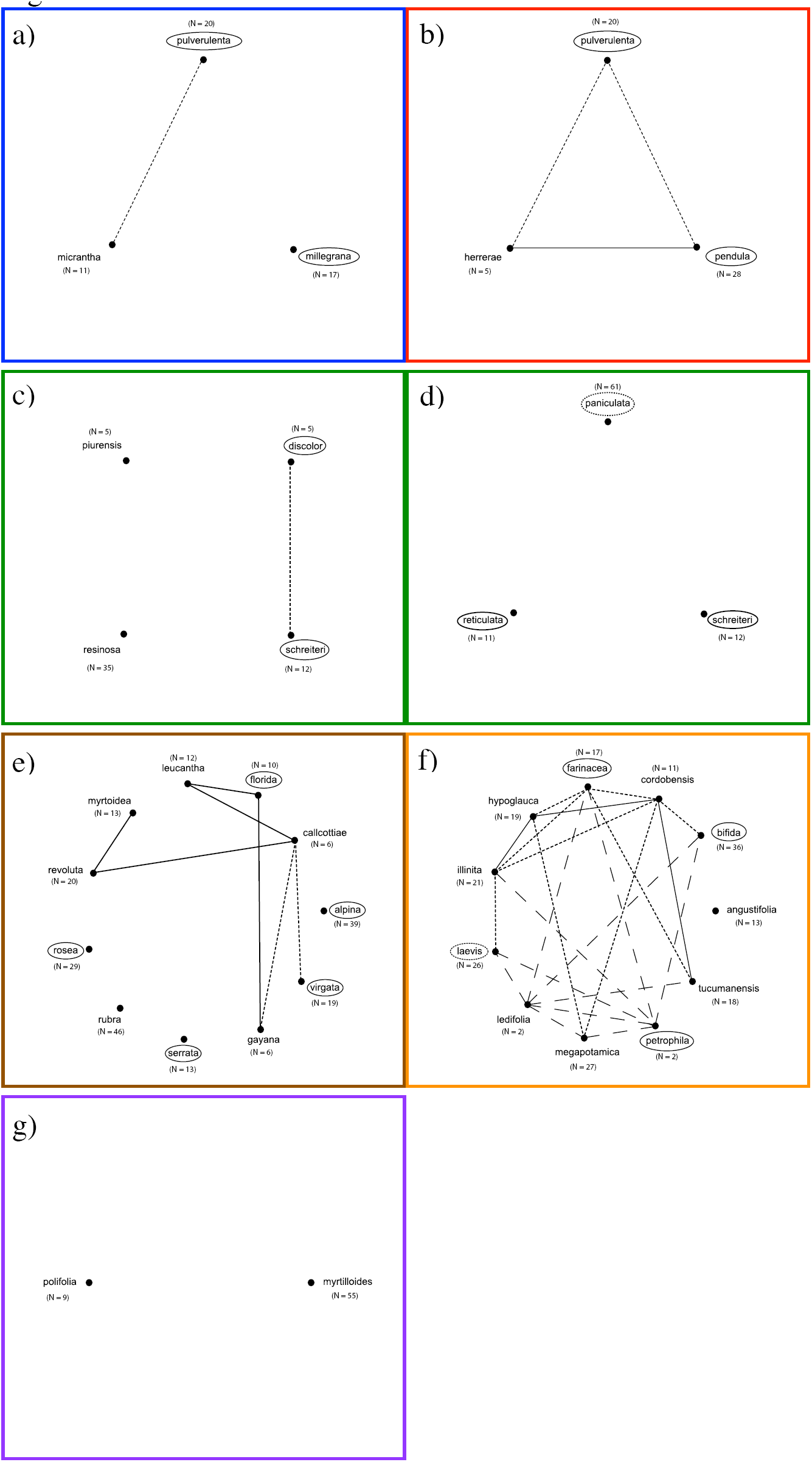
“Species polygons” (cf. Clausen et al., 1940) summarizing results of the evaluation of three operational species criteria to evaluate hypotheses of species boundaries within clades. Solid lines connect species when both morphological and bioclimatic evidence failed to meet the operational species criteria of morphological gaps and significant differences in realized ecological niche, respectively. Short-dashed lines connect species when the morphological gaps could be explained as geographic variation within a single species. Long-dashed lines connect species when there were no bioclimatic differences and this was the only operational species criterion evaluated because the method of Zapata and Jiménez (2012) cannot be used with sample size = 2 (see text for details). Circles around species names indicate whether species are monophyletic (continuous) or paraphyletic (dotted). N = sample size used in morphological and bioclimatic analyses. a) clade A plus *E. pulverulenta*; b) clade B plus *E. pulverulenta*; c) clade D.A; d) clade D.B; e) clade F; f) clade G; g) clade H. Colored boxes around each panel according to the color of the clade the species belongs to (for clade color, see Fig. 1). Note that *E. pulverulenta* is shown with clades A and B. For details, see text.

<u>Clade A.</u> Molecular and bioclimatic data supported the hypothesis of a species boundary between *E. millegrana* and *E. micrantha* (Fig. 11a). Haplotypes of *E. millegrana* were genealogically exclusive and concordant with geography (Table 6), and both species showed differences in their realized ecological niche (Table 8). Morphological similarity may be attributable to an instance of allopatric speciation, whereby the fragmentation of a species’ ancestral geographic range occurred when climatic change in the geographic space between diverging populations occurred more rapidly than the rate at which morphological (and maybe anatomical, see Stern, 1972) adaptations may have evolved in each population (Wiens, 2004; Kozak and Wiens, 2006).

<u>Clade B.</u> Molecular evidence from one locus provided support to the hypothesis of a species boundary between *E. herrerae* and *E. pendula* (Fig. 11a) because haplotypes of *E. pendula* were genealogically exclusive (Table 6). These haplotypes, however, were sampled in close geographic proximity (ca. 100 km apart) and covered a small portion of the geographic range of this species (Fig. 5d). Although morphological and bioclimatic data did not support the hypothesis of a species boundary (Table 7, 8), sample size was too small for *E. herrerae* raising the possibility of lack of statistical power to detect morphological discontinuities and/or bioclimatic differences (Stockman and Bond, 2007; Zapata and Jiménez, 2012). Further geographic sampling is necessary to evaluate this species boundary with increasing rigor.

<u>Clade D.A.</u> The data supported all the hypotheses of species limits among *E. discolor, E. piurensis, E. resinosa,* and *E. schreiteri* (Fig. 11c). The species boundary between *E. schreiteri* and all other species received support from all data sets, while the hypotheses of species boundaries among the other species received support from either molecular, morphological or bioclimatic data (Tables 6-8). This means that some species were genealogically exclusive, or were morphologically distinct, or displayed differences in their bioclimatic niche, or showed different combinations of these properties. Although there was evidence to suggest that the morphological gap between *E. schreiteri* and *E. discolor* could be explained by geography alone (morphological variation within a single species), this seems unlikely because *E. resinosa* and *E. piurensis*, which occur at intermediate geographic localities between *E. discolor* and *E. resinosa*, were separated by a species boundary from *E. schreiteri*. *E. discolor* is isolated geographically from the other species in this clade suggesting a likely case of allopatric speciation (see also clade A). In general, sample sizes were too small for *E. discolor* and *E. piurensis*, and further sampling is desirable to assess these species boundaries more thoroughly.

<u>Clade D.B.</u> All lines of evidence provided support to the hypotheses of species boundaries among *E. paniculata, E. reticulata,* and *E. schreiteri* (Fig, 11d). As in clade D.A., the hypothesis of the species boundary between *E. schreiteri* and other species was supported by all data sets, while the species boundary between *E. reticulata* and *E. paniculata* was supported only by molecular and bioclimatic data (Table 6-8). That *E. paniculata* was paraphyletic with respect to the monophyletic *E. reticulata* likely reflects the budding nature of an incipient speciation event (parapatric speciation) given the large geographic range (and likely large population size) of *E. paniculata* (Rieseberg and Brouillet, 1994). It is noteworthy that this species boundary corresponded to differences in bioclimatic conditions (Table 8), suggesting the possibility of ecological speciation along an environmental gradient (Nosil et al., 2009) that needs to be studied in closer detail.

<u>Clade F.</u> Molecular, morphological and/or bioclimatic data supported most hypotheses of species boundaries within this clade (Fig. 11e). Although none of these hypotheses was supported concurrently by all data sets, the data supported most hypotheses with evidence from at least one operational species criterion (Tables 7-9). This means that some species were only genealogically exclusive, others were only morphologically distinct, others differed only in their realized ecological niche, and others showed different combinations of these properties. It is worth noting that bioclimatic differences overlaid what are effectively species with sympatric and parapatric distributions (Fig. 5) and no differentiation in flowering time (Fig. 10). This is consistent with the intriguing possibility that within clade F environmentally-mediated selection maybe an important evolutionary force driving speciation (or at least maintaining species differences), perhaps reinforced by the positive effect of interspecific gene flow on genetic variation and adaptation (Rieseberg et al., 2003; Grant and Grant, 2008). The evidence for rejecting the hypotheses that there are species boundaries between *E. callcottiae* and *E. gayana* and other species (Fig. 11e) was weak and may be attributable to lack of statistical power to detect significant results (Stockman and Bond, 2007; Zapata and Jiménez, 2012); exhaustive geographic sampling is necessary before these hypotheses can be rejected confidently. Beyond issues of statistical power, that the species boundary between *E. myrtoidea* and *E. revoluta* was weakly supported is noteworthy because these species occur at different elevations (Appendix 3) and microhabitats (Table 1). It is likely that the fine microclimatic differences of the habitats where these species occur was not captured by the broad scale trends in bioclimatic variation I used here (Hijmans et al., 2005).

<u>Clade G.</u> As for clade F, molecular, morphological and/or bioclimatic data supported several hypotheses of species boundaries within this clade (Fig. 11f). Most hypotheses received support from at least one operational species criterion (Tables 6-8), implying that species were genealogically exclusive, or morphologically distinct, or differed in their bioclimatic niche, or showed different combinations of these properties. The hypothesis that *E. petrophila* was a distinct species was supported by molecular data. However, both this species and *E. ledifolia* are poorly sampled, thus a critical evaluation of the operational criterion of morphological gaps was not possible (Table 7), and the evaluation of the operational criterion of differences in realized ecological niche was compromised likely by lack of sampling (Fig. 11f); more samples are necessary to evaluate rigorously these species boundaries. The possibility that several morphological gaps between pairs of allopatric species (i.e, species from Brazil and species from the Andes) could be explained as geographic variation within a single species is intriguing (Fig. 11f). However, given that morphological gaps between these same pairs of allopatric species and other species at geographically intermediate localities (sympatric or parapatric) represented species boundaries, this possibility seems unlikely. Nonetheless, examining this complex pattern of morphological variation in the light of a better resolved molecular phylogeny will help to better understand how morphology is evolving within clade G. *E. cordobensis* is poorly sampled and this may explain the lack of statistical power to detect gaps in morphology between this and other species (Table 7; Fig. 11f). This statistical issue aside, it is puzzling that four species with allopatric (*E. illinita, E. cordobensis, E. hypoglauca and E. tucumanensis)* and parpapatric *(E. hypoglauca and E. tucumanensis*) distributions showed little morphological and bioclimatic differentiation. Whether these species are a species complex (perhaps unlikely given that alleles of MYC and NIA for *E. illinita* seem to be unrelated to the alleles of the other species), a recent speciation event with little time for morphological/bioclimatic differentiation, or a case of allopatric speciation with niche conservatism (Wiens, 2004, Kozak and Wiens, 2006) remains to be determined by further sampling.

<u>Clade H.</u> Morphological data supported the hypothesis a species boundary between *E. myrtilloides* and *E. polifolia* (Fig. 11g; Table 4). Since the geographic range of *E. polifolia* is fully embedded within the range of *E. myrtilloides–*at a lower elevation (Appendix 3)–and there is a lack of genealogical exclusivity of MYC and NIA alleles for both species, it is possible that this species boundary emerged from a recent parapatric speciation event (Table 1).

<u>E. pulverulenta</u>. Molecular, morphological and bioclimatic data supported the hypothesis that *E. pulverulenta* is a distinct species (Figs. 11a, b). Although there was evidence to suggest that the morphological gap separating this species from the species in clades A and B could be explained as geographic differentiation within a single species (Appendix 4), this seems unlikely given that several evolutionary isolated species (from other clades) occur at intermediate geographic localities between *E. pulverulenta* and species from clades A and B (Fig. 1).

In short, my confrontation of empirical evidence against three operational species criteria to evaluate 35 hypotheses of species boundaries in the genus *Escallonia* revealed that 27 (70%) species were supported as independently evolving segments of population-level lineages (de Queiroz, 2005, 2007). Clearly, not all species differed concurrently in molecular, morphological and bioclimatic characters; rather some species were either genealogically exclusive, others morphologically distinct, others ecologically different, and others showed combinations of these properties. This is not surprising given the nature of species (Mishler and Donoghue, 1982; Baum, 1998; de Quieroz, 2005, 2007) and the timeframe within which *Escallonia* has likely diversified (Zapata, 2013). The weight of the evidence to reject the few hypotheses of species boundaries for which the data did not meet the operational species criteria was weak and likely compromised by lack of sampling. Creating taxonomic turmoil in the systematics of *Escallonia* by rejecting these hypotheses with weak evidence is premature at this point. Therefore, I prefer taxonomic stability and retain the current hypotheses of species boundaries as a useful framework to guide further sampling, and evaluate critically these hypotheses with thorough analyses in future studies. Interestingly, Rieseberg et al. (2006) reported that 70-75% of plant species represent biologically real entities. The results presented here fall within this range.

## ACKNOWLEDGEMENTS

I thank Peter Stevens, Elizabeth Kellogg, Kenneth Olsen, and Peter Hoch for comments on earlier versions of this manuscript. Jim Solomon and Andrea Voyer (MO) provided help requesting and processing the specimen loans I received from the following herbaria: CORD, CTES, E, F, GH, GOET, K, L, LIL, MO, NY, RB, REU, RSA, SP, UC, and US; thanks to the collections’ managers of those herbaria for granting access to their collections. Thanks to Iván Jiménez for insightful discussions on data analysis. Funding to conduct this research was provided by the National Science Foundation (OISE-0738118 to P. F. Stevens and F. Zapata), the Whitney R. Harris World Ecology Center, the Federated Garden Club of Missouri, the American Society of Plant Taxonomists, the Garden Club of America, Idea Wild, the University of Missouri–St. Louis, and the Missouri Botanical Garden.

## APPENDIX 1.

List of all specimens analyzed in this study (It will be available upon final submission at https://bitbucket.org/fzapata/Escallonia)

## APPENDIX 2.

Matrix of morphological measurements and bioclimatic variables (It will be available upon final submission at https://bitbucket.org/fzapata/Escallonia)

## APPENDIX 3.

Elevational ranges of species within clades. Each panel is surrounded by a colored box according to the color of the clade the species belongs to (for clade color, see Fig. 1). Each species is color coded (for species color, see Fig 5). a) clade A; b) clade B; c) clade D; d) clade F plus *E. virgata* and *E. gayana* (see text for details); e) clade G; f) clade H; g) *E. pulverulenta* plus clades A and B (see text for details).

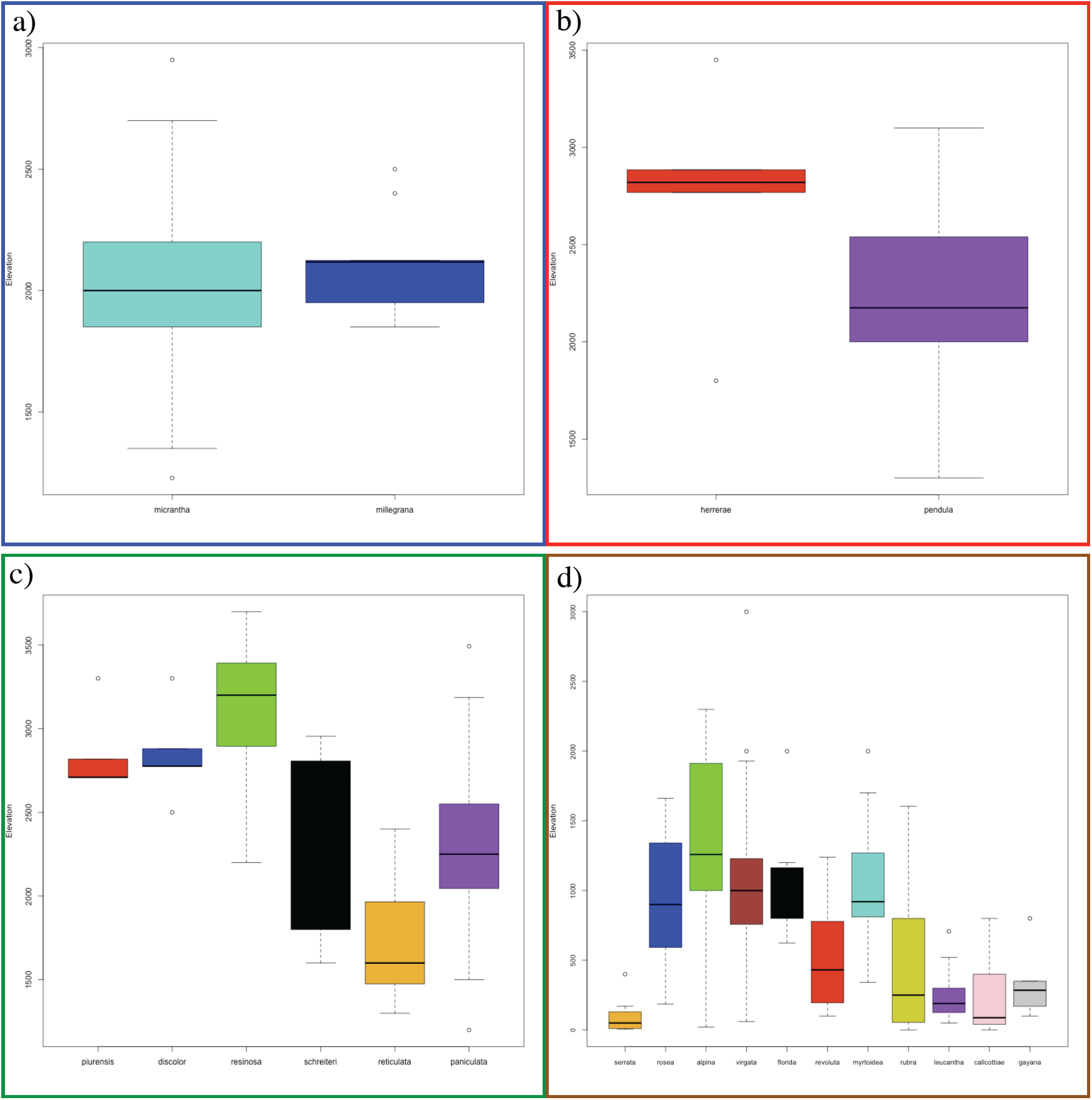
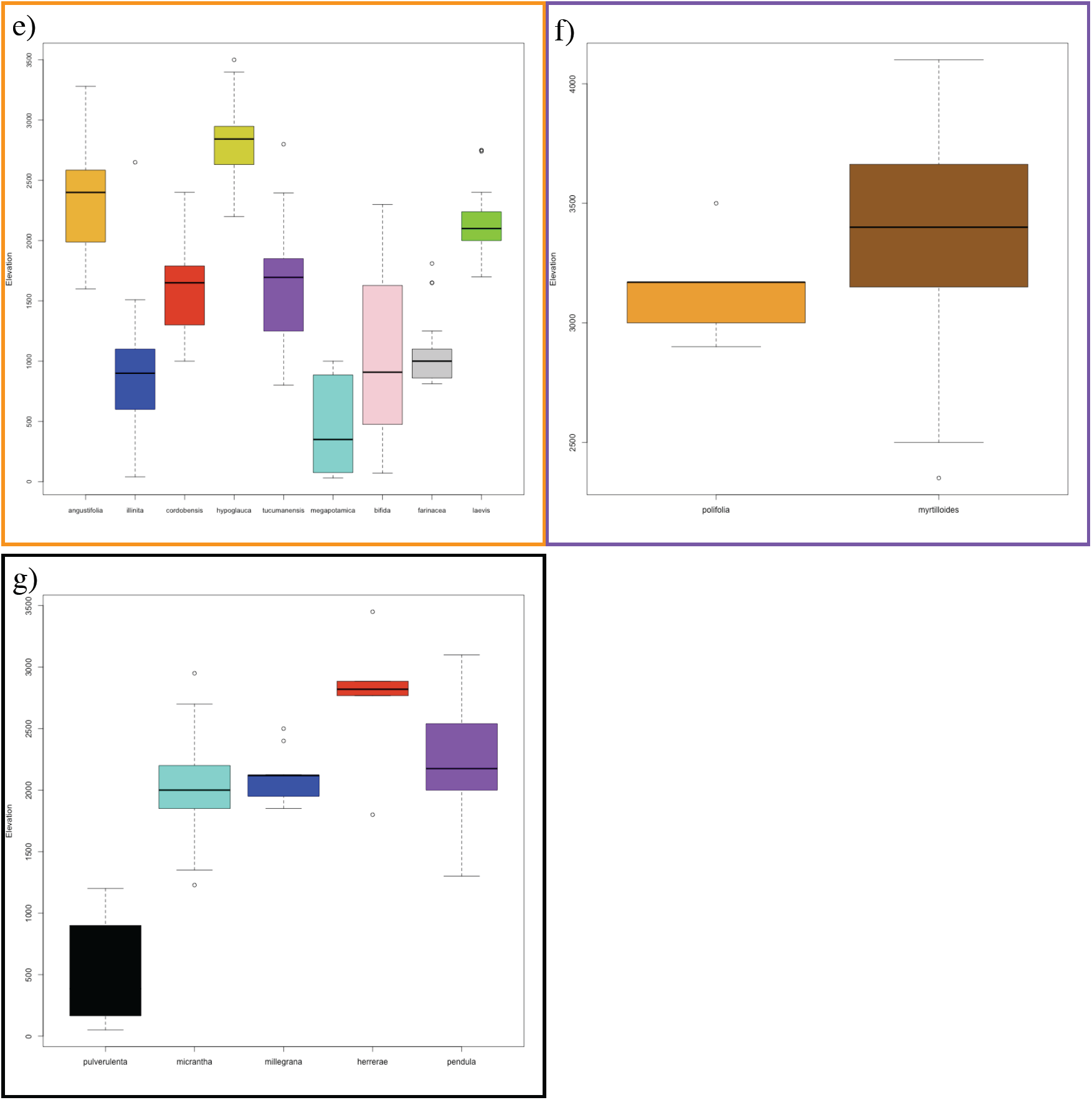

## APPENDIX 4.

Pairwise comparisons contrasting a model that does not require a species boundary to explain a morphological gap *vs*. a model that requires a species boundary (shaded box for each pairwise comparison). Comparisons that did not require detrending of the original response variable (see Zapata and Jiménez, 2010): clade F: *E. rose-E. virg, E. E. rubr.-E. gaya*, and *E. leuc.-E. gaya;* clade G: *E. bifi.-E. fari*. Significant regression (RDA) coefficients (*P* < 0.05) in bold. SE: spatial eigenvector; sp.: species boundary (i.e., matrix [0, 1]); sp*SE: interaction of species boundary with spatial eigenvector. For clade names, see Fig. 1. For details on statistical test, see Zapata and Jiménez (2012).

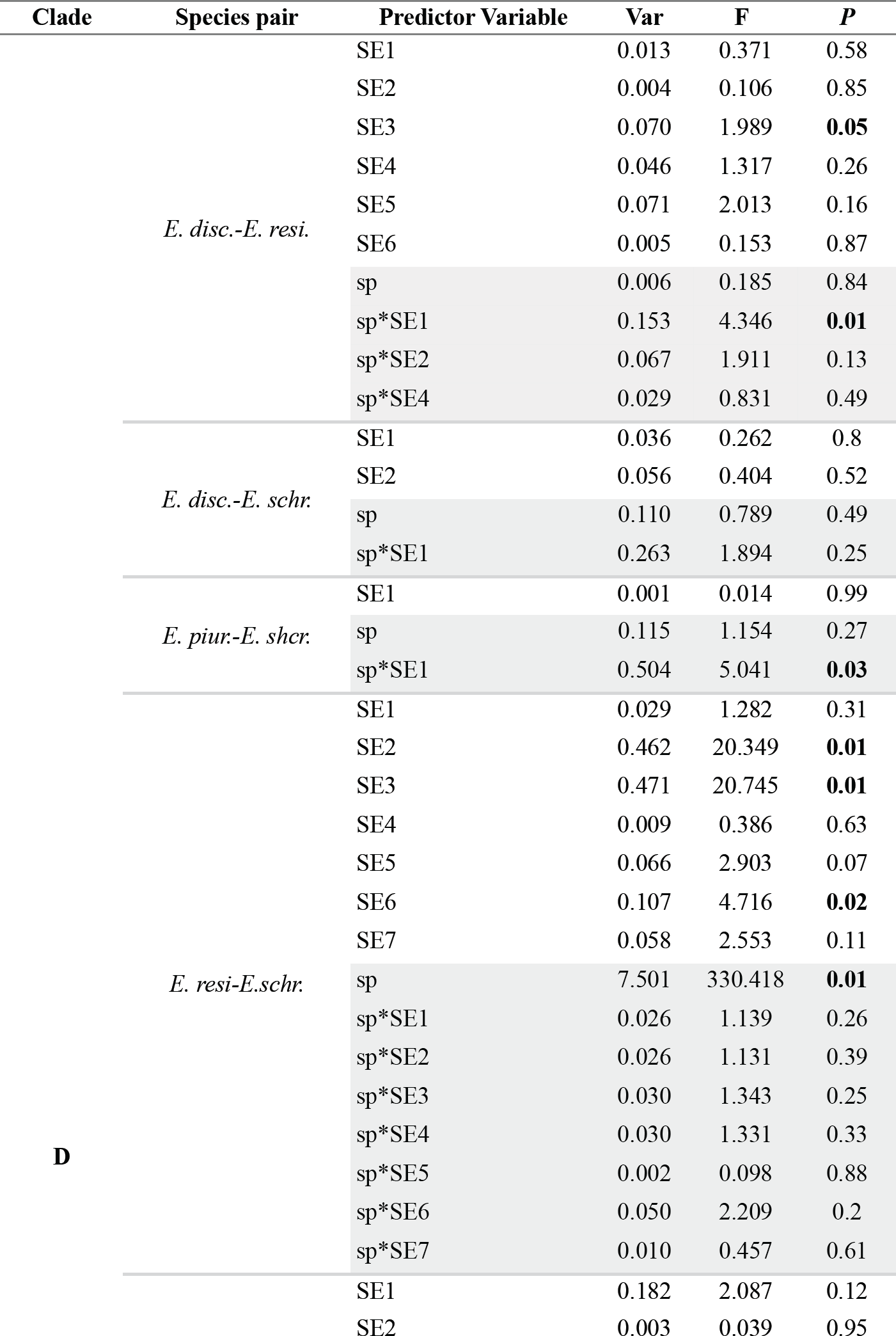
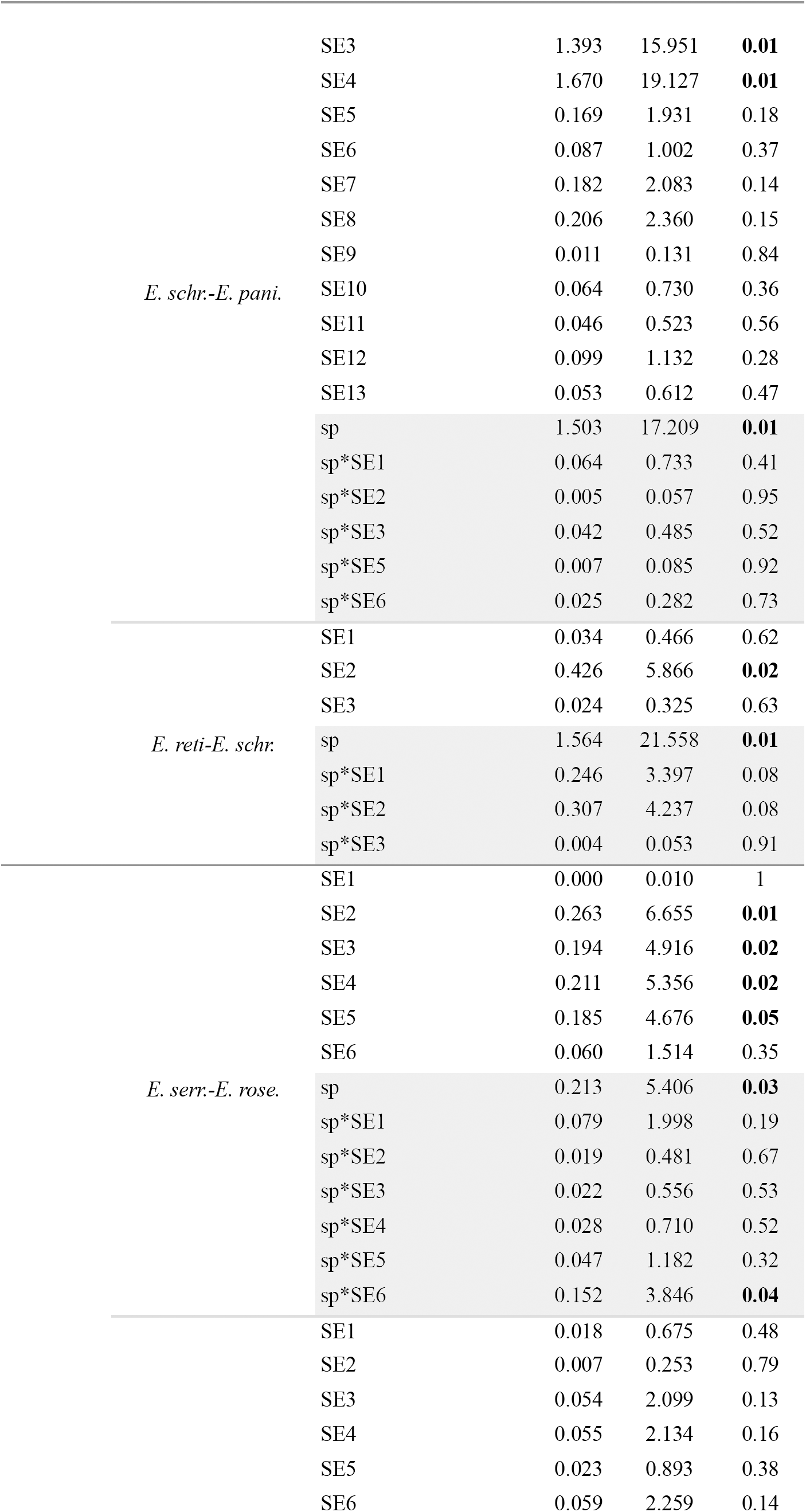
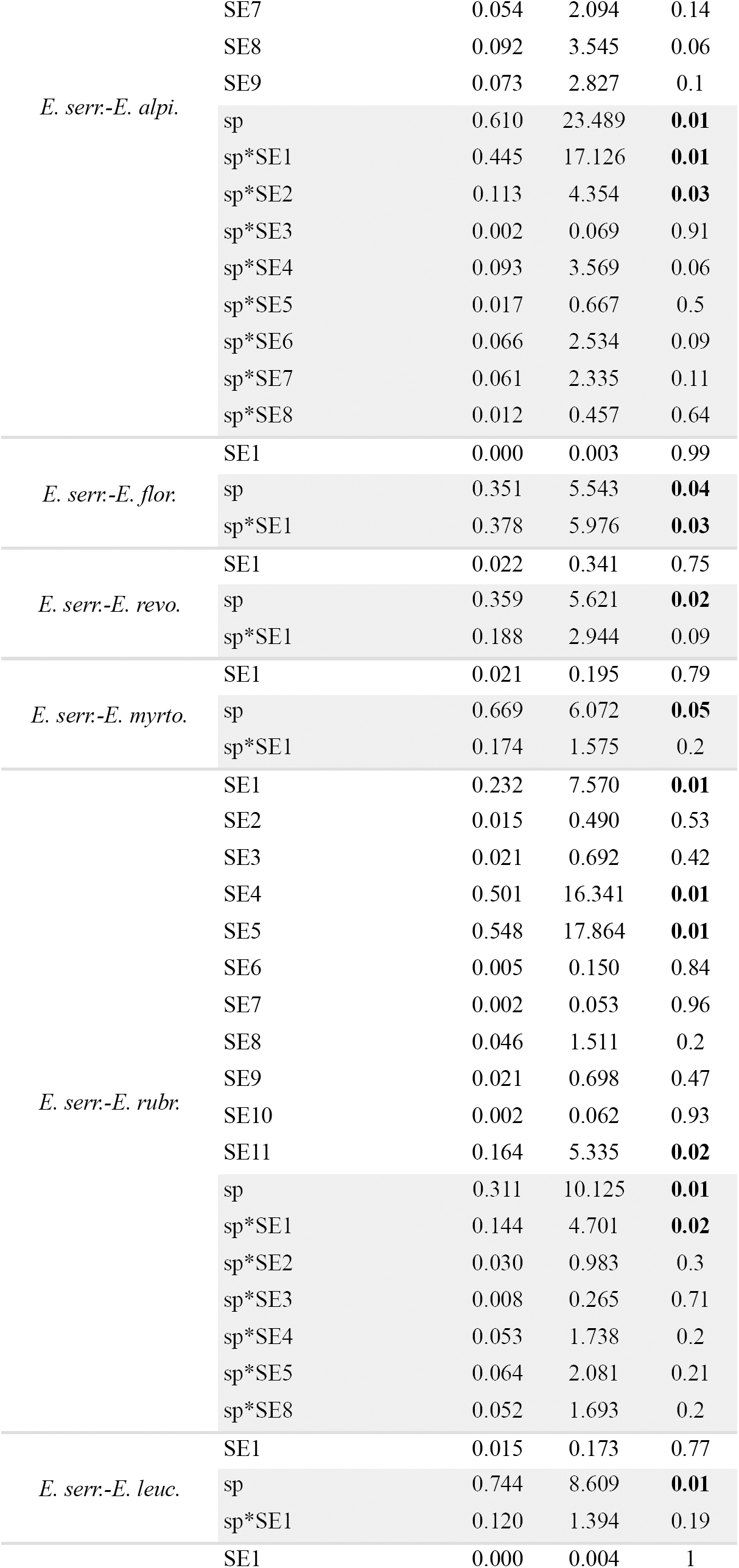
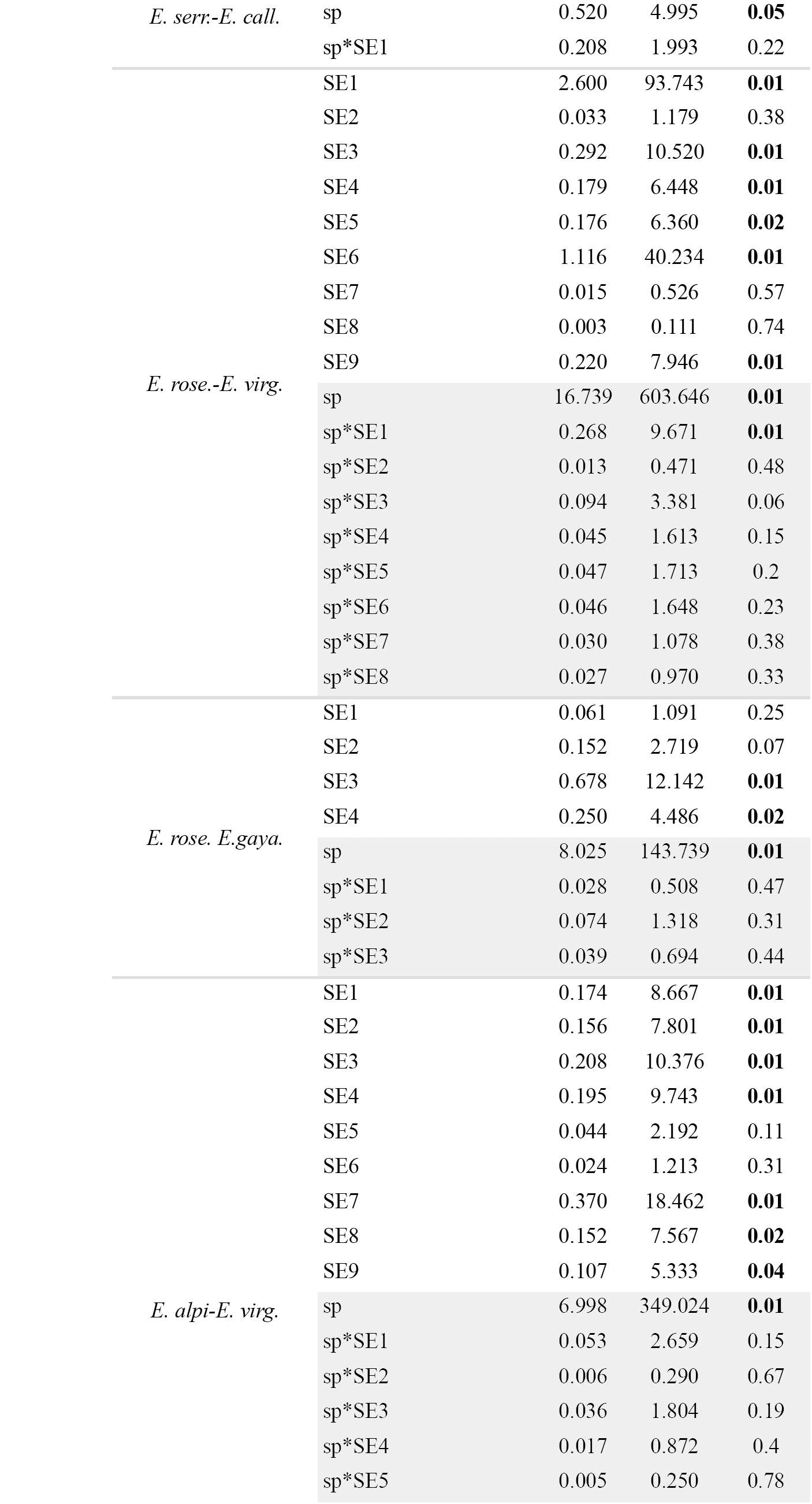
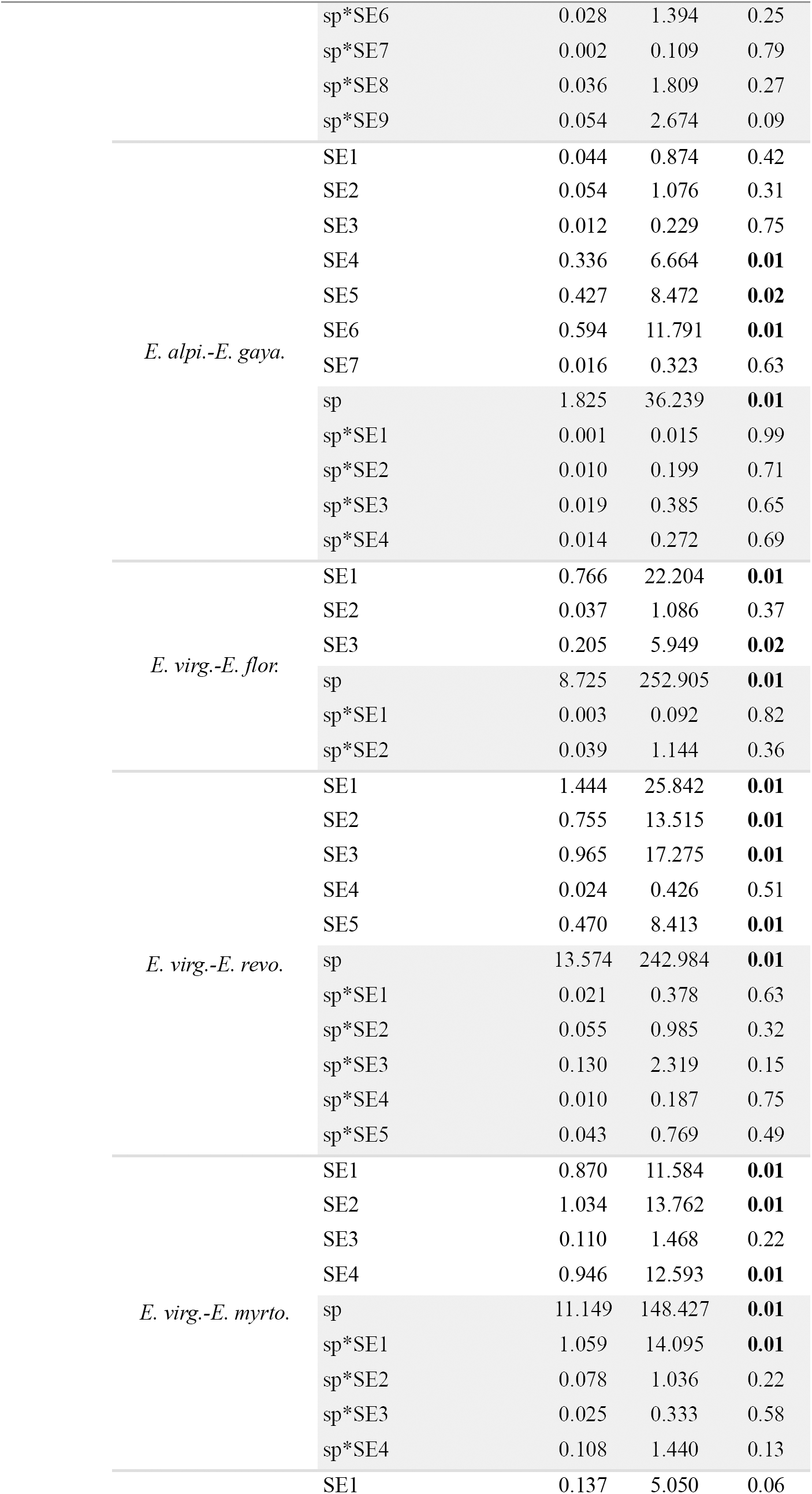
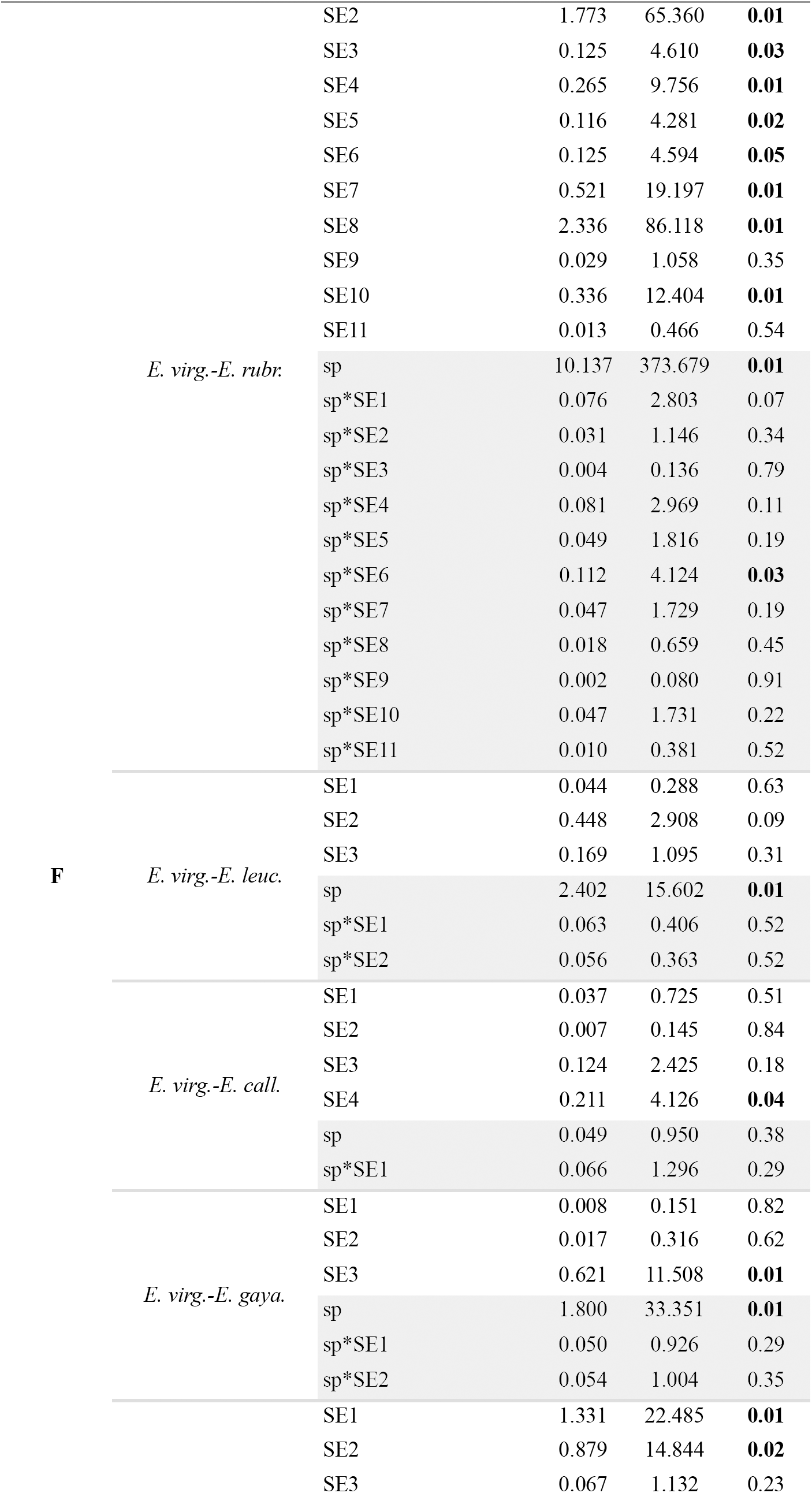
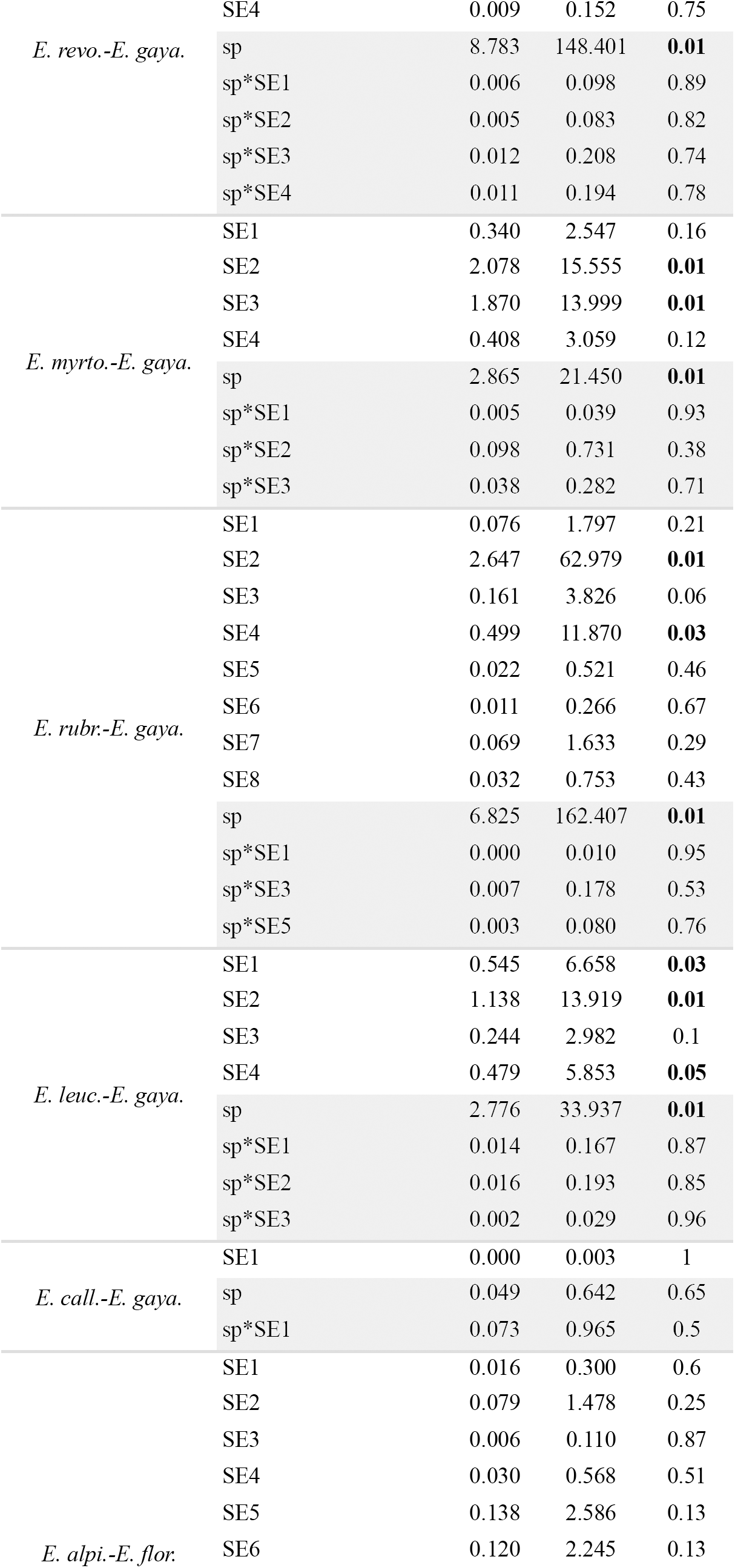
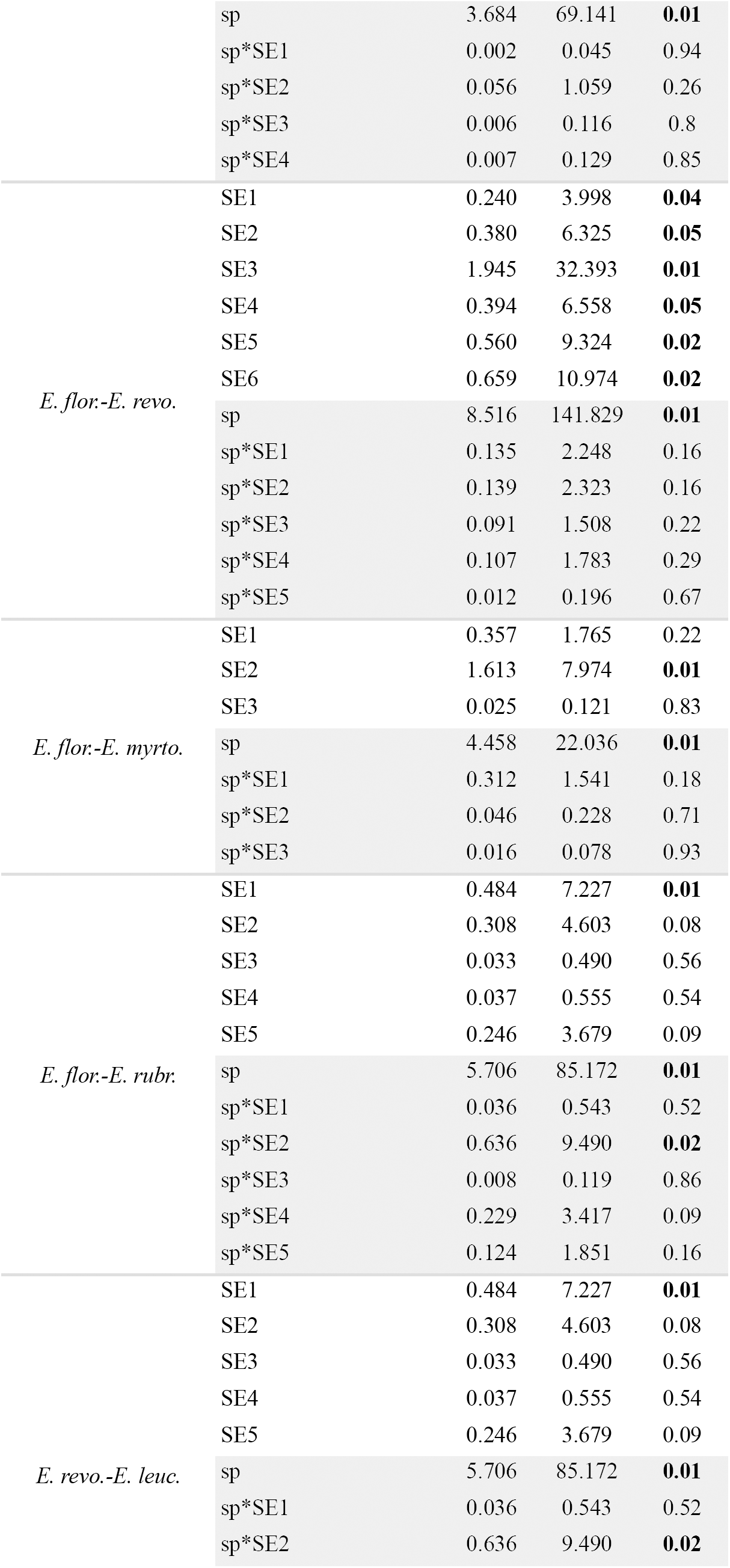
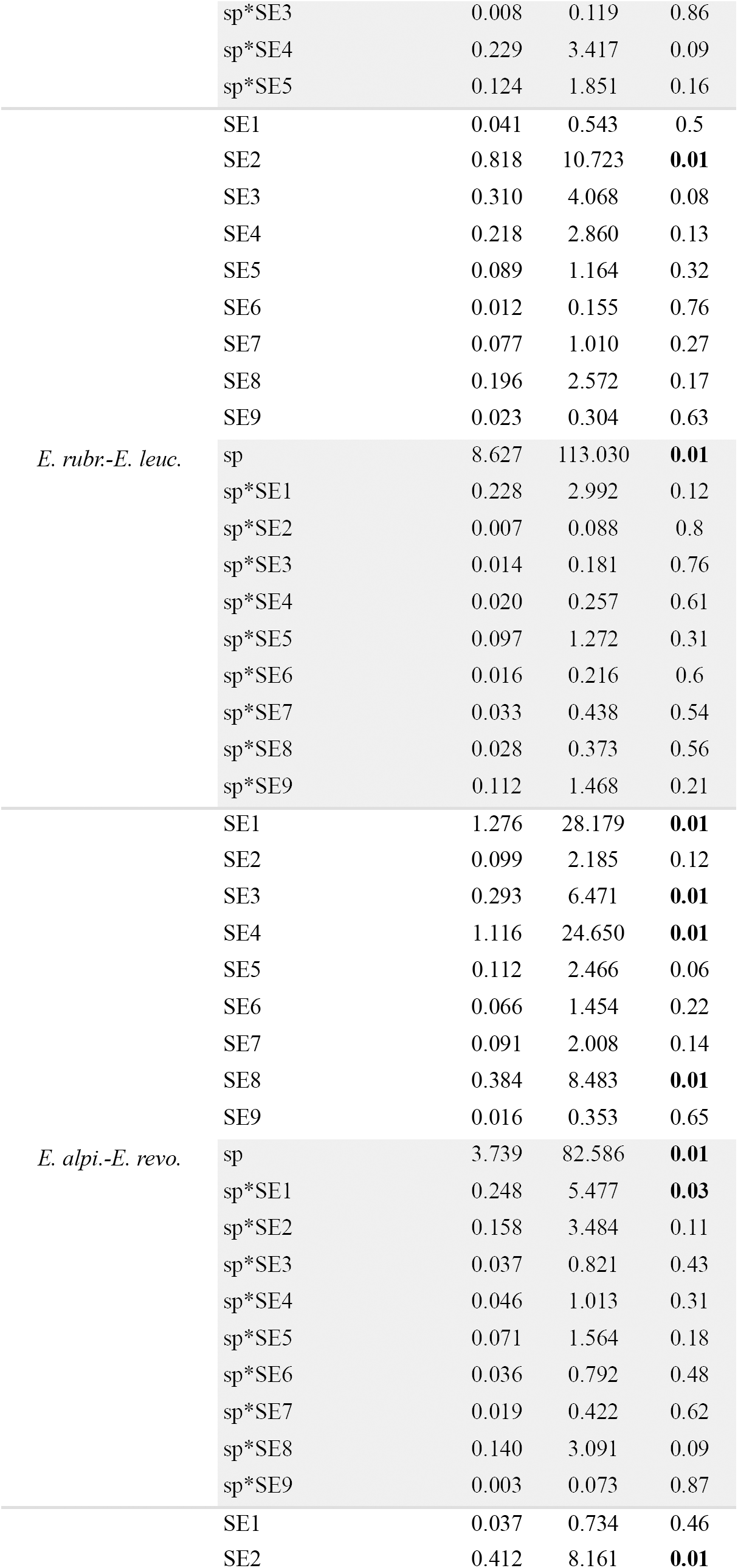
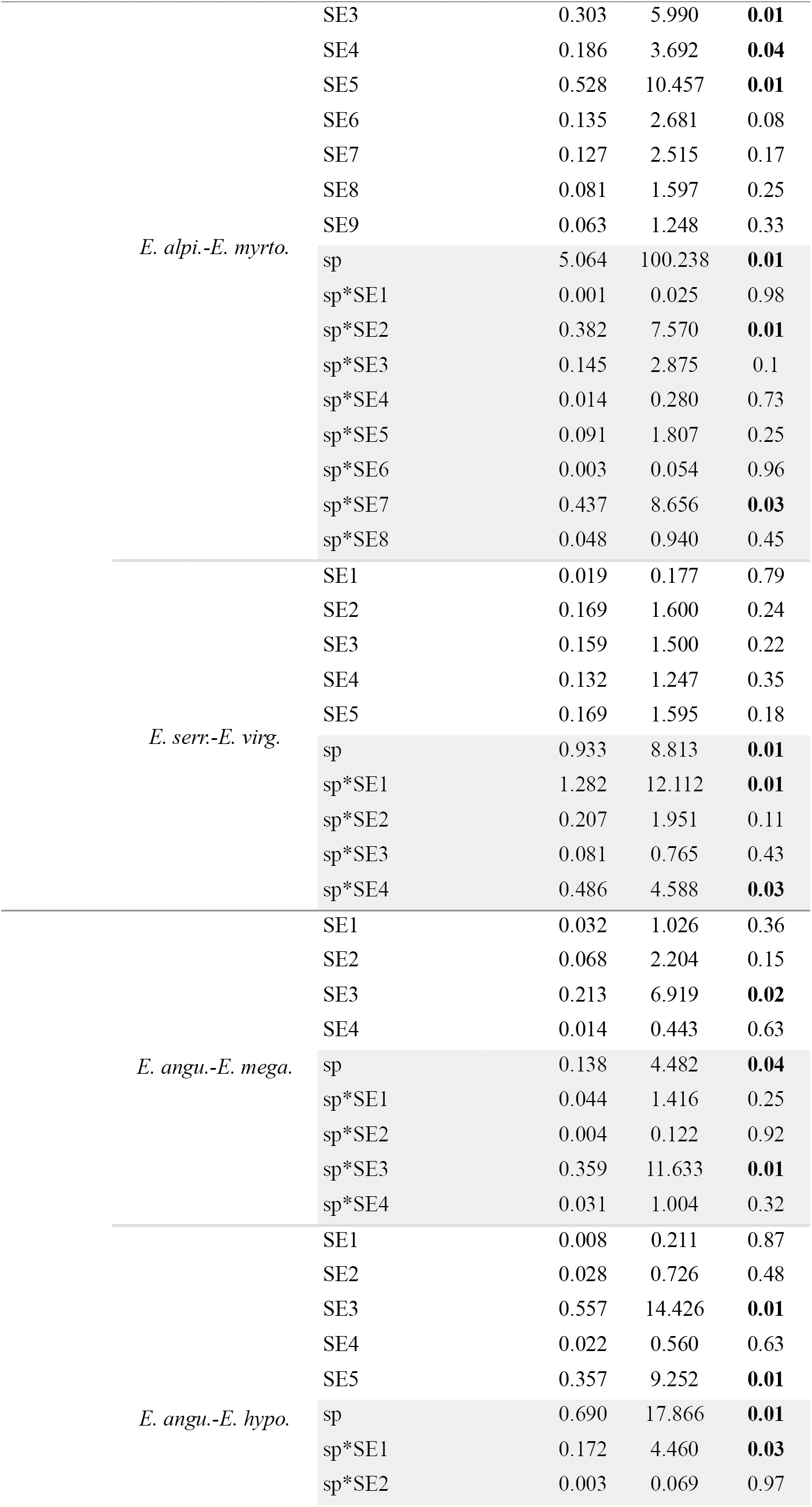
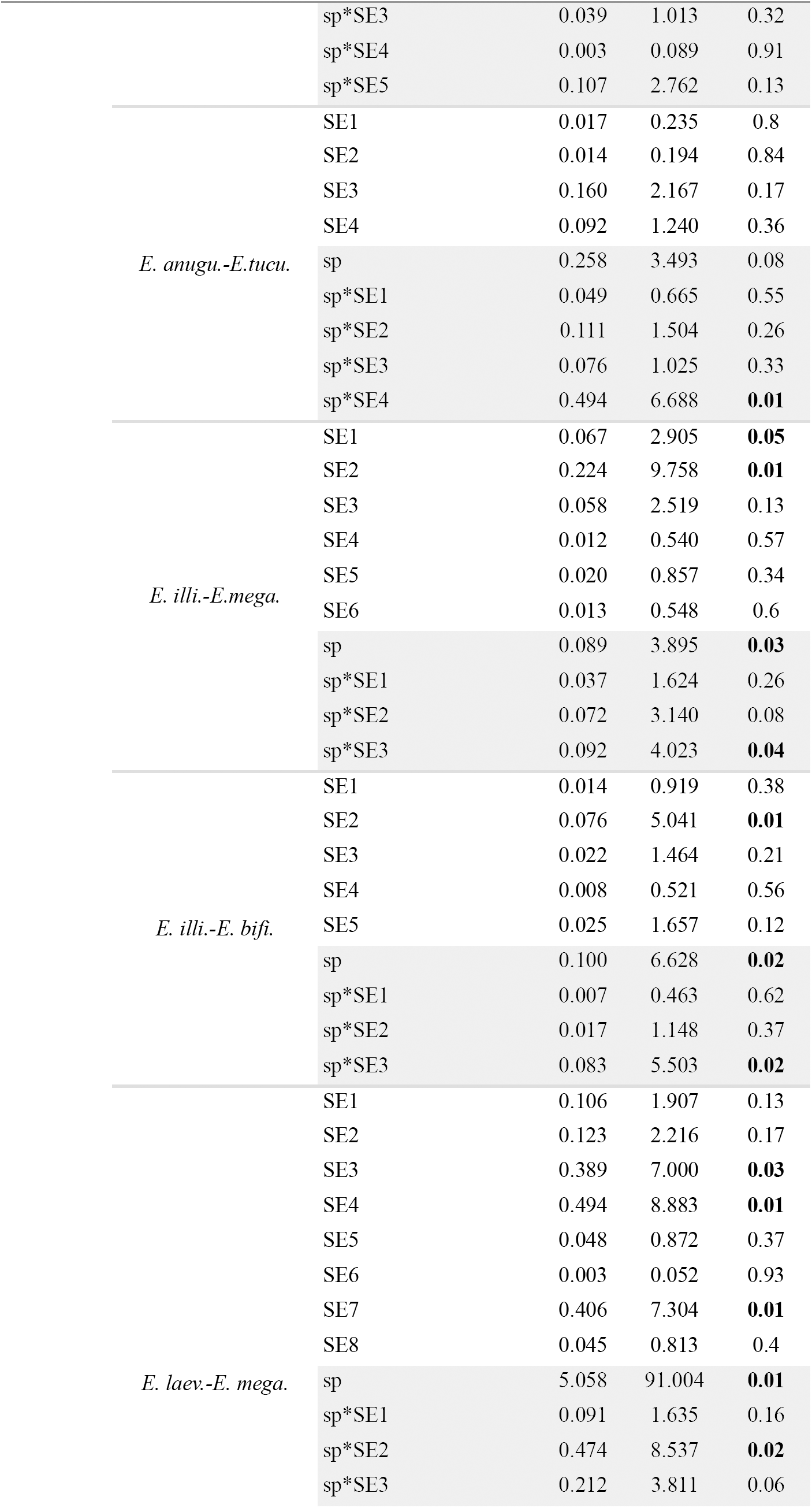
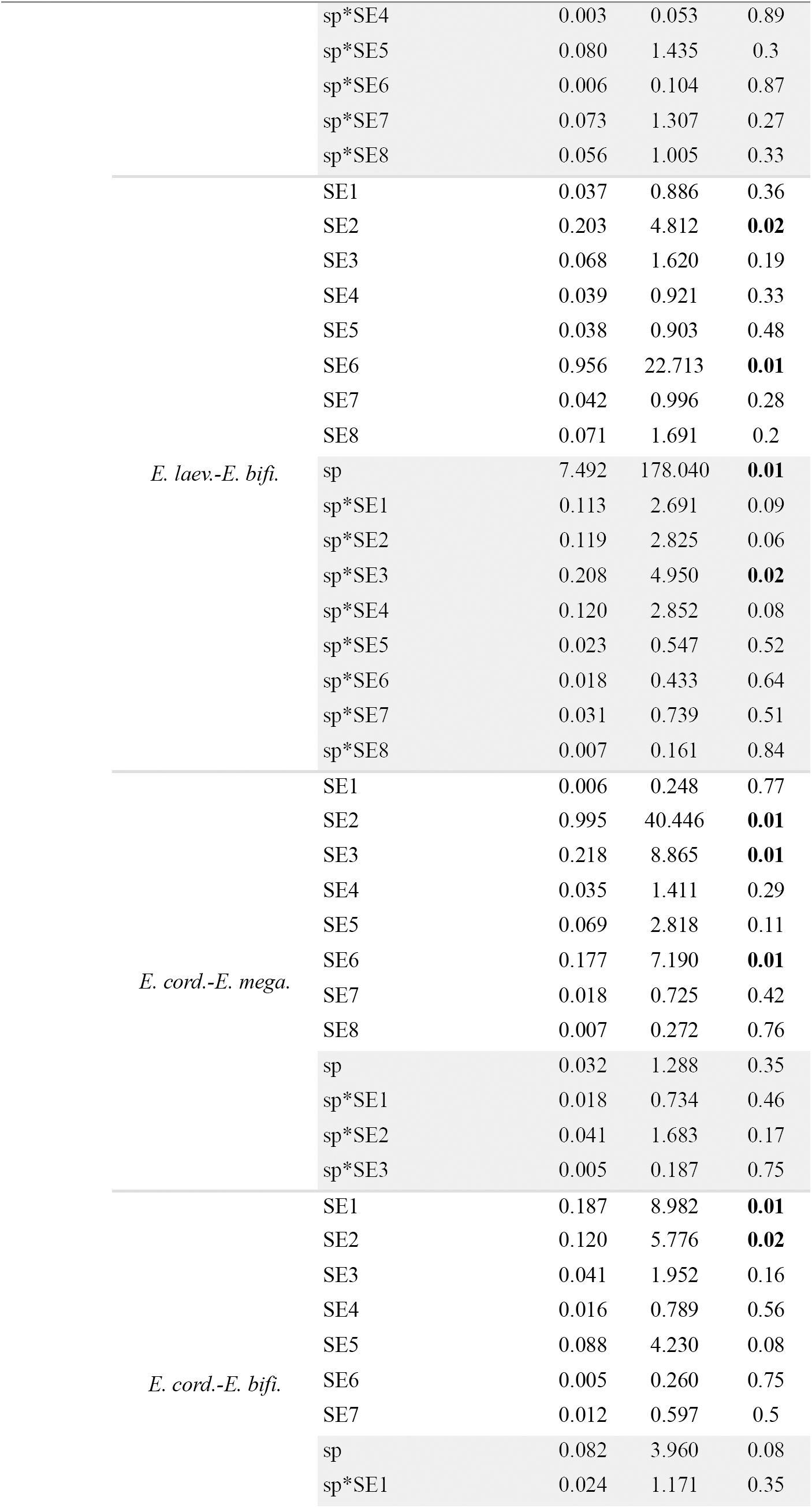
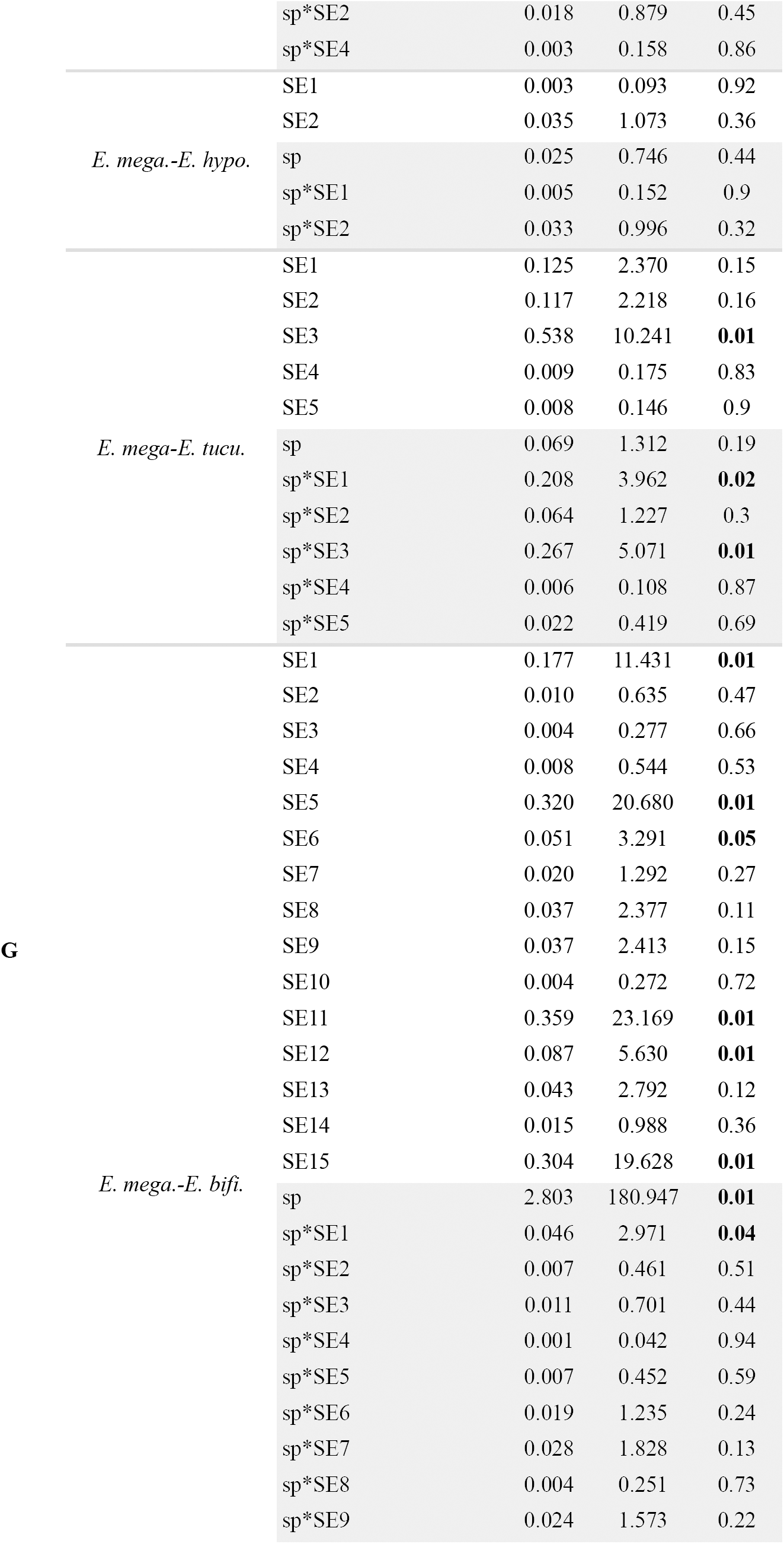
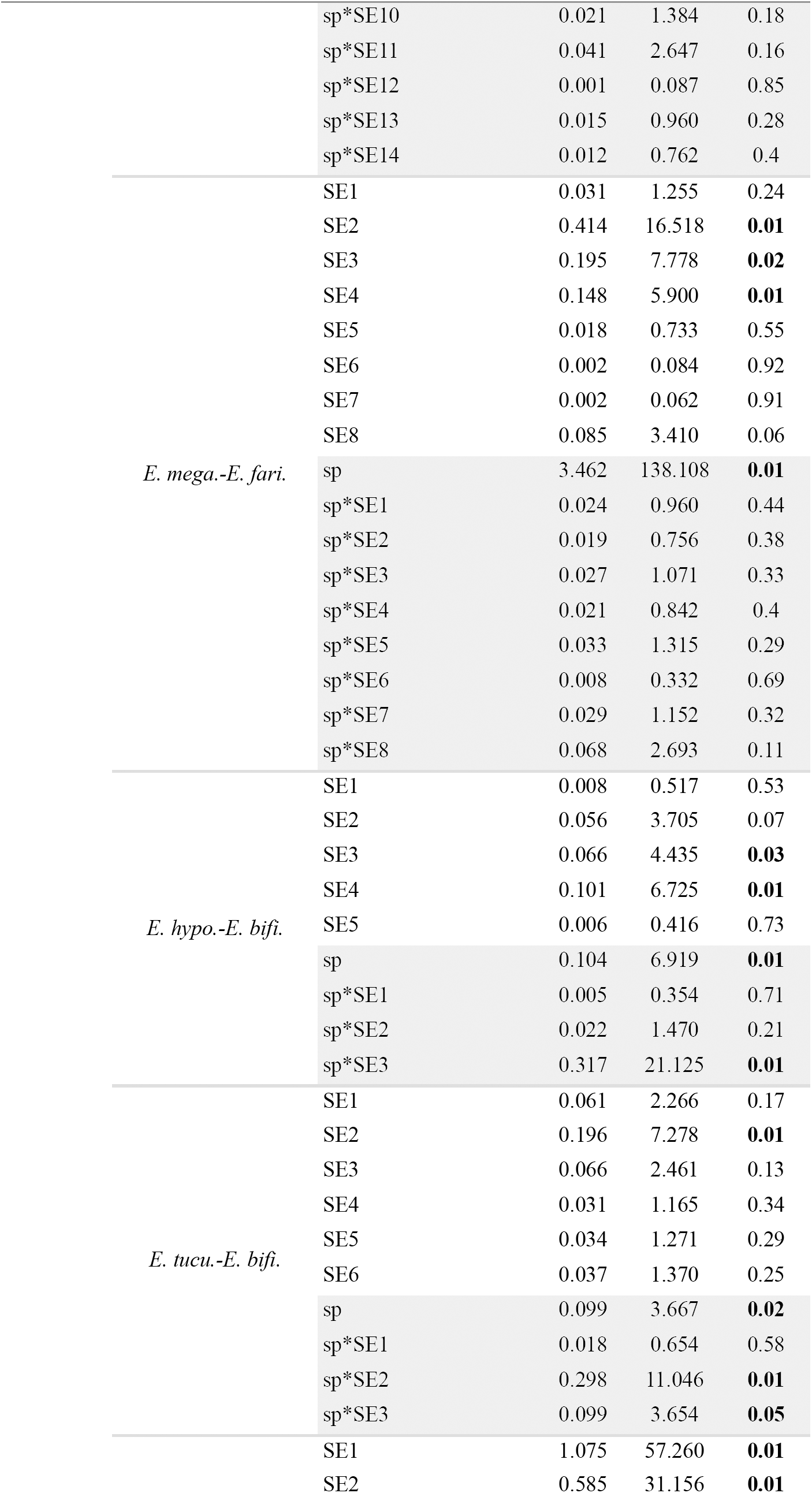
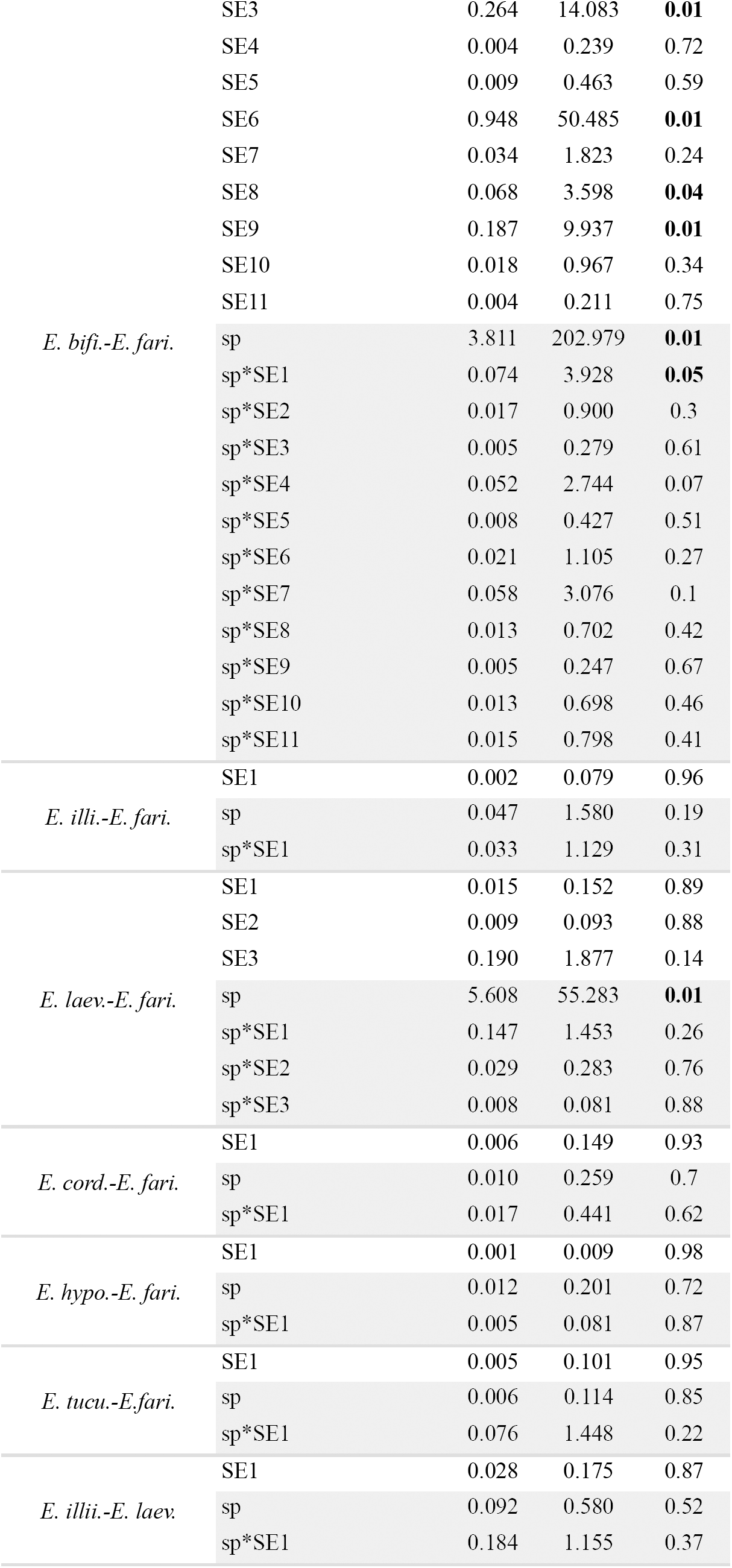
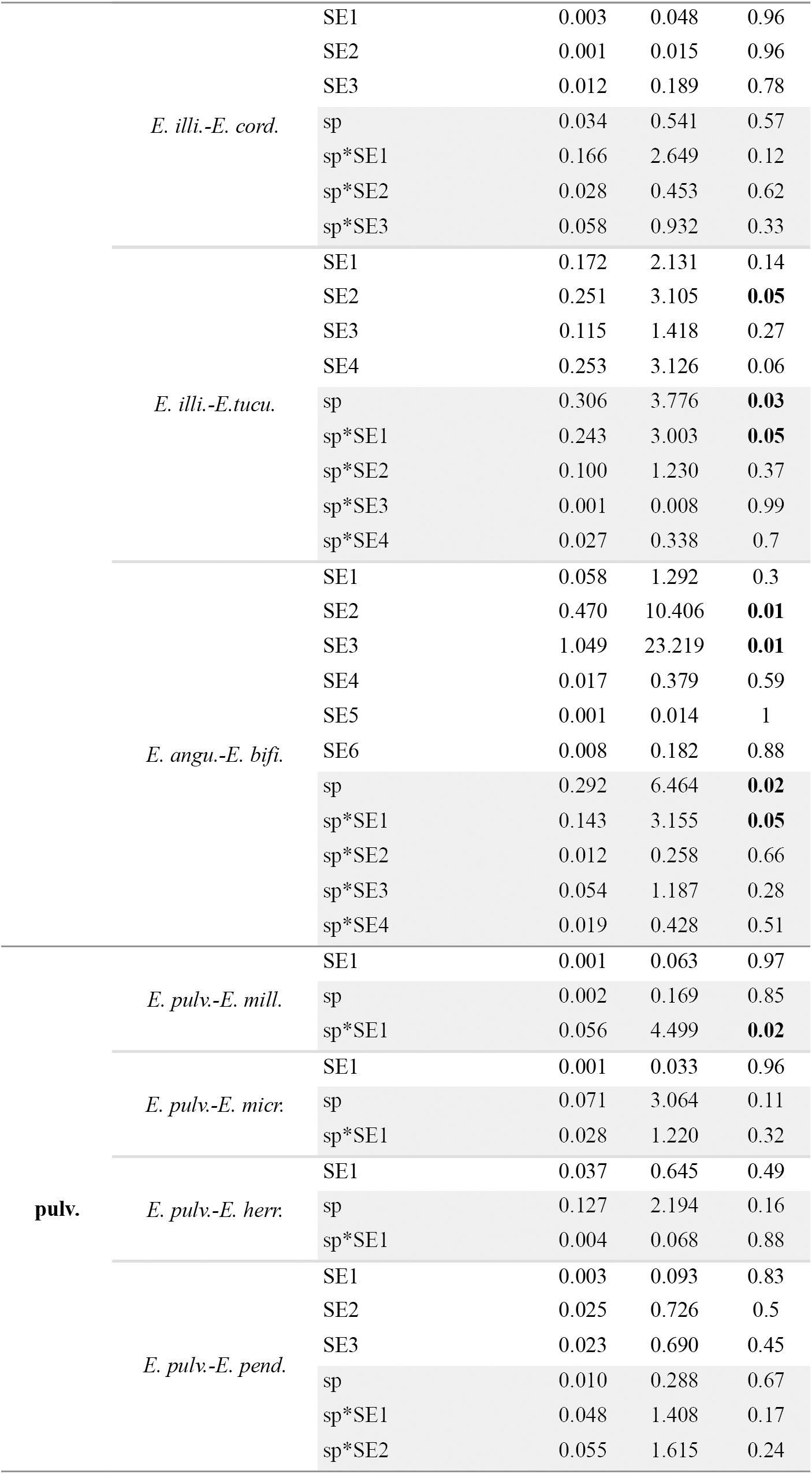

